# ConvexML: Fast and accurate branch length estimation under irreversible mutation models, illustrated through applications to CRISPR/Cas9-based lineage tracing

**DOI:** 10.1101/2023.12.03.569785

**Authors:** Sebastian Prillo, Akshay Ravoor, Nir Yosef, Yun S. Song

## Abstract

Branch length estimation is a fundamental problem in Statistical Phylogenetics and a core component of tree reconstruction algorithms. Traditionally, general time-reversible mutation models are employed, and many software tools exist for this scenario. With the advent of CRISPR/Cas9 lineage tracing technologies, there has been significant interest in the study of branch length estimation under *irreversible* mutation models. Under the CRISPR/Cas9 mutation model, irreversible mutations – in the form of DNA insertions or deletions – are accrued during the experiment, which are then read out at the single-cell level to reconstruct the cell lineage tree. However, most of the analyses of CRISPR/Cas9 lineage tracing data have so far been limited to the reconstruction of single-cell tree *topologies*, which depict lineage relationships between cells, but not the amount of time that has passed between ancestral cell states and the present. Time-resolved trees, known as *chronograms*, would allow one to study the evolutionary dynamics of cell populations at an unprecedented level of resolution. Indeed, time-resolved trees would reveal the timing of events on the tree, the relative fitness of subclones, and the dynamics underlying phenotypic changes in the cell population – among other important applications. In this work, we introduce the first scalable and accurate method to refine any given single-cell tree topology into a single-cell chronogram by estimating its branch lengths. To do this, we perform regularized maximum likelihood estimation under a general irreversible mutation model, paired with a conservative version of maximum parsimony that reconstructs only the ancestral states that we are confident about. To deal with the particularities of CRISPR/Cas9 lineage tracing data – such as *double-resection* events which affect runs of consecutive sites – we avoid making our model more complex and instead opt for using a simple but effective data encoding scheme. Similarly, we avoid explicitly modeling the missing data mechanisms – such as heritable missing data – by instead assuming that they are missing completely at random. We stabilize estimates in low-information regimes by using a simple penalized version of maximum likelihood estimation (MLE) using a minimum branch length constraint and pseudocounts. All this leads to a convex MLE problem that can be readily solved in seconds with off-the-shelf convex optimization solvers. We benchmark our method using both simulations and real lineage tracing data, and show that it performs well on several tasks, matching or outperforming competing methods such as TiDeTree and LAML in terms of accuracy, while being 10 ~ 100 × faster. Notably, our statistical model is simpler and more general, as we do not explicitly model the intricacies of CRISPR/Cas9 lineage tracing data. In this sense, our contribution is twofold: (1) a fast and robust method for branch length estimation under a general irreversible mutation model, and (2) a data encoding scheme specific to CRISPR/Cas9-lineage tracing data which makes it amenable to the general model. Our branch length estimation method, which we call ‘ConvexML’, should be broadly applicable to any evolutionary model with irreversible mutations (ideally, with high diversity) and an approximately ignorable missing data mechanism. ‘ConvexML’ is available through the `convexml` open source Python package.

## Introduction

Many important biological processes such as development, cancer progression and adaptive immunity unfold through time, originating from a small progenitor cell population and progressing through repeated cell division. A realization of these processes can be described by a single-cell *chronogram*: a rooted tree that represents the history of a clone, where each edge represents the lifetime of a cell, and internal nodes represent cell division events, as depicted in Figure 1a. Single-cell chronograms thus capture the entire developmental history of the cell population, allowing us to understand when and how cells commit to their fates. Because of this, single-cell chronograms have been of interest for decades.

**Fig. 1.**
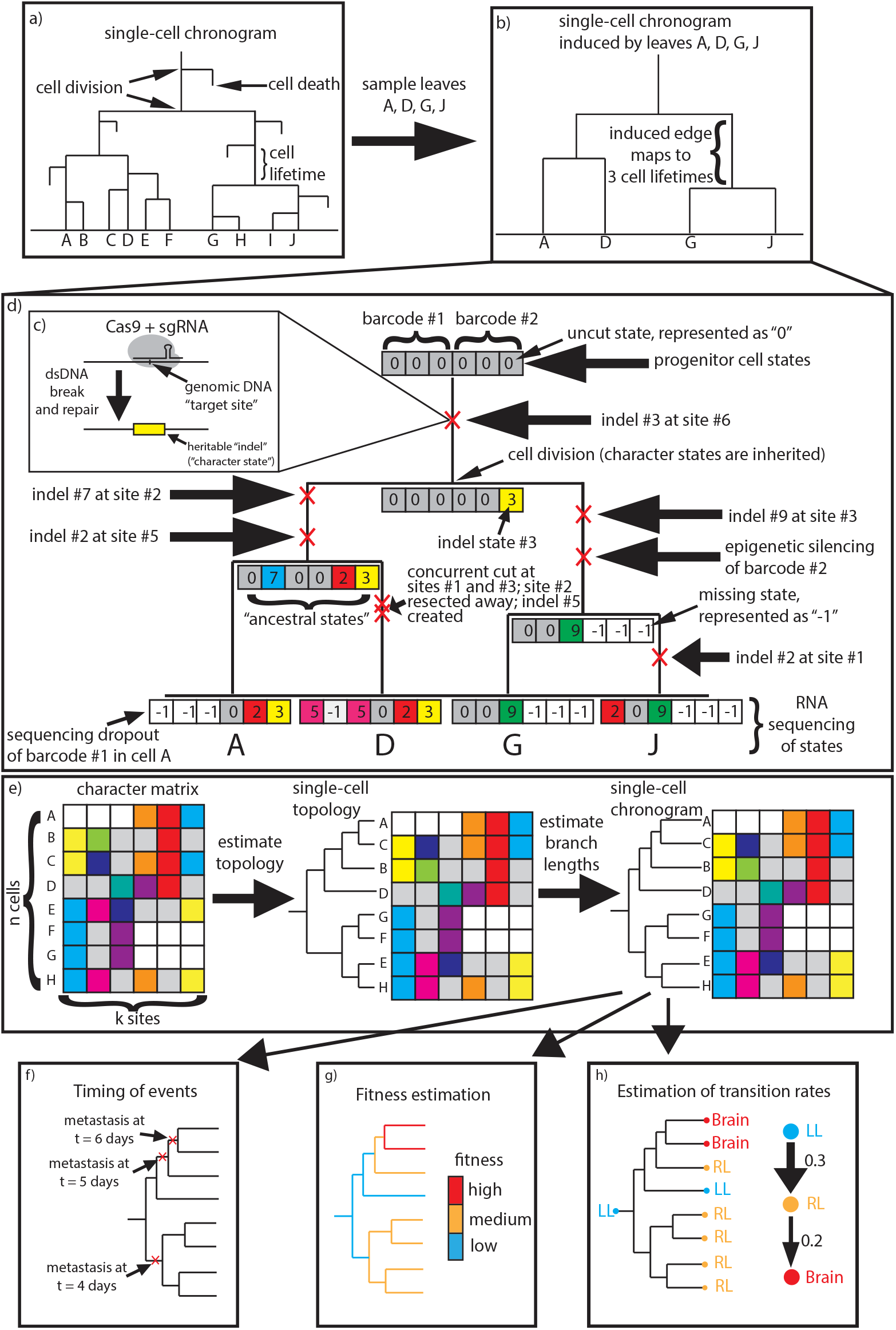
a) A single-cell chronogram over 10 cells A-J. b) The single-cell chronogram induced by leaves A, D, G, J. c) CRISPR/Cas9 binds and cuts a target site, introducing an indel at that site. d) The CRISPR/Cas9 lineage tracing system recording the lineage history of a cell population. We represent the uncut state with ‘0’, distinct indels with positive integers, and missing data with ‘−1’. Note specifically our choice of representation of double-resection events. e) Our approach to estimating single-cell chronograms from CRISPR/Cas9 lineage tracing data: A single-cell topology is first estimated using any of many available methods, such as those provided by the `cassiopeia` package (Jones et al., 2020) or otherwise (Gong et al., 2021). Next, our novel branch length estimator is applied to estimate branch lengths for that topology. (Note that the data here are unrelated to that of panel d)). f)-h) Single-cell chronograms reveal the timing of events on the tree, single-cell fitness scores, and phenotypic transition rates, among other important applications.

Over 40 years ago, the first single-cell chronogram for the development of *C. elegans* was determined through visual observation over the timespan from zygote to hatched larva (Sulston et al., 1983; Deppe et al., 1978). Since then, the advent of CRISPR/Cas9 genome editing and high-throughput single-cell sequencing technologies has enabled lineage tracing for thousands of cells *in vivo* (McKenna et al., 2016; Raj et al., 2018; Spanjaard et al., 2018; Wagner et al., 2018; Kalhor et al., 2018; Chan et al., 2019). However, unlike microscopy-based techniques, these approaches record lineage history indirectly through irreversible heritable Cas9 mutations – called *indels* – at engineered DNA *target sites*, as depicted in Figure 1c. These target sites (or simply *sites*) are arranged into lineage tracing *barcodes* (or *character arrays*) that act as fake genes which mutate stochastically and are transcribed and measured through the transcriptome. Figure 1d depicts the evolution and inheritance of CRISPR/Cas9 lineage tracing barcodes on a single-cell chronogram. Inferring a lineage tree from this CRISPR/Cas9 lineage tracing data is challenging, and has led to the development of many computational methods (Gong et al., 2021).

Single-cell chronograms provide a detailed account of the cell population’s history, revealing the timing of events on the tree, the relative fitness of subclones, and the dynamics underlying phenotypic changes in the cell population – among others, as depicted in Figure 1f-h. This is in contrast to single-cell tree *topologies*, which are like single-cell chronograms except that they do not have branch lengths. Although single-cell tree topologies reveal the lineage relationships between cells, the lack of time resolution precludes them from tackling the aforementioned tasks.

Unfortunately, the task of estimating single-cell chronograms from CRISPR/Cas9 lineage tracing data is daunting, and methods developed so far have abandoned the hope of estimating single-cell chronograms to their full resolution. For one, only extant cells are sequenced, so it is not possible to pinpoint cell death events. Also, only a fraction of the cells in the population are sampled and sequenced, so one can only expect to estimate the single-cell chronogram *induced* by the sampled cells. Formally, the induced chronogram is defined as the subtree whose leaves are the sampled cells. For example, Figure 1b illustrates the induced chronogram obtained by sampling cells A, D, G, J in the chronogram from Figure 1a. Note that edges in the induced chronogram no longer map one-to-one to the lifetime of a cell; instead, they map one (edge)-to-many (cell lifetimes). Sampling cells greatly affects the distribution of branch lengths in the tree, as shown in Supplementary Figure 1. Most methods developed so far have not attempted to estimate branch lengths, opting instead to estimate just topologies (Gong et al., 2021), limiting their value.

The task of estimating single-cell chronograms is complicated by the fact that lineage tracing data are especially prone to go missing. There are three primary reasons for this. The first one is *sequencing dropouts*, whereby the limited capture efficiency of single-cell RNA-sequencing technologies leads to some barcodes not being sequenced. The second reason is heritable *epigenetic silencing* events. When this occurs, chromatin state is modified in such a way that a lineage tracing barcode is no longer transcribed and thus cannot be read out through the transcriptome. The third reason is *double-resection* events, wherein concurrent CRISPR/Cas9 cuts at proximal barcoding sites cause the flanked sites to be lost. Formally, in a double-resection event, two sites *i* and *j* in the same barcode are cut close in time, resulting in the whole segment of sites between *i* and *j* to disappear, and a shared indel to be created spanning all sites *i* through *j*. Missing data from epigenetic silencing and double-resection events is inherited upon cell division. Double-resection events are particularly challenging to represent because they create an indel that spans multiple barcoding sites and introduce complex correlations between barcoding sites which complicate branch length estimation (Yang, 1995). These three sources of missing data are illustrated in Figure 1d; note specifically our choice of representation of double-resection events, which will turn out to be crucial for branch length estimation.

The estimation of time-resolved trees from molecular data has a rich history outside of single-cell lineage tracing. The field of statistical phylogenetics has studied the problem extensively, with the goal of estimating gene trees and species trees from multiple sequence alignments of DNA and amino-acid sequences. Popular software for reconstructing phylogenetic trees includes FastTree (Price et al., 2010), PhyML (Guindon et al., 2010), IQ-Tree (Minh et al., 2020) RAxML (Stamatakis, 2014), and BEAST2 (Bouckaert et al., 2019). Unfortunately, these methods are not suitable for CRISPR/Cas9 lineage tracing data because (1) they were designed for small, finite state spaces such as the 20 amino acid alphabet, (2) they require a known substitution model which is usually assumed to be reversible, or (3) they have an elevated computational cost characteristic of Monte-Carlo Bayesian methods, forbidding them from scaling beyond a few hundred sequences. CRISPR/Cas9 lineage tracing datasets contain hundreds of unique states, an unknown substitution model which is irreversible, and hundreds to thousands of cells.

Although attempts have been made to adapt Statistical Phylogenetics models to CRISPR/Cas9 lineage tracing data, they still suffer from elevated computational costs. The recent work TiDeTree (Seidel and Stadler, 2022) implemented within the BEAST2 platform (Bouckaert et al., 2019) takes several hours to infer a chronogram for a tree with just 700 leaves. This comes from the need to run MCMC chains for inference. Similarly, the GAPML method (Feng et al., 2021) which was designed specifically for the GESTALT technology takes up to 2 hours on a tree with just 200 leaves. This comes from modeling the correlations between sites caused by double-resection events, and the cost of marginalizing out the character states of the ancestral (unobserved) cells - which we shall call the *ancestral states* for short. The LAML method (Mai et al., 2024), which is based on maximum likelihood estimation rather than Bayesian inference, also marginalizes out the ancestral states, leading to increased computational complexity. This calls for new methods that can improve the trade-off between computational efficiency and statistical efficiency.

In this work, we introduce the first scalable and accurate method to estimate single-cell chronograms from CRISPR/Cas9 lineage tracing data. Our approach is modular: we propose first estimating a single-cell topology using any of many available methods (Jones et al., 2020; Gong et al., 2021), and then refining it into a single-cell chronogram by estimating its branch lengths, as depicted in Figure 1e. To estimate branch lengths, we leverage a quite general irreversible mutation model, and use a conservative version of maximum parsimony to reconstruct most – but not all – of the ancestral states, while still producing essentially unbiased estimates. We pay particular attention to CRISPR/Cas9 double-resection events, and propose a novel encoding and treatment that are well-suited to our model. Importantly, we do not explicitly model double-resection events – instead, we encode them in a way that the model observes two independent mutations at the flanking sites; this is in stark contrast to prior methods such as GAPML (Feng et al., 2021) which explicitly model double-resection events. We also do not explicitly model heritable missing data or sequencing dropouts, opting for statistical ignorability instead, which is in stark contrast to the recently proposed LAML method (Mai et al., 2024). Because lineage tracing data can be of varying quality, we propose the use of a penalized version of maximum likelihood estimation (MLE) using a minimum branch length constraint and pseudocounts. Taken together, all this leads to a straightforward convex MLE problem that can be readily solved with off-the-shelf convex optimization solvers. Our method typically takes only a few seconds to estimate branch lengths for a tree topology with 400 leaves using a single CPU core. Our method, which we call ‘ConvexML’, is simple and general enough that it should be broadly applicable to other irreversible mutation models with a large enough state space. In particular, our pseudocount regularization approach may be of broader interest.

We develop a benchmarking suite with three tasks to asses the performance of single-cell chronogram estimation methods: (1) estimation of the times of the internal nodes in the tree, (2) estimation of the number of ancestral lineages halfway (time-wise) through the experiment, and (3) fitness estimation. We show that our ConvexML-method performs well on these tasks, outperforming more naive baselines as well as the recently proposed LAML method (Mai et al., 2024). We also compare our method’s performance against that of an ‘oracle’ model that has access to the ground truth ancestral states and show that it has comparable performance. In contrast, naively employing maximum parsimony leads to biased branch length estimates. This validates the suitability of conservative maximum parsimony for CRISPR/Cas9 lineage tracing data, a key methodological innovation of our work which underlies our scalability. Similarly, we show the suitability of our novel representation of missing data, specifically that concerning double-resection events. We further benchmark our method on a real lineage tracing dataset (intMEMOIR (Chow et al., 2021)), where we show that our method matches or outperforms both LAML and TiDeTree while being significantly faster. To finish, we discuss some of the many extensions of our method that are made possible by its simplicity, such as branch length estimation for non-ultrametric trees. We name our method ‘ConvexML’ and make it available through the convexml open source Python package.

## Materials and Methods

### A Statistical Model for CRISPR/Cas9 Lineage Tracing Data

We start by describing how the CRISPR/Cas9 lineage tracing data are represented (or *encoded*). Let *n* be the number of cells assayed and let *k* be the number of lineage tracing sites or characters. The observed lineage tracing data – called the *character matrix* – is represented as a matrix *X* of size *n* × *k* with integer entries. The *i*-th row of *X* represents the observed character states of the *i*-th cell. The character states are grouped consecutively into *barcodes* of a known fixed size. Each non-missing entry of the character matrix represents an indel - an insertion or deletion of nucleotides at the given site. Indels are represented with distinct positive integers, and 0 is used to represent the uncut state. The integer −1 is used to represent missing data. Figure 1d illustrates the use of this notation. Of particular interest is our representation of double-resection events, which we shall discuss shortly. Our statistical model defines a probability distribution over *X*, and we shall subsequently use maximum likelihood estimation (MLE) to estimate branch lengths.

#### Data generation process

To define our statistical model for the data *X*, we will define the data generating process in two steps. In the first step, lineage tracing data, which we call *Z*, without any missingness is generated. In other words, we pretend that there are no sequencing dropouts, epigenetic silencing, or double-resection events. In the second step, we *retrospectively* analyze the data *Z* and determine what entries *R should have gone missing*, thus obtaining *X*. We will formalize this two-step process below, but it is an important conceptual leap because by decoupling the missing data mechanism from the CRISPR/Cas9 mutation process, the distribution of *Z* is easier to analyze since independence between sites holds, whereas this is **not** true for *X*. Indeed, sequencing dropouts, epigenetic silencing and double-resections all create correlations between different sites of *X*, since if there is a *−*1 in some site of a barcode, a *−*1 is more likely to also be present in an adjacent site of the barcode, as seen in Figure 1d.

To provide intuition for how this two-step process works, it is instructive to consider a simpler statistical model: one in which we roll *n* dice, and each die roll has a probability *ϕ* of being made to go missing. There are two equivalent ways to model this: in the first way, *before* rolling a die, we decide whether it will go missing. If so, we do not roll the die. In the second way, we first roll *all* the dice, obtaining die rolls *Z*, and *afterward* determine which die rolls to hide (by replacing their values with *−*1) to obtain the observed data *X*. Both models are equivalent in that they induce the same probability distribution for the observed data *X*. However, the second approach is richer because it provides us with extra random variables in *Z* – essentially, counterfactuals for the values of the missing entries. This way of thinking is convenient for analyzing CRISPR/Cas9 lineage tracing data, as it simplifies the mathematical derivation of the MLE.

#### Statistical model

Let us thus formally define the statistical model for CRISPR/Cas9 lineage tracing data *X*. Our model is quite general and thus more broadly applicable to other irreversible mutation models; the nuances of CRISPR/Cas9 lineage tracing data, such as double-resection events, will be handled during the data *encoding* stage, sparing the model from this burden. We start by defining a model for *Z* – the process *without* missing data.

Let 𝒯 be the given single-cell tree topology over the *n* cells – that is to say, a leaf-labeled, rooted tree whose leaves correspond to the *n* cells. The model is parameterized by branch lengths 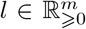 for 𝒯; here *m* is the number of edges of 𝒯. Let 𝒯_*l*_ be the single-cell chronogram obtained by assigning the branch lengths *l* to 𝒯. The generative model for *Z* is as follows:

1. The *k* sites are independent and identically distributed.
2. Each site evolves down the tree 𝒯_*l*_ following a continuous-time Markov chain defined as follows:
  a. Each site starts uncut at the root (in state 0).
  b. CRISPR/Cas9 cuts each site with a rate of *c*, where *c* is a nuisance parameter.
  c. When a site is cut, it takes on state *s* ∈ ℕ = {1, 2, 3, …} with probability *q*_*s*_ ⩾ 0, where *q* is nuisance probability distribution over ℕ. Note that ∑_*s*∈ℕ_*q_s_*=1
  d. Once a state is cut, it can no longer be cut again.

Explicitly, the transition rate matrix for our model – which is infinite-dimensional – is given by:

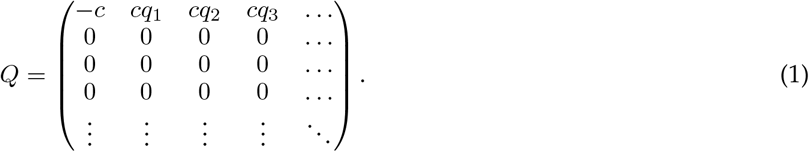

Thus, the transition probability matrix in time *t* for our model is given by:

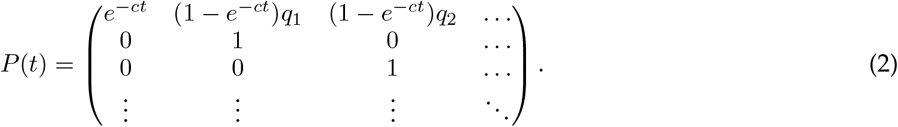

The root distribution is given by *π* = (1, 0, 0, 0, …). Note that this is a general irreversible mutation model that is not specifically tailored to CRISPR/Cas9 lineage tracing data.

We denote by *θ* = (*l, c, q*) the full set of model parameters including the nuisance parameters. By the end of the process we obtain *Z*, the states for all the leaves in the tree. More generally, the process described so far defines a statistical model 𝒫= {*p*_*θ*_} for lineage tracing data representing the character states for *all* the nodes in the tree 𝒯; we denote these data by 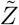, which is a random matrix of size *ñ*×*k* where *ñ*is the number of nodes in 𝒯 (including internal nodes).

#### Modeling missing data

The final step is to model the missing data mechanism occurring over the tree, which explains how to obtain *X* from *Z*. For this, following Little et al. (2017), let 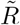 be the binary response/missingness mask matrix of size *ñ*× *k* indicating which entries of 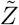 ought to (not) go missing (with 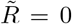 for missingness and 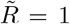 for response). Given 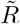 and 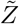, the actual data generated by the process at all nodes of the tree, which we denote as 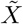, is:

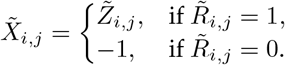

The subset of 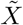 corresponding to the leaf nodes is then *X* – what we observe and were looking to model.

We need to specify how 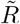 is jointly distributed with 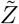. We will take a very general approach of Mealli and Rubin (2015), and assume that missing data is *missing always completely at random* (MACAR), meaning the missing data mask 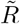 is independent from the lineage tracing data 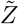. Formally, we assume a joint factorization 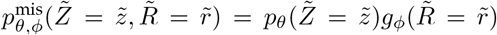, where *ϕ* are the parameters of the missing data mechanism (such as the sequencing dropout probability, epigenetic silencing rate, or temporal proximity of cuts required to induce a double-resection). However, we assume nothing else about the distribution 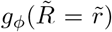, meaning that we take a non-parametric approach to missing data. In particular, the entries of the missing data mask 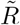 may be correlated, as in sequencing dropouts, epigenetic silencing and double-resection events. Lastly, we will assume that *ϕ* and *θ* are *distinct*, meaning that the value of *θ* puts no constraints on the value of *ϕ* and vice versa; formally (*θ, ϕ*) ∈ Θ ×Φ for some sets Θ and Φ. The MACAR and distinctness assumptions are together called *ignorability* of the missing data mechanism, and are crucial for justifying maximum likelihood estimation with missing data (Mealli and Rubin, 2015), as we will do shortly. By assuming ignorability – rather than trying to model the missing data mechanism – our model remains simple and decoupled from the nuances of CRISPR/Cas9 mutation process. This is in contrast to competing approaches such as LAML (Mai et al., 2024) which explicitly model heritable missing data, leading to more complex statistical models and slower optimization procedures.

#### Encoding double-resection

An important use of ignorable missing data is to implicitly model double-resection events. Recall that in a double-resection event, two sites *i* and *j* in the same barcode are cut close in time, resulting in the whole segment of sites between *i* and *j* to disappear, and a shared indel to be created spanning all sites *i* through *j*. Double-resection events are quite common and can be identified from the sequencing reads. Just like epigenetic silencing events, double-resection events are heritable. Prior work has chosen to represent double-resection events explicitly using complex indel states called ‘indel tracts’ as in GAPML (Feng et al., 2021). Unfortunately, this substantially complicates branch length estimation since complex correlations between barcoding sites need to be modeled. Instead, we propose a novel encoding of double-resections which is computationally tractable and does not compromise the quality of branch length estimates. Mathematically, we choose to encode and model double-resection events implicitly via an ignorable missing data mechanism, as follows: if concurrent cuts at sites *i* and *j* cause a double-resection, then sites *i* + 1 through *j* − 1 inclusive become ignorable missing data, as illustrated in Figure 1d. Note that sites *i* + 1 through *j* − 1 could have had mutations after the double-resection event in a reality with no missing data; these are precisely the counterfactuals modeled by 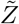. We choose not to model the fact that the indel created at sites *i* and *j* is shared; instead, we just retain the indels created by the model without missing data, i.e. two independent indels drawn from *q* as if double-resections did not exist in the first place. As we will show later in a targeted experiment, this representation and treatment of double-resection events leads to accurate branch length estimates.

We have now defined a statistical model 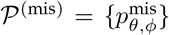 for the lineage tracing data *X* with missingness. The full list of random variables defined is outlined in Table 1. We now proceed to discuss how this statistical model for *X* aligns with the biology behind the CRISPR/Cas9 lineage tracing assay, and how it is misspecified.

**Table 1.** Random variables used throughout. Upper case letters with tilde denote random matrices of size ñ × k with one row per node in the tree while capital letter without tilde denote the submatrix of size n × k corresponding to the leaf nodes.

| Random Variables | Sizes | Meaning |
| --- | --- | --- |
| $\tilde{Z}, Z$ | $\tilde{n} \times k, n \times k$ | Lineage tracing states without missing data |
| $\tilde{R}, R$ | $\tilde{n} \times k, n \times k$ | Binary response (1)/missingness (0) mask |
| $\tilde{X}, X$ | $\tilde{n} \times k, n \times k$ | Lineage tracing states with missing data |

#### Discussion of model assumptions

First, let us discuss the assumption that the sites in 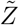 are independent. Since we choose to model sequencing dropouts, epigenetic silencing and double-resection events as missing data mechanisms, the missing data they introduce do not violate the independence of sites in 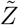 (instead, it is the sites in 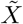 that are correlated). Secondly, CRISPR/Cas9 uses a different guide RNA sequence for each target site in a given barcode, so target sites within the same barcode evolve independently. Thirdly, barcodes are integrated randomly into the genome, so that they are far enough apart to interact with CRISPR/Cas9 independently. The only source of non-trivial model misspecification comes from our assumption that double-resections create two independent alleles at the two cut sites, when in fact the resulting indel should be shared and thus correlated, as seen in Figure 1d. This independent treatment of correlated indels may seem inappropriate, but it can be viewed as a form of composite-likelihood, which in general is known to retain consistency for parameter estimation under weak assumptions (Varin et al., 2011; Prillo et al., 2023). As we show later, when paired with conservative maximum parsimony (CMP), our method can produce highly accurate estimates of branch length even with large double-resection events. Thus, overall, independence of sites in 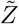 turns out to be a reasonable assumption for branch length estimation.

Now let us discuss the assumption that sites in 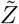 are identically distributed. Due to the local state of chromatin, some lineage tracing barcodes might be more prone to CRISPR/Cas9 cutting than others, leading to different cut rates across sites. Site rate variation has received attention in the field of statistical phylogenetics, where it has led to the development of more sophisticated methods, such as Yang (1995); Kalyaanamoorthy et al. (2017); Minh et al. (2021). However, models that assume equal rates across sites still perform well and have been used extensively, such as the seminal work of Whelan and Goldman (2001). Modeling site rate variation also leads to slower runtimes or risks overfitting. Therefore, in this work we will assume that all sites are cut at the same (unknown) rate, leaving analysis of more sophisticated models for future work.

The Markov assumption governing site evolution is standard for analyzing molecular data, and is the workhorse of Statistical Phylogenetics, enabling complex yet tractable statistical models. Hence, we adopt it in our work.

Regarding the ignorability of the missing data mechanism, we must consider the three different sources of missing data: sequencing dropouts, epigenetic silencing, and double-resections. Ignorability is true for sequencing dropouts, since all barcodes are equally likely to be dropped out during RNA-sequencing, regardless of their indel state, and the dropout probability is distinct from *θ*. Ignorability can also hold for epigenetic silencing. For example, suppose that barcodes get silenced at each node *v* of the tree 𝒯 with some probability *ϕ*_*v*_. In this case, the parameter of the missing data mechanism would be the vector of probabilities *ϕ* of dimension *ñ*. It is distinct from *θ*, and 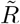 is independent of 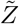 for any *θ, ϕ*, so that ignorability holds. On the other hand, suppose that the probability of epigenetic silencing at node *v* of 𝒯_*l*_ depends on the branch lengths, for example *ϕ*_*v*_ = 1 −exp(−*tη*) where *t* is the distance from the parent of *v* to *v*, and *η* is a rate parameter. In this case, the epigenetic silencing mechanism is not ignorable because *ϕ* is not distinct from *θ*. For similar reasons, missing data created by double-resection events is generally not ignorable. However, because it hinders scalability and avoids imposing potentially-misspecified parametric assumptions on the epigenetic silencing mechanism and on double-resection events, we also assume they are ignorable. Any other missing data mechanisms, should they exist, are also treated as ignorable. As we show later, this ignorable model of missing data yields accurate branch lengths despite misspecification, showing its suitability.

While we assume that the state probabilities *q*_1_, *q*_2_, *q*_3_, … are shared across all sites, this assumption can be relaxed to allow for site-specific probabilities 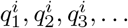 for each site *i*. Indeed, as we show below, thanks to our use of CMP, these state probabilities are irrelevant for computing the MLE because they appear as additive constants in the log-likelihood function.

### Maximum Likelihood Estimation of Branch Lengths

Having defined a statistical model for lineage tracing data *X* and discussed the suitability of the modeling assumptions made, we proceed to show how to perform maximum likelihood estimation of branch lengths under this model. Crucially, we show how to deal with unobserved ancestral states, missing data, and the nuisance parameters *c* and *q*.

We first deal with unobserved ancestral states. Let *x* be the observed value of *X*. The maximum likelihood estimator (also abbreviated as ‘MLE’) of branch lengths is given by:

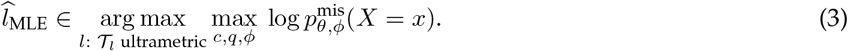

Here, *ultrametric* means that all leaves in the tree have the same distance from the root. This condition is imposed because all cells are sampled at the same time, but it is possible to relax this condition to account for temporal sampling of cells.

#### Unobserved ancestral states

To deal with unobserved ancestral states, we reconstruct most – but not all – of the ancestral states with a *conservative* version of maximum parsimony. Concretely, we only reconstruct ancestral states that are unambiguous under all maximum parsimony reconstructions consistent with the evolutionary process, as described in detail in **C**onservative **M**aximum **P**arsimony in Appendix. In other words, we only reconstruct ancestral states which we are confident about. Ambiguous states are not reconstructed – represented with the new symbol `NONE` (not to be confused with the missing data state *−*1) – and marginalized over by considering all possibilities, just like in an ignorable missing data mechanism. We call this reconstruction the ‘*CMPR*’, which stands for **C**onservative **M**aximum **P**arsimony **R**econstruction. Concrete examples are illustrated in Appendix Section Conservative Maximum Parsimony, where it is shown that CMP in fact agrees with naive maximum parsimony when there are no missing data. Thus, CMP only makes a difference when there are missing data. Some intuitive properties of naive maximum parsimony are also satisfied by CMP: for instance, if two leaves have the same state *x* (in particular, if both have the missing state *−*1), then their parent will have state *x* in the CMPR. It is only when one node has state *s >* 0 and the other node has the missing state *−*1 that CMPR and naive maximum parsimony may disagree. Whereas naive maximum parsimony will impute the parent node with state *s* or 0 (which are the only two options that are valid under the CRISPR-Cas9 mutation process), CMP may impute with the symbol `NONE`. When this happens, it is because CMP is unsure whether the parent node has state *s* or 0.

All in all, with CMP’s partial ancestral state reconstruction, what we have is a missing data mechanism (with missing data state `NONE`) on top of the original missing data mechanism (with missing data state *−*1). We assume both are ignorable, so that this is mathematically equivalent to a single ignorable missing data mechanism. Therefore, in what follows, we drop the `NONE` symbol and replace it with the missing data state *−*1, which thus now stands for sequencing dropouts, heritable missing data, double-resection missing data, *and* ambiguous ancestral states. All these missing data are treated as ignorable and hence marginalized out during MLE.

While it makes intuitive sense for CMPR to propagate the missing state *−*1 up the tree when it is shared by both nodes (as it is consistent with a heritable missing data event), the attentive reader may note that in the case when the true model has abundant sequencing dropouts but no heritable missing data, our CMP approach will incorrectly propagate missing data (*−*1) up the tree, which is clearly not an accurate ancestral state reconstruction. While in this border case the CMPRs are clearly incorrect, they do not bias branch length estimation, for the following reason. Informally, these propagated missing states *−*1 can be viewed as ignorable missing data. Formally, augment the missing data mechanism 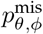 previously described such that at the end of the process, after 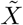 is generated, we consider all internal nodes all of whose leaves have missing data, and mark their states as missing (*−*1). This updates 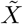 by propagating missing data up the tree as in CMPR. **Crucially, this missing data mechanism is a function of the original missing data mask, therefore it is ignorable if the original missing data mechanism is**. This way, CMPR can be viewed as reconstructing ancestral states under this augmented missing data mechanism, and CMPR is clearly accurate at propagating missing states in this case. It is thus intuitively clear that branch length estimation using our CMPR algorithm is unbiased even when there is no heritable missing data and abundant sequencing dropouts. We illustrate this extreme case in our benchmarking section, where we simulate data with 60% sequencing dropouts and 0% epigenetic silencing, and confirm that the error of our branch length estimation method is nearly zero.

Another explanation for why our method works even when it may incorrectly propagate missing data up the tree is as follows: when our CMP approach propagates missing data (*−*1) up the tree, these reconstruction can equivalently be interpreted as `NONE`, that is to say, as ambiguous reconstructions. Since there is no mathematical difference in the treatment of imputed *−*1s and the `NONE` state (both are treated as ignorable missing data and marginalized out), we thus see that propagating *−*1 up the tree is the right thing to do as it is equivalent to marking a state as ambiguous. Note that this equivalence between *−*1 and ambiguous states only holds because, unlike other methods like LAML and TiDeTree, we chose not to model missing data as its own separate state *s >* 0.

Having discussed how CMPR is accurate, and how missing data *−*1 and ambiguous states `NONE` are mathematically equivalent (thus we may replace all `NONE` by *−*1), we next show in Theorem A.4 that CMP has the key property that missing data states *−*1 can be marginalized out trivially. Importantly, the likelihood is essentially as easy to compute as if there were no missing data. Letting 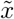 be the CMPR of ancestral states (with `NONE` replaced by *−*1 as discussed, since it is mathematically equivalent), the MLE amounts to:

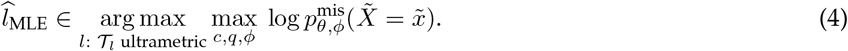

We assume that the solution to the optimization problem Eq. (3) is close to that of Eq. (4). As we have argued, and as we demonstrate later through simulation, CMP introduces essentially no bias, unlike naive maximum parsimony, and hence the reconstructions in 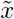 are reliable for the purpose of branch length estimation. Note, however, that as argued previously the ancestral states imputed as *−*1 will be inaccurate if there are significant sequencing dropouts and no heritable missing data; said imputations are then better interpreted as the ambiguous `NONE` state.

#### MLE with missing data

We next deal with missing data. The theory for dealing with ignorable missing data is well established (Rubin, 1976; Mealli and Rubin, 2015; Little et al., 2017), but we reproduce the necessary derivations here for a self-contained exposition; the key result is that under ignorability, we can treat the non-missing entries of 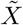 as having been sampled from their marginal distribution.

We start by defining 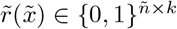 to be the missing data mask corresponding to 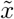. We will use the following notation:

Definition 1 Let *a* and *b* be matrices of the same dimensions, with *b* being binary. We denote by *a*[*b*] the vector obtained by concatenating the entries of *a* for which the corresponding entries in *b* equal 1. *A*[*b*] is similarly defined for a random matrix *A* of the same dimensions as *b*. We still index *a*[*b*] and *A*[*b*] as matrices for convenience, but it is important to note that they are just a *subset* of the matrix entries.

The following result establishes that we can ‘ignore’ missing data, as done in Rubin (1976); Little et al. (2017): Proposition 1 For all 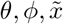 it holds that:

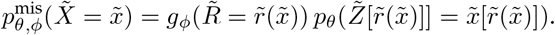

Importantly, since *ϕ* is distinct from *θ*, then *ϕ* and *θ* can be optimized independently in Eq. (4). Therefore, the MLE reduces to:

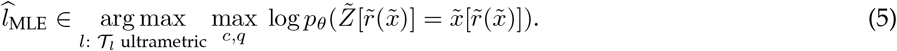

*Proof*. We have the following identities, each explained in turn below:

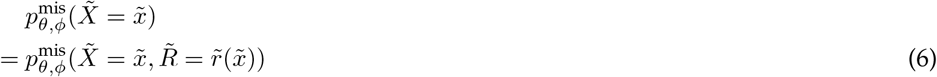

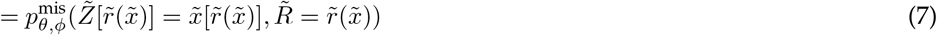

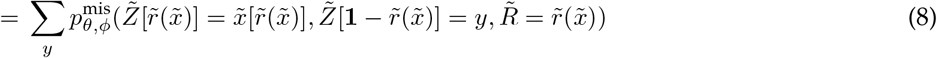

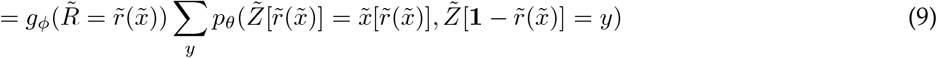

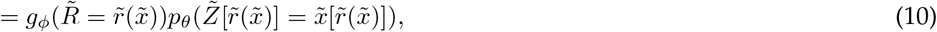

where identity 6 is because 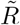 is a function of 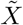; identity 7 follows from the definition of 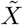; identity 8 follows from marginalizing out the missing data; identity 9 follows from the MACAR factorization; identity 10 follows from marginalizing out the missing data again. The expression **1** denotes an all-ones matrix of the same dimensions as 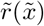.

#### Nuisance parameters

Finally, it remains to deal with the nuisance parameters *c* and *q*. To do this, we first unclutter notation by using the standard notation 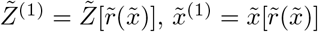, and so on (Mealli and Rubin, 2015) (note that the dependence on 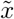 is now implicit), using the superscript ‘1’ to subset the non-missing entries of the object. Let *V* be the vertex set of 𝒯, and *E* the edge set of 𝒯. Since 𝒯 is rooted, we give edges their natural orientation pointing away from the root. For a node *v* and site *i*, we denote by gpa_*i*_(*v*) the first ancestor of *v* which has a non-missing state at site *i* (read as ‘grandparent’). The log-probability in the MLE objective in Eq. (5) is then given by

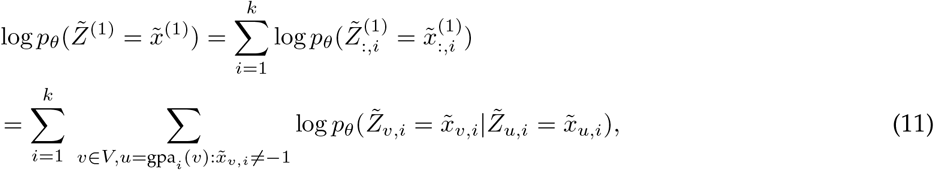

where the first equality follows from the independence of sites. The second equality requires using combinatorial properties of the CMPR, and is proved in Theorem A.4 in Appendix Section Conservative Maximum Parsi-mony. Finally, letting *l*_*uv*_ be the length of the path from *u* to *v* in the tree, we can compute the probability of an observed transition between nodes *u → v* where *u* = gpa_*i*_(*v*) as follows:

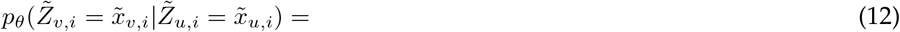

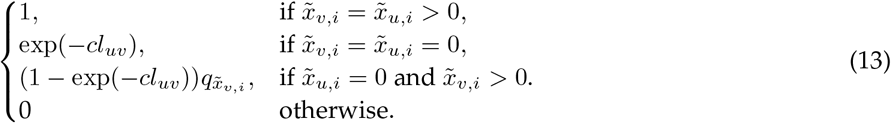

Plugging this into Eq. (11), we arrive at

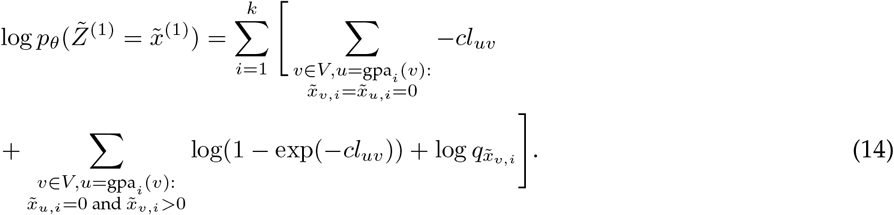

Note that in Eq. (14), the term 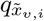 does not depend on *l*, so we can drop it from the objective function and hence, crucially, the MLE does not depend on *q*. To finish, let us denote by uncuts(*u* → *v*) and cuts(*u* → *v*) the number of characters that go uncut and cut, respectively, on the path from (*u, v*). In other words,

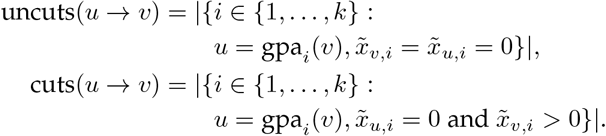

The MLE problem hence simplifies to

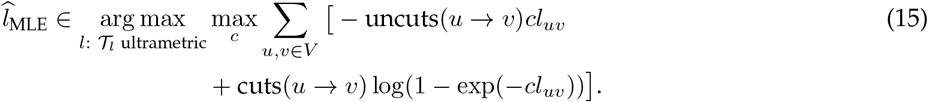

Although the MLE no longer depends on *q*, it still depends on the nuisance parameter *c*. It is a well known fact that branch lengths *l* and cut rate parameter *c* are unidentifiable without further assumptions, since they can be scaled up and down by the same constant without changing the likelihood. To resolve this ambiguity, without loss of generality, we assume that the chronogram has depth exactly equal to 1, which is equivalent to assuming that without loss of generality the duration of the experiment is normalized to 1. In other words, we solve for:

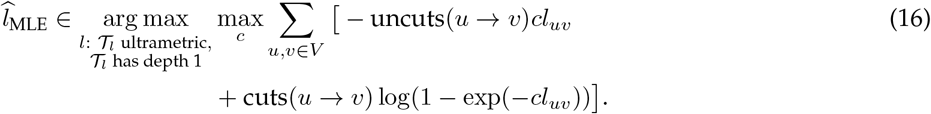

This is equivalent to solving the MLE problem in Eq. (15) using a cut rate of *c* = 1, and afterwards scaling the chronogram to unit depth. Since the MLE problem in Eq. (15) with *c* = 1 is a convex optimization problem in the variables{*l*_*uv*_ : (*u, v*) ∈ *E*}, it can be readily solved with off-the-shelf convex optimization solvers. When we implement the method, we exclude any terms in the objective where uncuts(*u* → *v*) = 0 or cuts(*u* → *v*) = 0 since they do not contribute, leading to a computation graph which has size essentially *O*(*n*) rather than *O*(*n*^2^) and thus leads to significantly faster runtime. This concludes optimization under the proposed model. In our implementation, for trees with 400 leaves and as many as 150 characters, the optimizer takes just a few seconds. In contrast, a method like TiDeTree takes several *hours* on a tree with 700 leaves (Seidel and Stadler, 2022). We leverage the cvxpy Python library (Diamond and Boyd, 2016; Agrawal et al., 2018) and the CLARABEL and SCS (O’Donoghue et al., 2016) solvers as the backend. We find that CLARABEL is faster; in case it fails, we fallback to the SCS solver.

### Regularizing the MLE

Due to limited lineage tracing capacity, it is not unusual for some cells in the population to have the exact same lineage tracing character states. When this happens, the single-cell tree topology will contain subtrees whose leaves all have the same character states. We call these *homogeneous* subtrees. As we show next, the MLE will estimate branch lengths of 0 for these subtrees. Moreover, the result is very general: it does not depend on whether ancestral states have been reconstructed with maximum parsimony or marginalized out, and it does not depend on the continuous-time Markov chain model. Concretely, we prove the following result:

#### Theorem 1

(Homogeneous Proper Subtree Collapse). *Consider any continuous-time Markov chain model (for example, the Jukes-Cantor model of DNA evolution, the WAG model of amino acid evolution (Whelan and Goldman, 2001), or the CRISPR/Cas9 lineage tracing model in this paper with fixed c, q). Let p*_*l*_ *be the associated probability measure when running this Markov chain on* 𝒯_l_ . *Let* 𝒮 *be a proper subtree of* 𝒯 *(meaning that it is distinct from* 𝒯*). Let z be states for the leaves of the tree such that all leaves of* 𝒮 *have the same state, and let* 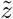 *be a reconstruction of the ancestral states of z where all nodes of* 𝒮 *have the same state (as in a maximum parsimony reconstruction). Let l be any ultrametric branch lengths for* 𝒯. *Then there exist other ultrametric branch lengths l*^*′*^ *for* 𝒯*that satisfy:*

a. *All the branch lengths of* 𝒮 *are zero under l*^*′*^.
b. 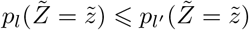.
c. 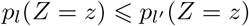.

*Proof*. See Appendix Section Proof of Subtree Collapse Theorem for a proof. The proof relies on a probabilistic coupling argument.

As a consequence, the MLE can always collapse homogeneous proper subtrees. We coin this phenomenon *subtree collapse*. More generally, we observed empirically that the MLE defined in Eq. (16) tends to estimate many branch lengths as 0, a phenomenon which we coin *edge collapse*.

Subtree collapse, and more generally edge collapse, are undesirable because it suggests that some cells have divided arbitrarily fast. To address this issue, and more generally to make the MLE robust in low-information regimes where there is little information to support branch lengths, we propose to combine two simple regularizers. The first is to impose a minimum branch length *ϵ*. This can be easily accomplished by adding the minimum branch length constraint to the MLE optimization problem of Eq. (16). The second is to add pseudocounts to the data, in the form of *λ* cuts and *λ* uncuts per branch. This is like pretending that we have observed *λ* characters getting cut on each edge, as well as *λ* characters remaining uncut on that edge. Intuitively, this regularizes the branch lengths towards trees that look more neutrally evolving. This also has the benefit of making the optimization problem strongly convex, ensuring there is a unique solution. With this, our method falls into the category of penalized likelihood methods (Kim and Sanderson, 2008) like GAPML (Feng et al., 2021), where the MLE is stabilized with a regularizer to aid in low-information regimes. Other alternatives include Bayesian methods such as TiDeTree (Seidel and Stadler, 2022), but these suffer from higher computational costs.

Incorporating our two regularizers to the optimization problem, we obtain:

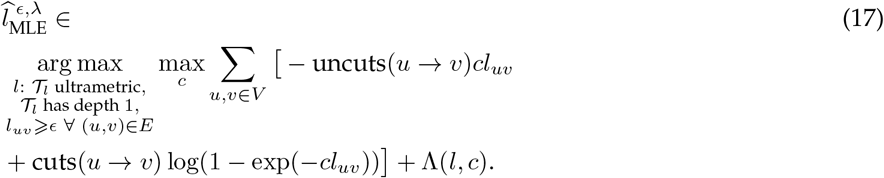

where Λ(*l, c*) = Σ _*u,v ∈E*_ *[λcl*_*uv*_ + *λ* log(1 *−*exp(*−cl*_*uv*_)] is the likelihood of the pseudocounts; therefore, using pseudocounts is equivalent to performing MAP estimation with the prior Λ(*l, c*) on the branch lengths.

Optimization problem Eq. (17) is equivalent to solving the MLE problem Eq. (15) while adding *λ* pseudocounts to the edges and imposing the constraints *c* = 1 and *l*_*uv*_ ⩾ *ϵd* ∀ (*u, v*) ∈ *E* where *d* is the depth of 𝒯_*l*_, and then finally normalizing the tree to have a depth of 1. This is still a convex optimization problem in the {*l*_*uv*_ : (*u, v*) ∈ *E*} variables and can thus still be solved easily with off-the-shelf convex optimization solvers. Note that technically, since leaf nodes do not correspond to cell division events but rather to a point *between* cell division events, the minimum branch length constraint does not strictly apply to pendant edges in the tree. A good compromise is thus to use a minimum branch length of *ϵ/*2 for pendant edges, which aligns more closely with the expected length of pendant edges assuming a cell division time of *ϵ*. We use this refined constraint of *ϵ/*2 for pendant edges.

The value of *ϵ* is interpretable and can thus be selected based on prior biological knowledge. Concretely, if the shortest amount of time that can pass between two consecutive cell division events is known to be approximately *t*, and if the length of the experiment is *T*, then *ϵ* can be set to *t/T*. Imposing a minimum branch length completely avoids edge collapse.

Pseudocounts are a popular regularization technique with the advantage that the regularization strength *λ* is interpretable in terms of a data perturbation. We explore using values of *λ* equal to 0 (no said regularization), 0.1 (small regularization) and 0.5 (large regularization). Note that pseudocounts do not guarantee that a minimum branch length is satisfied, which is why we use both forms of regularization.

Other regularization strategies are possible, such as the one used by GAPML (Feng et al., 2021) where large differences in branch lengths are discouraged via an 𝓁_2_ penalty in log-space. However, we use a minimum branch length and pseudocounts because their regularization strengths *ϵ* and *λ* are interpretable and thus easier to select without needing to perform an extensive grid search with cross-validation.

All in all, our ConvexML method is based on a general irreversible mutation model paired with conservative maximum parsimony and regularization. This means that it should be more broadly applicable to any irreversible mutation model with a large enough state space and approximately ignorable missing data mechanisms. Nuances of the CRISPR/Cas9 lineage tracing data, such as double-resections, are handled with data encoding instead. Missing data mechanisms are also not explicitly modeled, retaining simplicity. These aspects underlie the scalability and accuracy of our approach.

## Results

### Benchmarked Models

We benchmark several models, including ablations (versions of our model without certain components, specifically regularization), oracles, and baselines. First, on controlled simulations with 400 leaves, we benchmark our MLE using ground truth ancestral states, conservative maximum parsimony, or naive maximum parsimony, which we denote as 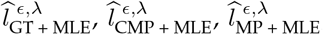, respectively. We set *ϵ* to a small value of *ϵ* = 0.01 and vary *λ* ∈{0, 0.1, 0.5}. The value of *ϵ* = 0.01 was chosen to mimic biological knowledge of minimum cell division times (and may vary depending on the application); Supplementary Figure 2 shows that indeed *ϵ* = 0.01 is a reasonable lower bound on cell division times in our simulated trees.

We also benchmark a baseline model 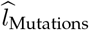 which sets each branch length to the number of mutations on the branch. This baseline shows what happens if we ignore the mutation model. To avoid assigning a length of zero to edges without mutations, we set their edge length to 0.5 instead. To make the tree ultrametric, we extend all tips to match the depth of the deepest leaf. Finally, we scale the tree to have a depth of 1. Analogously to the MLE, we benchmark three versions of this model: 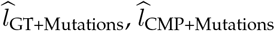 and 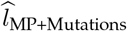 respectively. When using CMP, mutations that are mapped to *m* edges are made to contribute 1*/m* mutations to each of those edges.

In addition, we benchmark the recently proposed LAML method (Mai et al., 2024), which is based on maximum likelihood estimation much like our method; however, unlike our method, (1) LAML marginalizes out the ancestral states, requiring that the user provides the state distribution *q*_1_, *q*_2_, *q*_3_, … as a hyperparameter (which must also be finite-dimensional), and (2) LAML does not perform regularization of branch lengths. We ran LAML by providing the fixed tree topology as input and disabling topology search; hence only branch lengths are estimated. Exact details are shown in Appendix Section LAML Command Line. As we will see, LAML is both slower (due to marginalization of ancestral states) and less accurate (due to omission of regularization) compared to our method ConvexML.

Finally, on the real intMEMOIR lineage tracing dataset (Chow et al., 2021), we benchmark ConvexML against LAML and TiDeTree using the ground truth tree topology. As a note, we encountered difficulty running TiDeTree on simulated data and therefore excluded it from the simulation benchmarks; other recent works such as Mai et al. (2024) also do not benchmark TiDeTree on their simulations.

### Simulated Data Benchmark

#### Simulation steps

We evaluate the performance of the models on simulated data. Our simulations involve three steps: (1) simulating a ground truth single-cell chronogram, (2) simulating lineage tracing data on the chronogram, and (3) sampling leaves. Briefly, our ground truth single-cell chronograms are simulated under a birth-death process where the birth and death rates are allowed to change with certain probability at each cell division, allowing us to model changes in the fitness of subclones. The simulation ends when a specified population size is reached; if the whole population of cells dies, we retry. Our simulated lineage tracing assay controls the number of barcodes, the number of sites per barcode, the indel distribution, the CRISPR/Cas9 site-specific mutation rates, and the rates of epigenetic silencing and sequencing dropouts. Sampling leaves is done by specifying the number of sampled leaves.

#### Evaluation tasks

Using our simulation framework, we set out to evaluate the performance of the different methods on several tasks, and across varying qualities of lineage tracing data. We considered the following three evaluation tasks and associated metrics:

1. **“Internal node time” task:** Mean absolute error at estimating the time of the internal nodes in the induced chronogram.
2. **“Ancestral lineages” task:** Relative error at estimating the number of ancestral lineages halfway (time-wise) through the induced chronogram.
3. **“Fitness estimation” task:** Spearman correlation at estimating the fitness of each sampled cell. The ground truth fitness of a cell is defined in our simulations as the difference between its birth and death rates. We use the Neher et al.’s LBI estimator (Neher et al., 2014) as implemented in the jungle package^†^ to derive fitness estimates from our single-cell chronograms, as in the work of Yang et al. (2022).

We also profile the runtime of each method, which can be considered a fourth evaluation metric.

#### Using reconstructed tree topology

Furthermore, since in real applications we do not have access to the ground truth single-cell tree topology, we also benchmark the performance of the branch length estimation methods when the tree topology must first be estimated. To do this, we first estimate the single-cell topology using the Maxcut solver based on Snir and Rao (2006) from the `Cassiopeia` package (Jones et al., 2020) – which is a supertree method based on triplets – and then apply the branch length estimator. We chose the Maxcut solver since it is a top performer at the triplets correct and Robinson-Foulds metric commonly used to evaluate topology reconstruction quality (Supplementary Figure 3) and runs much faster than the ILP algorithm. We resolve multifurcations into binary splits using a Huffman-tree algorithm (Huffman, 1952) wherein subtrees with the least size are merged first. Since the ground truth topology and the reconstructed topology differ, the internal node time task becomes harder to evaluate. To resolve this discrepancy, we create estimates for the time of each internal node *v* in the ground truth tree by taking the mean time of the MRCA (most recent common ancestor) in the reconstructed tree of all pairs of leaves whose MRCA in the ground truth tree is *v*.

#### Questions of interest

With our simulated data benchmark we seek to answer the following questions: (i) Does our method significantly outperform the naive baseline which estimates branch lengths as the number of mutations? (ii) How much does accuracy drop when the single-cell topology must be estimated first – as in real applications – as compared to when the ground truth topology is known? (iii) Does regularization stabilize branch length estimates, particularly in low-information regimes? (iv) Does CMP provide accurate branch length estimates, as compared to naive maximum parsimony and to the oracle method which has access to ground truth ancestral states? How does it compare to the more sophisticated LAML method, which attempts to marginalize out the ancestral states? (v) Similarly, does our treatment of double-resection event enable unbiased branch length estimates?

#### Simulating chronograms

To answer the above questions, we perform the following simulations. We simulate 50 chronograms with 40,000 extant cells each, scaled to have a depth of exactly 1. We then sample exactly 400 leaves from each tree, thus achieving a sampling probability of 1%. The simulation is performed using a birth-death process with rate variation to emulate fitness changes, and is described in detail in Appendix Section Tree Simulation Details. The resulting trees display nuanced fitness variation, as can be seen in Supplementary Figure 4, which shows 9 of our 50 ground truth induced chronograms.

#### The default parameter regime

For each chronogram, we simulate a lineage tracing experiment with 13 barcodes and 3 target sites per barcode, for a total of 39 target sites; we choose 3 target sites per barcode based on the technology used in Chan et al. (2019); Quinn et al. (2021); Yang et al. (2022). The per-site CRISPR/Cas9 mutation rates are splined from real data to achieve an expected 50% of mutated entries in the character matrix, and are shown in Supplementary Figure 5. We consider 100 possible indel states with a non-uniform probability distribution *q*, as in real data, such that some states are much more common than others. Concretely, the *q*_*s*_ are taken to be the quantiles of an exponential distribution with scale parameter 10^*−*5^. This non-uniform probability distribution is shown in Supplementary Figure 6. The indels are encoded as integers between 1 and 100 inclusive. We introduce sequencing dropouts and epigenetic silencing missing data mechanisms such that on average 20% of the character matrix is missing, with 10% arising from sequencing dropouts and 10% from epigenetic silencing. The missing data they introduce is represented with the integer *−*1. Epigenetic silencing occurs similarly to CRISPR/Cas9 mutations, happening with a fixed rate during the whole length of the experiment (which as discussed before, is in fact *not* an ignorable missing data mechanism). Double-resections are also simulated; they occur whenever two sites in the same barcode are cut before a cell divides, as illustrated in Figure 1d. When a double-resection event occurs at positions *i* and *j*, we create an identical indel at positions *i* and *j* with integer encoding 10^8^ + 2^*i*^ + 2^*j*^ such that we can identify the endpoints of the double-resection as in real data; positions *i* + 1 through *j* −1 go missing and are thus set to −1. If more than two sites in a barcode are cut before the cell divides, *i* and *j* are taken to be the leftmost and rightmost of these sites in the barcode. We call the collection of all these lineage tracing parameters the ‘default’ lineage tracing parameter regime. The branch length estimation methods are then evaluated on the three benchmarking tasks described above using the 50 simulated trees.

#### Varying parameters

We then proceed to repeat the benchmark, this time varying each of the lineage tracing parameters in turn. This allows us to explore lineage tracing datasets with varying levels of quality, as in real life. We vary the number of barcodes in the set {3, 6, 13, 20, 30, 50}, the expected proportion of mutated character matrix entries in the set {10%, 30%, 50%, 70%, 90%}, the number of possible indels in the set {5, 10, 25, 50, 100, 500, 1000}, and the expected missing data fraction in the set {10%, 20%, 30%, 40%, 50%, 60%}, always keeping the expected sequencing missing data fraction at 10%, and adjusting the expected heritable epigenetic missing data fraction accordingly.

#### Assessing ancestral state reconstruction

To specifically analyze the bias of CMP, we perform a second targeted experiment where we simulate a very large number of lineage tracing characters – 100,002 characters consisting of 33,334 barcodes of size 3 – on one of the previously described trees and then apply the MLE using either (i) ground truth ancestral states, (ii) conservative maximum parsimony, or (iii) naive (standard) maximum parsimony. It is the scalability of our method that allows us to perform such a large-scale experiment to analyze its bias. Since the state space size is crucial for parsimony methods, we vary the number of indel states in the set {1, 2, …, 9} ∪{10, 20, …, 100}, so that in particular we explore a binary tracer with a unique indel state. Since missing data makes ancestral state reconstruction challenging, we use 60% expected missing data, with half coming from epigenetic silencing and the other half from sequencing dropouts. Since double-resection events indirectly increase the state space size (as they encode the indices of the outer sites), we initially exclude them from the experiment. Finally, unlike the default parameter regime, to ensure that the main source of model misspecification comes from the reconstruction of unobserved ancestral states, we use the same cut rate for all sites, thus excluding site rate variation from this specific experiment; the cut rate is chosen such that an observed entry of the character matrix has 50% change of being mutated. As we have theoretically argued previously, CMPR works even when there are abundant sequencing dropouts and no heritable missing data (epigenetic silencing). To further support this empirically, we repeated the experiment and used 60% sequencing dropouts and 0% epigenetic silencing.

#### Assessing the effect of double-resections

To specifically explore the effect of double-resections, we repeat the previous experiment which has 100,002 characters but turning on double-resection events, which introduces more missing data but also indirectly increases the state space size. Moreover, to exacerbate the effect of double-resections, we perform a third version of the experiment where we not only simulate double-resections, but also increase the cassette size from 3 to 10 (while keeping the total number of characters controlled and equal to 100,000 by using 10,000 barcodes).

### Performance Comparison

#### MLE outperforms the baseline estimator

The performance of each model on the default parameter regime, with and without access to the ground truth topology, is shown in Figure 2. We can see that all variants of our MLE method outperform the baseline estimator. Our method takes on average approximately 2 seconds per tree, and is thus significantly faster than LAML, which takes approximately 5 minutes per tree. All these patterns hold across all lineage tracing parameter regimes, shown in Supplementary Figures 7, 8, 9 and 10.

**Fig. 2.**
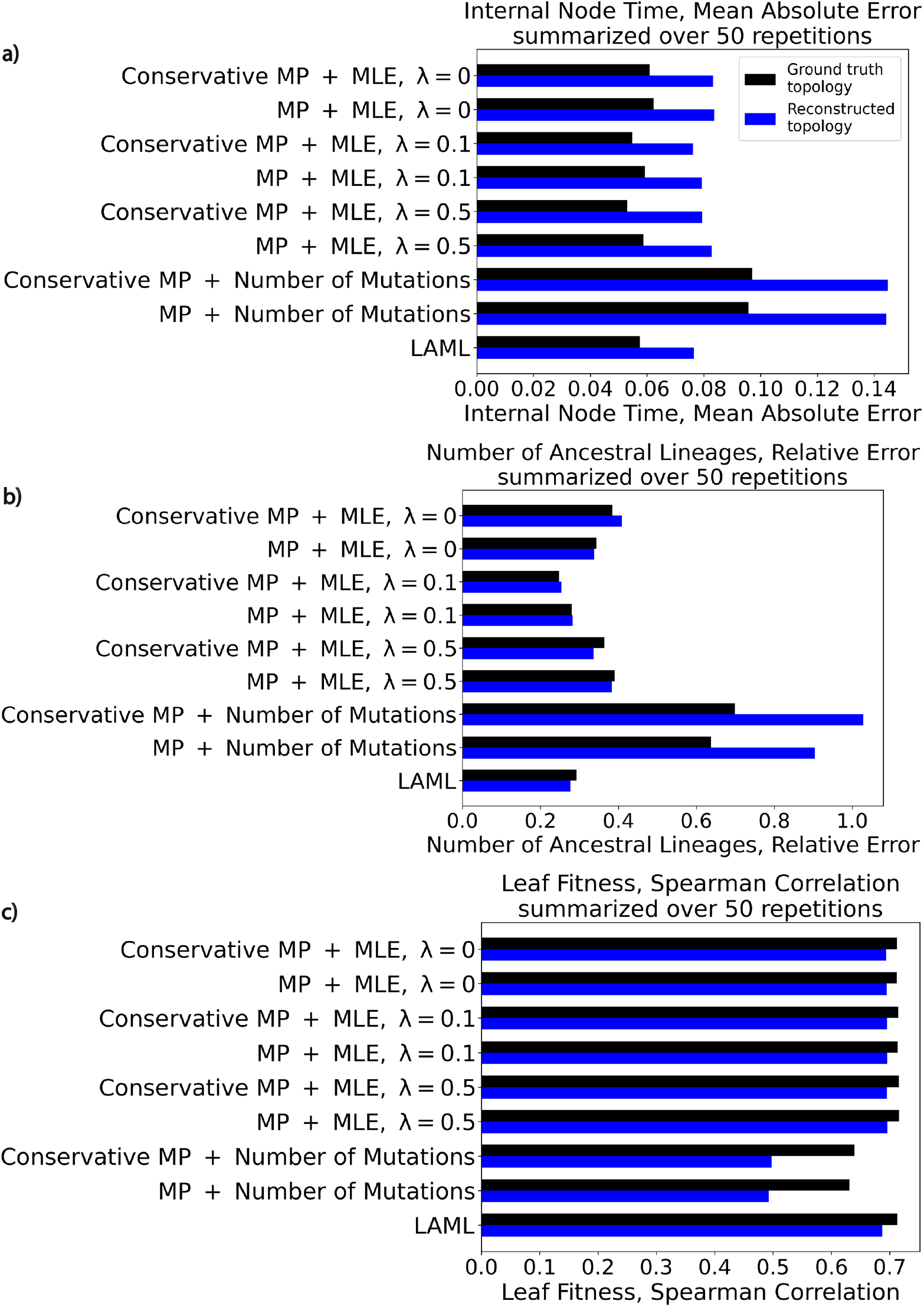
Performance of branch length estimation methods under the default parameter regime. For each branch length estimation method, we show the performance both with and without access to the ground truth topology (in black and blue respectively). When the topology is not provided, it is reconstructed with the Maxcut solver from the Cassiopeia package (Jones et al., 2020). Branch length estimators are evaluated on three tasks: a) Estimation of the times of the internal nodes in the tree, b) Estimation of the number of ancestral lineages midway through the experiment, c) Estimation of the relative fitness of the cells in the population.

#### The effects of using reconstructed tree topologies

We find that using reconstructed single-cell tree topologies – as opposed to ground truth topologies – consistently degrades performance on the internal node time and fitness prediction tasks, as expected. On the ‘ancestral lineages’ task, however, this is not always the case. For example, on the default regime (Figure 2) using a value of *λ* = 0.5 in our MLE leads to better performance with the reconstructed topology. We attribute this to the MLE being over-regularized and the bias introduced by regularization cancelling off with the bias of the topology reconstruction algorithm. The interaction between topology, branch length estimation procedure, and downstream metric is thus nuanced, and errors in the topology and branch length estimation steps can compound, or more interestingly, cancel each other.

#### The effects of regularization

Using regularization improves performance most noticeably in low-information regimes such as when the amount of missing data increases, the expected proportion mutated decreases, or the number of lineage tracing barcodes decreases, as seen in Supplementary Figures 7, 8, and 9. This is expected since the purpose of regularization is precisely to stabilize branch length estimates in low-information regimes. Other than low-information regimes, the regularization strength *λ* makes the most difference on the ancestral lineages task, where a small amount of regularization *λ* = 0.1 provides the best results. Generally, using some level of regularization as opposed to none appears to be beneficial across the board.

#### Conservative maximum parsimony vs. naive maximum parsimony

CMP tends to outperform naive maximum parsimony, the gap being most noticeable in high-information regimes such as when the number of lineage tracing barcodes is 50. This makes sense, as only at high sample sizes the risk of the estimator becomes dominated by the bias rather than by the variance. For the ‘internal node time’ task as we increase the number of barcodes, the performance of conservative maximum parsimony remains close to the performance of the oracle model with access to ground truth ancestral states. In contrast, the performance of naive maximum parsimony improves more slowly. For example, the performance of naive maximum parsimony with 50 barcodes is comparable to the performance of CMP with just 30 barcodes. We found that our method performs better than the more sophisticated LAML method which marginalizes out the ancestral states. We attribute the improved performance of our method to using regularization in the form of pseudocounts and a minimum branch length, which LAML lacks. Indeed, the performance of LAML is most similar to the performance of our method ConvexML without pseudocounts. Thus, despite marginalizing out the ancestral states, LAML performs worse than our method, since proper use of regularization outweighs the importance of marginalizing out ancestral states. This provides further evidence that CMP is a powerful idea for branch length estimation in single-cell lineage tracing – marginalizing out ancestral states is not that important, and (2) careful regularization is important. As a note, we encountered difficulty running TiDeTree on simulated data and thus excluded it from the simulation benchmark; however, we do report comparisons against TiDeTree on the real intMEMOIR dataset in a subsequent section.

In our second experiment, we dissected the bias of our CMP approach by simulating a very large number of characters (100,002) on one of our trees and evaluating the performance of the MLE when using either (i) ground truth ancestral states, (ii) conservative maximum parsimony, or (iii) naive maximum parsimony. In the first version of the experiment, we excluded double-resection events since they indirectly increase the state space size. Recall that we use 60% missing data, with 30% coming from epigenetic silencing and 30% coming from sequencing dropouts. The results are shown in Figure 3. As the number of indel states increases, the bias of CMP seems to converge to nearly 0 in a geometric fashion. The mean absolute error achieved is negligible when compared to the error for the default lineage tracing regime as seen in Figure 2, which is at least 0.05. Naive maximum parsimony, on the other hand, shows large bias with a MAE of 0.06, which is comparable to the MAEs seen in Figure 2. Since CRISPR/Cas9 lineage tracing systems are characterized by their large state space, this targeted experiment shows the suitability of CMP for branch length estimation. When we repeated the experiment with 60% sequencing dropouts and 0% epigenetic silencing, we observed similar results, shown in Supplementary Figure 11. This confirms that CMPR works well even when there is no heritable missing data, as we argued on theoretical grounds previously.

**Fig. 3.**
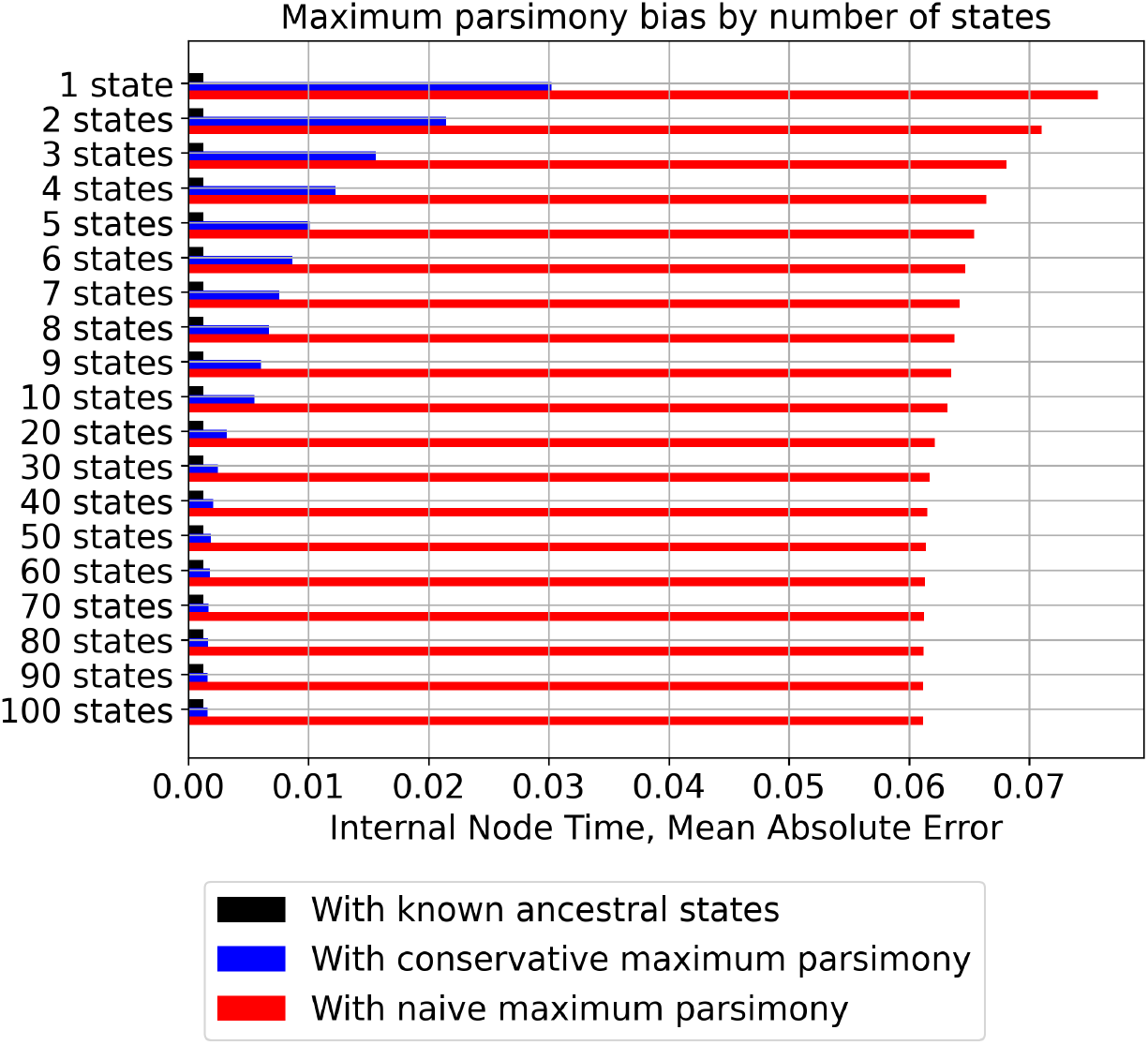
Conservative maximum parsimony enables negligible bias. By simulating a massive number of lineage tracing characters on a tree and performing MLE, we explored the bias of our CMP approach as a function of the number of indel states. We can see that as the number of indel states increases, the bias quickly goes away, unlike for naive maximum parsimony. Because CRISPR/Cas9 lineage tracing systems are characterized by their large state space, this makes CMP a powerful tool in this setting.

#### The effect of double-resections

We finished by turning on double-resection events in our previous simulation with 100,002 characters. The results are shown in Supplementary Figure 12a. The indirectly increased state space size due to double-resections improves performance of our branch length estimator for a low number of states, while the misspecified treatment of double-resections minimally degrades performance for larger state space sizes. Increasing the barcode size from 3 to 10 while keeping the number of characters roughly the same magnifies these trends, as shown in Supplementary Figure 12b. Even with these barcodes of length 10, the error of CMP remains close to 0 for large state space sizes. These results show that paired with CMP, independent modelling of sites is accurate even in the presence of double-resection events – which is the key to the scalability of our method as compared to previous methods such as GAPML (Feng et al., 2021). Similar results were observed when using 60% sequencing dropouts and 0% epigenetic silencing.

One may wonder what happens if one uses a different representation of double-resection events, e.g., if one encodes them by assigning the indel state to *all* sites *i* through *j*, instead of only to sites *i* and *j*; in Figure 1d, this would mean assigning state 5 to sites 1, 2, and 3, instead of assigning state 5 to sites −1 and 3 and 1 to site 2. This is the current preprocessing behaviour of the `Cassiopeia` package (Jones et al., 2020). The results are shown in Supplementary Figure 13. As we can see, performance deteriorates dramatically, even for the model with ground truth ancestral states. Intuitively, our novel representation of ancestral states is effective because it entails exactly two cutting events, which is the correct number.

#### Benchmarking on smaller and larger trees

We further benchmarked the methods on larger trees with 2, 000 leaves instead of 400 (details in Appendix Section Tree Simulation Details). The results are shown in Supplementary Figures 14, 15, 16, and 17. Our method performs comparably to or better than LAML in terms of accuracy while being significantly faster (10 ~ 100 ×). Similarly, we benchmarked the methods on smaller trees with just 40 leaves instead of 400 (details in Appendix Section Tree Simulation Details). The results are shown in Supplementary Figures 18, 19 and 20. Our method again performs comparably to or better than LAML in terms of accuracy while being significantly faster (10 ~100 ×). Generally, LAML’s poor performance in lower information regimes continues to hold, which we attribute to the lack of regularization.

#### Recommendation

Based on these results, our recommendation to practitioners is to use CMP with the regularized MLE 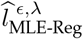 setting *ϵ* to the best known lower bound on the minimum time between cell division events, and explore values of *λ* of 0.0, 0.1 and 0.5 as we did. For example, if cells are expected to take at least 1 day to divide and the experiment last 60 days, our recommendation would be to set the minimum branch length to *ϵ* = 1*/*60 ≈ 0.016. Character-level cross-validation is a promising avenue to automate the choice of *λ* (and even *ϵ*) and we leave it to future work. Double-resections should be encoded using our novel representation, with −1 (i.e., missing data) at the internal sites of the double-resection, and the indel state duplicated at the flanking sites, as seen in Figure 1d. To ensure that the indel state created by a double-resection event is distinct from any indel created by a standard single-site cut, we recommend using a modified encoding such as 10^8^*s* + 2^*i*^ + 2^*j*^, where *s* is the original encoding of the indel and *i, j* are the sites of the double-resection. (In particular, note that in our simulations, we are considering a worse-case scenario where only one indel state is possible during a double-resection.)

For users who desire to obtain a notion of uncertainty for their branch length estimates, character-level bootstrapping, as well as sampling of the regularization strength *λ* may provide a simple way of obtaining uncertainty estimates for the branch lengths. The speed of our method enables this, unlike other competing methods such as LAML (Mai et al., 2024) which provide no uncertainty estimates and are prohibitively slow to bootstrap from. TiDeTree (Seidel and Stadler, 2022) provides uncertainty estimates via posterior samples from the MCMC chain. It is important to note that uncertainty estimates are uncertain themselves (since all models are misspecified), so ultimately complementary validations (for instance with prior biological knowledge) are typically performed. Moreover, single-cell lineage tracing analysis typically comprise hundreds of trees, so that uncertainty is usually controlled by averaging results over many trees and not a single one. In particular, randomness is controlled by considering many trees each with a point estimate, rather than a single tree with many estimates. See, for example, the analyses in recent publications (Quinn et al., 2021; Yang et al., 2021; Simeonov et al., 2021).

### Real intMEMOIR dataset

Finally, we benchmarked our method ConvexML on the real intMEMOIR dataset (Chow et al., 2021), which contains 106 ground truth single-cell chronograms obtained by ocular inspection. While these trees are small, they contain ground truth topology and branch lengths, making them a powerful resource to benchmark competing methods. For each tree, we evaluated the internal node time mean absolute error (MAE) as we did on our simulation benchmark. We chose *λ* = 0.1 pseudocounts following the results of our simulation benchmark, and selected a minimum branch length of *ϵ* = 0.15 using our prior biological knowledge on mouse embryonic stem cells: the intMEMOIR datasets were obtained by letting cells grow for a total of 54 hours, and cell divisions typically take at least 8 hours in this culture, hence *ϵ* = 8*/*54 ≈ 0.15. We report the scatterplot of errors of ConvexML vs. TiDeTree and ConvexML vs. LAML in Figure 4. ConvexML outperforms both methods: it outperforms TiDeTree on 60% of trees and LAML on every one of them. We attribute our improved performance to (1) the negligible bias of CMP, and (2) the effective use of regularization of our method. In particular, LAML’s lack of regularization significantly degrades branch length estimates. Supplementary Figure 21 shows sample branch length estimates on the first of the intMEMOIR clones. Despite being an MLE method, ConvexML produces high-quality estimates which are quantitatively comparable to the more sophisticated Bayesian TiDeTree method. In contrast, despite marginalizing out ancestral states, the MLE method LAML collapses branch lengths as predicted by our Theorem 1 due to the lack of regularization. Hence, LAML’s lack of regularization leads to poor quality branch length estimates.

**Fig. 4.**
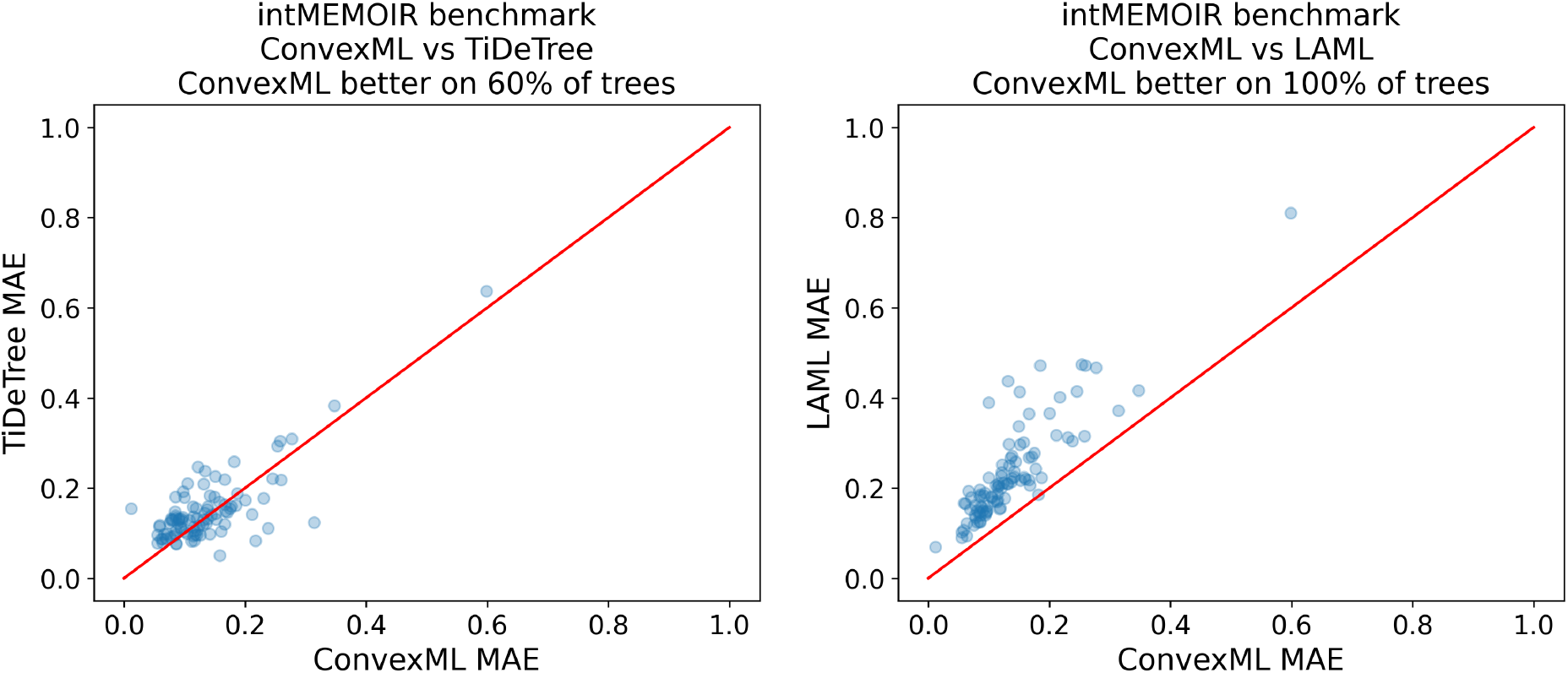
ConvexML outperforms TiDeTree and LAML on the intMEMOIR dataset. On 60% of the trees, ConvexML attains a lower branch length estimation error (‘node time MAE’ metric) than TiDeTree. ConvexML outperforms LAML on all trees, which is explained by LAML’s lack of regularization.

In Supplementary Figure 22, we show ConvexML’s error for each node as a function of node depth. As we can see, the error is stable across different node depths, showing the robustness of our evaluation metric.

## Discussion

In this work, we introduced the first scalable and accurate method to estimate branch lengths for single-cell tree topologies, refining them into single-cell chronograms. Our method consists of regularized maximum likelihood estimation under a general irreversible mutation model, together with regularization and a conservative version of maximum parsimony. We call our method ‘ConvexML’ to highlight the tractability of the resulting optimization procedure.

Our use of parsimony is central to the scalability of our approach, and we showed through detailed experiments that the bias introduced by conservative maximum parsimony is essentially negligible thanks to the large state space of CRISPR/Cas9 mutations. As a note, our method can still be applied to irreversible models with smaller state spaces too, but the bias might become noticeable and undesirable. The use of parsimony to speed up maximum likelihood-based tree estimation methods was also recently applied successfully to scaling up estimation of traditional reversible mutation models to pandemic-scale data in the MAPLE method (‘MAximum Parsimonious Likelihood Estimation’) (De Maio et al., 2023). With MAPLE, the authors report estimating SARS-CoV-2 phylogenies up to a thousand times faster and using a hundred times less memory. The key insight is that for large trees with very similar sequences, likelihood concentrates on parsimonious solutions, therefore shortcuts can be made when computing likelihoods during Felsentein’s pruning algorithm (Felsenstein, 1981). In our case of ConvexML, the concentration of likelihood on parsimonious solutions is due to the irreversibility and high diversity of alleles. The use of conservative maximum parsimony in particular also has its analogous counterpart for time-reversible models, where it is known as the DOWNPASS algorithm in MacClade (Maddison and Maddison, 2000) and has been used to provide a baseline for reconstructing ancestral scenarios (Ishikawa et al., 2019).

To deal with the specific nuances of CRISPR/Cas9 data – such as double-resections and epigenetic silencing – we proposed using an ignorable missing data mechanism and a new encoding of double-resection events. This keeps our model simple and decoupled from the particularities of CRISPR/Cas9 lineage tracing data. This is in stark contrast with recently proposed methods such as LAML (Mai et al., 2024) and TiDeTree (Seidel and Stadler, 2022), which explicitly model epigenetic silencing, or GAPML (Feng et al., 2021), which explicitly models double-resection events, leading to more complex optimization procedures.

Our regularization approach based on a minimum branch length and pseudocounts plays an important role for stabilizing estimates in lower information regimes. To the best of our knowledge, the use of pseudocounts is not widespread in statistical phylogenetics. We recently also used pseudocounts successfully to regularize the estimation of site-specific rate matrices in our SiteRM model (Prillo et al., 2024), which extends the popular LG model (Le and Gascuel, 2008) by allowing for a different rate matrix at each column of a multiple sequence alignment. Our pseudocount-based regularization approach may therefore be of broad interest – it could be applied to regularize other methods like LAML (Mai et al., 2024) and see use in more traditional statistical phylogenetic analyses.

While our method was designed and benchmarked for the CRISPR/Cas9 setting, our method should in fact be broadly applicable to any evolutionary model with irreversible mutations and an (approximately) ignorable missing data mechanism. Indeed, we emphasize the following: (1) Our method does not assume any particular missing data mechanism (we just assume that the mutation outcomes are decoupled from the propensity for missing data), so our treatment of missing data is in fact agnostic to the CRISPR/Cas9 case, unlike prior works like TiDeTree and LAML. (2) Our treatment of double-resection events (which are idiosyncratic to the Cas9 system) is a pre-processing step (i.e., data representation step) and not a specific statistical component of our model. In effect, we ‘reduce’ or ‘transform’ double-resection events into independent single-site events.

The simplicity and scalability of our method enable numerous extensions, some of which we are currently pursuing. For instance, we are working on accounting for site rate variation (i.e., the fact that different target sites evolve at different rates) by allowing a different cut rate for each site; we expect that as lineage tracing technologies get better, site rate variation may become more and more relevant. The resulting optimization problem is essentially identical to the original one if the site-specific rates are known. If the site-specific rates are not known, they can be estimated using a simple coordinate ascent procedure wherein the branch lengths and site rates are optimized in turn. Another important extension is to estimate branch lengths for trees that are not ultrametric. This arises in applications where cells are sampled at different moments in time.

We also plan to explore the use of character-level cross-validation to automatically perform hyperparameter selection; although we aimed to make our hyperparameters as interpretable as possible, a cross-validation scheme would nonetheless decrease the burden on the user. Lastly, we look forward to leveraging our branch length estimator to infer the transcriptional dynamics of cell populations, a problem that is akin to learning amino acid substitution rate matrices and which has its own rich history in the field of statistical phylogenetics. By doing so, we seek to shed light into the transcriptional dynamics of cancer development; in this application, branch lengths are crucial to obtain good quantitative estimates.

All in all, our approach to branch length estimation – with CMP, simple yet effective treatment of missing data, and regularization – provides a state-of-the-art branch length estimator procedure for single-cell phylogenetics, and a branch length estimation method for irreversible mutation processes more broadly.

## Funding Acknowledgement

This research is supported in part by NIH grants R56-HG013117 and R01-HG013117, and the European Union Council (ERC, Tx-phylogeography, 101089213). Views and opinions expressed are however those of the authors only and do not necessarily reflect those of the European Union or the European Research Council Executive Agency. Neither the European Union nor the granting authority can be held responsible for them.

## Data Availability Statement

No new real data were generated in this research. ConvexML is implemented in the convexml open source python package, available at: https://github.com/songlab-cal/ConvexML. Code to reproduce all results in our paper is also contained in that repository.

## SUPPLEMENTARY MATERIAL

Supplementary file contains the Appendix and all supplementary figures.

