## Supplementary Information for "ConvexML: Fast and accurate branch length estimation under irreversible mutation models, illustrated through applications to CRISPR/Cas9-based lineage tracing"

### APPENDIX

#### Proof of Subtree Collapse Theorem

We prove a slightly more general version of Theorem 1, which (essentially) implies that subtree collapse happens also when there is missing data and the missing data mechanism is ignorable. In phrasing this Theorem we use notation similar to Rubin's missing data notation (Rubin, 1976), but we make no reference to missing data mechanisms in order to make the result more self-contained. In what follows, let $L(\mathcal{T})$ denote the leaf set of tree $\mathcal{T}$ and $V(\mathcal{T})$ denote the node set of $\mathcal{T}$.

Theorem A.1 (Homogeneous Proper Subtree Collapse, Generalized). *Consider any continuous-time Markov chain model with state space $\mathbb{Z}_{\geqslant 0}$ (for example, the Jukes-Cantor model of DNA evolution, the WAG model of amino acid evolution (Whelan and Goldman, 2001), or the CRISPR/Cas9 lineage tracing model in this paper with fixed $c, q$). Given a tree topology $\mathcal{T}$ and branch lengths $l$ for $\mathcal{T}$, let $\tilde{Z} : V(\mathcal{T}) \to \mathbb{Z}_{\geqslant 0}$ be the stochastic process obtained by running this CTMC down the tree $\mathcal{T}_l$, and let $p_l$ be the associated probability measure for $\tilde{Z}$. Let $Z : L(\mathcal{T}) \to Z_{\geqslant 0}$ be the restriction of $\tilde{Z}$ to the leaves of $\mathcal{T}$. Let $v \in V(\mathcal{T})$ be different from the root, and let $\mathcal{S}$ be the subtree of $\mathcal{T}$ rooted at $v$. Let $L' \subseteq L(\mathcal{T})$ be a subset of the leaves of $\mathcal{T}$. Let $z^{(1)} : L' \to \mathbb{Z}_{\geqslant 0}$ be a state assignment for this subset $L'$ of the leaves of $\mathcal{T}$ such that all leaves of $\mathcal{S}$ in $L'$ have the same state – meaning that $z^{(1)}(l_1) = z^{(1)}(l_2)$ for all $l_1, l_2 \in L' \cap L(\mathcal{S})$ – and at least one leaf of $\mathcal{S}$ is assigned a state by $z^{(1)}$ – meaning that $L' \cap L(\mathcal{S})$ is non-empty. Let $V' \subseteq V(\mathcal{T})$ be such that $V' \cap L(\mathcal{T}) \supseteq L'$. Let $\tilde{z}^{(1)} : V' \to \mathbb{Z}_{\geqslant 0}$ be an extension of $z^{(1)}$ – meaning that $\tilde{z}^{(1)}(l) = z^{(1)}(l)$ for all $l \in L'$ – and such that all nodes in $\mathcal{S}$ are assigned the same state – meaning that $\tilde{z}^{(1)}(v_1) = \tilde{z}^{(1)}(v_2)$ for all $v_1, v_2 \in V' \cap V(\mathcal{S})$. Suppose that $l$ are ultrametric branch lengths for $\mathcal{T}$. Let $\tilde{Z}^{(1)}$ be the restriction of $\tilde{Z}$ to $V'$ and let $Z^{(1)}$ be the restriction of $Z$ to $L'$. Then there exist other ultrametric branch lengths $l'$ for $\mathcal{T}$ that satisfy:*

*a) All the branch lengths of $\mathcal{S}$ are zero under $l'$.*

*b) $p_l(\tilde{Z}^{(1)} = \tilde{z}^{(1)}) \leqslant p_{l'}(\tilde{Z}^{(1)} = \tilde{z}^{(1)})$*

*c) $p_l(Z^{(1)} = z^{(1)}) \leqslant p_{l'}(Z^{(1)} = z^{(1)})$*

**Note**. It turns out that it is important that $\tilde{z}^{(1)}$ assigns a state to at least one leaf of $\mathcal{S}$ (part (a) of the Theorem is false otherwise), but it actually does not matter whether $z^{(1)}$ assigns a state to some leaf of $\mathcal{S}$ (part (b) is trivially true in this case).

*Proof of Theorem A.1.* Let $w$ be the parent of $v$ (which exists since by hypothesis $v$ is not the root of $\mathcal{T}$). Let $u$ be a leaf of $\mathcal{S}$ assigned a state by $z^{(1)}$ (i.e. $u \in L' \cap L(\mathcal{S})$), which exists by hypothesis. Consider the following operation on $\mathcal{T}_l$, which we call the *subtree collapse operation*: we take the nodes in $\mathcal{S}$ and set their depths to the depth of the tree, thereby 'collapsing' $\mathcal{S}$ down onto the leaf $u$. We denote by $l'$ the new branch lengths after this operation, and thus the new chronogram as $\mathcal{T}_{l'}$. The operation is illustrated in Figure A.1.

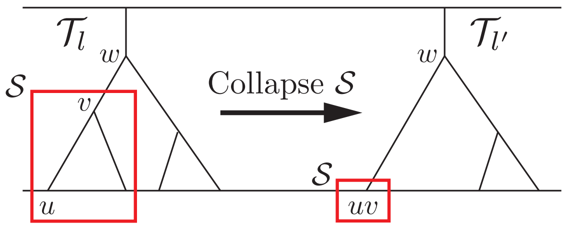


Fig. A.1. Subtree Collapse Operation

We claim that $l'$ satisfies the sought properties. It is clear that $l'$ are ultrametric branch lengths. To show that the likelihood of $z^{(1)}$ as well as of $\tilde{z}^{(1)}$ can only increase after the subtree collapse operation, we create a coupling between $p_l$ and $p_{l'}$, as follows:

1. Let the Markov chain run as usual down $\mathcal{T}_l$.
2. For the part of $\mathcal{T}_l$ that was not perturbed by the subtree collapse operation – meaning any point on the tree $\mathcal{T}$ which does not descend from $v$, nor is on the edge $(w, v)$ – copy all of its Markov chain transitions onto $\mathcal{T}_{l'}$, as in Figure A.2.

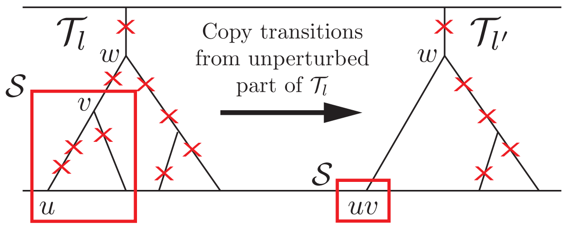


Fig. A.2. Copy of transitions from unperturbed part of $\mathcal{T}_l$ onto $\mathcal{T}_{\tilde{l}}$.

3. Copy the transitions on the path from $w$ to $u$ in $\mathcal{T}_l$ onto the same path in $\mathcal{T}_{l'}$, as in Figure A.3.

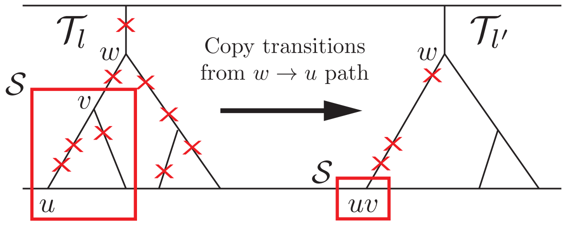


Fig. A.3. Copy of transitions from $w \to u$ path of $\mathcal{T}_l$ onto $\mathcal{T}_{l'}$.

The result of the two copy operations above results in transition operations getting created on all branches of $\mathcal{T}_{l'}$, and is depicted in Figure A.4.

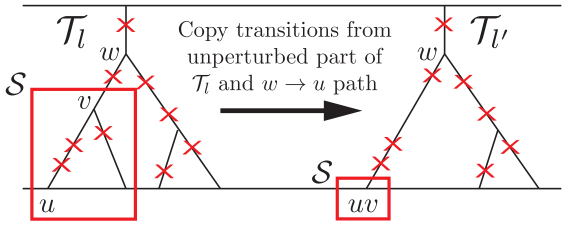


Fig. A.4. Copy of transitions from $\mathcal{T}_l$ onto $\mathcal{T}_{l'}$.

The key (trivial) observation is that the process induced over $\mathcal{T}_{l'}$ follows exactly $p_{l'}$. As such, let $p$ be the coupling thus created over $p_l(\tilde{Z}^{(1)})$ and $p_{l'}(\tilde{Z}^{(1)})$, and let $\tilde{Z}_l^{(1)}, \tilde{Z}_{l'}^{(1)}$ be the random variables corresponding to $\tilde{Z}^{(1)}$ under $p_l$ and $p_{l'}$ respectively. In other words, $p(\tilde{Z}_l^{(1)}, \tilde{Z}_{l'}^{(1)})$ is a probability density which satisfies

$$p(\tilde{Z}_l^{(1)}) = p_l(\tilde{Z}^{(1)}) \text{ and } p(\tilde{Z}_{l'}^{(1)}) = p_{l'}(\tilde{Z}^{(1)}).$$

We now claim that

$$Z_l^{(1)} = z^{(1)} \Rightarrow Z_{l'}^{(1)} = z^{(1)},$$

and, similarly,

$$\tilde{Z}_l^{(1)} = \tilde{z}^{(1)} \Rightarrow \tilde{Z}_{l'}^{(1)} = \tilde{z}^{(1)}.$$

Indeed, if $Z_l^{(1)} = z^{(1)}$ then this means that all the leaves of $\mathcal{S}$ which are assigned a state by $z^{(1)}$ have the same state as $u$, which is exactly the situation in $Z_{l'}^{(1)}$ by construction, thus $Z_{l'}^{(1)} = z^{(1)}$. Similarly, if $\tilde{Z}_l^{(1)} = \tilde{z}^{(1)}$ then it means that all the nodes of $\mathcal{S}$ which are assigned a state by $\tilde{z}^{(1)}$ have the same state as $u$, which is exactly the situation in $\tilde{Z}_l^{(1)}$ by construction, hence $\tilde{Z}_{l'}^{(1)} = \tilde{z}$; this is the step where it is important that at least one leaf of $\mathcal{S}$ be assigned a state by $\tilde{z}^{(1)}$. This completes the proof, for we obtain

$$p_l(Z^{(1)} = z^{(1)}) = p(Z_l^{(1)} = z^{(1)}) \leqslant p(Z_{l'}^{(1)} = z^{(1)}) = p_{l'}(Z^{(1)} = z^{(1)}) \tag{A.1}$$

and analogously

$$p_l(\tilde{Z}^{(1)} = \tilde{z}^{(1)}) = p(\tilde{Z}_l^{(1)} = \tilde{z}^{(1)}) \leqslant p(\tilde{Z}_{l'}^{(1)} = \tilde{z}^{(1)}) = p_{l'}(\tilde{Z}^{(1)} = \tilde{z}^{(1)}), \tag{A.2}$$

as claimed. □

### Conservative Maximum Parsimony

In what follows, we let $\mathcal{T}$ be a rooted tree topology with $n$ leaves and a total of $\tilde{n}$ nodes. We allow multifurcations (nodes with more than 2 children) and require that each internal node other than the root have at least 2 children (such that unifurcations are not allowed, except for the root node, as in a single-cell phylogeny). Let $V$ and $E$ be the vertex set and edge set of $\mathcal{T}$ respectively, and let $L$ be the set of leaves. We use $x : L \to \mathbb{Z}$ to denote the states of one specific character over the leaves of $\mathcal{T}$, including the missing data state "$-1$", and we use $\tilde{x} : V \to \mathbb{Z}$ to denote the states of that specific character over *all* the nodes of $\mathcal{T}$. We call the root of $\mathcal{T}$ simply 'root' when indexing.

We first define what it means for a state assignment $\tilde{x}$ to be *valid* under the irreversible CRISPR/Cas9 mutation process:

**Definition A.1 (Valid $\tilde{x}$)** We say that a state assignment $\tilde{x}$ is *valid* if it satisfies the following three properties:

- The root is uncut: $\tilde{x}_{\text{root}} = 0$.
- Heritability of missing data: $\forall (p, c) \in E, (\tilde{x}_p = -1 \Rightarrow \tilde{x}_c = -1)$.
- Irreversibility of the mutation process (not to be confused with the notion of irreversibility of a Markov Chain): $\forall (p, c) \in E, (\tilde{x}_p > 0 \Rightarrow \tilde{x}_c \in \{\tilde{x}_p, -1\})$.

An important quantity associated with a state assignment $\tilde{x}$ is its *parsimony score*:

**Definition A.2 (Parsimony score)** We define the *parsimony score* $\text{Par}(\tilde{x})$ of a state assignment $\tilde{x}$ as:

$$\text{Par}(\tilde{x}) = \sum_{(p,c) \in E} 1\{\tilde{x}_p \neq \tilde{x}_c\}, \tag{A.3}$$

where $1\{\cdot\}$ denotes the indicator function.

Unfortunately, we usually only know the state assignment $x$ for the leaves $L$ of the tree $\mathcal{T}$. Because of this, it is common to *reconstruct* the ancestral states $\tilde{x}$ from $x$ in some fashion. Since the mutation process is constrained, not every imputation of ancestral states is valid. We thus define what it means for $\tilde{x}$ to be a *valid reconstruction* of $x$:

Definition A.3 (Reconstruction $\tilde{x}$ of $x$) We say that $\tilde{x}$ is a *reconstruction* of $x$ if $\forall v \in L, \tilde{x}_v = x_v$.

Definition A.4 (Valid reconstruction $\tilde{x}$ of $x$) We say that $\tilde{x}$ is a *valid reconstruction* of $x$ if $\tilde{x}$ is valid, and if $\tilde{x}$ is a reconstruction of $x$.

Note that there always exists a valid reconstruction $\tilde{x}$ of $x$, by just setting all internal node states to $0$. However, we usually seek for the 'best' valid reconstruction, in some suitable sense. This notion of 'best' is commonly chosen to be the most parsimonious solution, which is the one that minimizes the parsimony score $\mathrm{Par}(\tilde{x})$:

Definition A.5 (Valid maximum parsimony reconstruction) We say that $\tilde{x}$ is a *valid maximum parsimony reconstruction* of $x$, abbreviated VMPR, if $\tilde{x}$ is a valid reconstruction of $x$ and $\tilde{x}$ achieves the lowest parsimony score (ties allowed) amongst all valid reconstructions of $x$; there may be multiple VMPRs for a given $x$.

We now give an algorithm to compute a VMPR from $x$. For this, we will need some definitions:

Definition A.6 (Set of leaf descendants) Given $v \in V$, we define $\mathcal{L}(v) \subseteq L$ as the set of leaves descending from $v$.

Definition A.7 (Subset of leaf states) Given a subset of leaves $L' \subseteq L$, we define $x_{L'} = \{x_u : u \in L'\}$. In particular, $x_{\mathcal{L}(v)}$ is the set of states at the leaves descending from $v$.

We are ready to state our algorithm:

**Algorithm 1 A Valid Maximum Parsimony Reconstruction**

1: **procedure** VMPR($\mathcal{T}$, $x$)
2:     **for** $v$ in postorder traversal of $\mathcal{T}$ **do**
3:         **if** $v$ is a leaf **then**
4:             $\tilde{x}_v \leftarrow x_v$
5:         **else if** $v$ is the root of $\mathcal{T}$ **then**
6:             $\tilde{x}_v \leftarrow 0$
7:         **else**
8:             Let $u_1, \ldots, u_c$ be the children of $v$.
9:             Let $S = \{\tilde{x}_{u_1}, \tilde{x}_{u_2}, \ldots, \tilde{x}_{u_c}\}$.
10:             **if** $S = \{s\}$ for some $s \in \mathbb{Z}$ **then**
11:                 $\tilde{x}_v \leftarrow s$
12:             **else**
13:                 **if** $-1 \in S$ **then**
14:                     **if** $S = \{-1, s\}$ for some $s \geqslant 0$ **then**
15:                         $\tilde{x}_v \leftarrow s$
16:                     **else**
17:                         $\tilde{x}_v \leftarrow 0$
18:                 **else**
19:                     $\tilde{x}_v \leftarrow 0$
20:     **return** $\tilde{x}$

We now show that Algorithm 1 indeed computes a VMPR, which we call $\tilde{x}^{\mathrm{VMPR}}$:

THEOREM A.2. *Algorithm 1 computes a VMPR $\tilde{x}$ of $x$.*

*Proof. Suppose otherwise. Consider the first step of the algorithm where the current state assignment (prior to executing the step) cannot be completed to any VMPR. We call this the 'failing' step. In the previous step, let $\tilde{x}$ be a VMPR compatible with the state assignment so far. We analyze several cases, depending on which line of the algorithm we fail on:*

- *We cannot fail in lines 4 nor 6 because $\tilde{x}$ is a valid reconstruction (Definition A.4).*
- *If we failed on line 11 then changing $\tilde{x}_v$ to $s$ in $\tilde{x}$ would remain a valid reconstruction and improve the parsimony score by at least $1$, contradiction.*
- *If we failed in line 15, then first note that $\tilde{x}_v \neq -1$ (or else by heritability of missing data $S = \{-1\}$, contradicting line 14). The only other option is $\tilde{x}_v = 0$. But then if we change $\tilde{x}_v = s$ in $\tilde{x}$ we still have a valid reconstruction and*

*the transition $0 \to 0$ going into $v$ becomes $0 \to s$, but at least one transition $0 \to s$ leaving $v$ becomes $s \to s$, meaning that the new state assignment must be VMPR and is consistent with the algorithm the step it failed, contradiction.*

- *We cannot fail in line 17 or 19 because there are at least two different non-missing states $s_1, s_2 \geqslant 0$. If one of these is a $0$, then $\tilde{x}_v = 0$ by validity. Otherwise, these two are distinct positive states, and again $\tilde{x}_v = 0$ by validity.*

*This concludes the proof.* □

We next give a direct, non-algorithmic characterization of the VMPR $\tilde{x}^{\text{VMPR}}$ computed by Algorithm 1:

**Proposition A.1 (Structure of $\tilde{x}^{\text{VMPR}}$)** Let $\tilde{x}^{\text{VMPR}}$ be the VMPR of $x$ computed by Algorithm 1. Then, for any internal node $v \in V$, the following properties hold:

1. If $v = \text{root}$, then $\tilde{x}_v^{\text{VMPR}} = 0$.
2. If $(v \neq \text{root}) \wedge (0 \in x_{\mathcal{L}(v)})$, then $\tilde{x}_v^{\text{VMPR}} = 0$.
3. If $(v \neq \text{root}) \wedge (0 \notin x_{\mathcal{L}(v)}) \wedge (\exists s_1, s_2 > 0 : s_1 \neq s_2, \{s_1, s_2\} \subseteq x_{\mathcal{L}(v)})$ then $\tilde{x}_v^{\text{VMPR}} = 0$.
4. If $(v \neq \text{root}) \wedge (x_{\mathcal{L}(v)} = \{-1\})$, then $\tilde{x}_v^{\text{VMPR}} = -1$.
5. If for some $s > 0$ we have $(v \neq \text{root}) \wedge (\{s\} \subseteq x_{\mathcal{L}(v)} \subseteq \{-1, s\})$, then $\tilde{x}_v^{\text{VMPR}} = s$.

Moreover, the five cases above are disjoint and exhaustive, so they completely characterize $\tilde{x}^{\text{VMPR}}$.

*Proof.* We first observe that the five cases are disjoint and exhaustive: case (1) handles the root node; case (2) handles an internal node with some leaf state of $0$; case (3) handles an internal node with no leaf state of $0$ but two distinct non-zero leaf states; case (4) handles an internal node with all missing leaf states. If none of these four first cases hold, then it means that it is an internal node and there exists some $s > 0$ such that $\{s\} \subseteq x_{\mathcal{L}(v)} \subseteq \{-1, s\}$, which is handled by case (5).

Now the proposition follows directly by induction on the nodes of $\mathcal{T}$ in postorder: The first three implications are true simply because $\tilde{x}^{\text{VMPR}}$ is valid; the fourth implication is true because all the $-1$ in the subtree at $v$ will be propagated up the tree by the algorithm; and the fifth implication is true because positive states take precedence over missing states in the algorithm, specifically, in line 15, so as states $-1$ and $s$ are propagated up the subtree at $v$, the state $s$ will make it to the top. □

Unfortunately, maximum parsimony imputation tends to lead to biased branch lengths, as shown in Figure 3. A key contribution of our work is the idea of a *conservative* maximum parsimony reconstruction, which gets rid of the maximum parsimony bias, while being computationally just as tractable as a typical maximum parsimony reconstruction. The idea of CMP is to just reconstruct ancestral states that all valid maximum parsimony reconstructions agree on, and leave the remaining ones without any state value, for which we shall use the new symbol `NONE`. Formally, we define:

**Definition A.8 (Conservative maximum parsimony reconstruction)** We say that $\tilde{x} : V \to \mathbb{Z} \cup \{\texttt{NONE}\}$ is the *conservative maximum parsimony reconstruction* of $x$, abbreviated CMPR, if for all nodes $v \in V$ it holds that:

- If there is some state $s$ such that $\tilde{x}'_v = s$ for all VMPR $\tilde{x}'$ of $x$, then $\tilde{x}_v = s$.
- If there is no state $s$ such that $\tilde{x}'_v = s$ for all VMPR $\tilde{x}'$ of $x$, then $\tilde{x}_v = \texttt{NONE}$.

Unlike the VMPR, the CMPR is unique by definition. We shall denote $\tilde{x}^{\text{CMPR}}$ the CMPR of $x$.

Figure A.5 shows a minimal example of CMP, while Figure A.6 shows a larger example. Just as for the VMPR, we can give a non-algorithmic characterization of the CMPR which will be convenient for analyzing its properties. Unlike the VMPR, the structure of the CMPR is more involved, and determining the state of some nodes requires looking up though its list of ancestors, which we define as follows:

**Definition A.9 (Ancestor)** Given nodes $g, v$ in a $\mathcal{T}$, we say that $g$ is an *ancestor* of $v$ if there is a path from $g$ to $v$ in $\mathcal{T}$. A node is considered an ancestor of itself.

We can now prove:

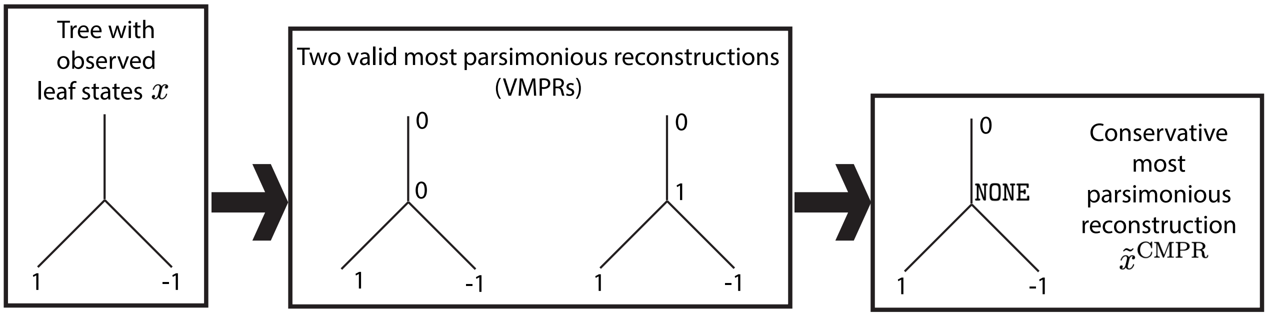


Fig. A.5. A minimal example of CMP. There exist two valid most parsimonious reconstructions (VMPRs) for the leaf states $x$ on the left. Importantly, these two VMPRs disagree on the state of the (non-root) internal node. In other words, the mutation to state 1 observed in the first leaf cannot be unambiguously mapped to an edge in the tree. Therefore, the conservative maximum parsimony reconstruction (CMPR) assigns this node the ambiguous label NONE, so as not to make an arbitrary mapping decision. This ambiguous state is thus marginalized out during MLE computations, avoiding the bias that would result from having maximum parsimony choose an arbitrary mapping, as shown in Figure 3.

THEOREM A.3 (Structure of the CMPR). *Let $\tilde{x}^{CMPR}$ be the CMPR of $x$. Then, for any internal node $v \in V$, the following properties hold:*

1. *If $v = root$, then $\tilde{x}^{CMPR}_v = 0$.*
2. *If $(v \neq root) \wedge (0 \in x_{\mathcal{L}(v)})$, then $\tilde{x}^{CMPR}_v = 0$.*
3. *If $(v \neq root) \wedge (0 \notin x_{\mathcal{L}(v)}) \wedge (\exists s_1, s_2 > 0 : s_1 \neq s_2, \{s_1, s_2\} \subseteq x_{\mathcal{L}(v)})$ then $\tilde{x}^{CMPR}_v = 0$.*
4. *If $(v \neq root) \wedge (x_{\mathcal{L}(v)} = \{-1\})$, then $\tilde{x}^{CMPR}_v = -1$.*
5. *If for some $s > 0$ we have $(v \neq root) \wedge (\{s\} \subseteq x_{\mathcal{L}(v)} \subseteq \{-1, s\}) \wedge (\exists g$ ancestor of $v$ different from the root such that $\{s\} \subseteq x_{\mathcal{L}(g)} \subseteq \{-1, s\}$ and $\exists u_1, u_2$ distinct children of $g$ such that $\{s\} \subseteq x_{\mathcal{L}(u_1)}, x_{\mathcal{L}(u_2)} \subseteq \{-1, s\})$, then $\tilde{x}^{CMPR}_v = s$.*
6. *If for some $s > 0$ we have $(v \neq root) \wedge (\{s\} \subseteq x_{\mathcal{L}(v)} \subseteq \{-1, s\}) \wedge (\nexists g$ ancestor of $v$ different from the root such that $\{s\} \subseteq x_{\mathcal{L}(g)} \subseteq \{-1, s\}$ and $\exists u_1, u_2$ distinct children of $g$ such that $\{s\} \subseteq x_{\mathcal{L}(u_1)}, x_{\mathcal{L}(u_2)} \subseteq \{-1, s\})$, then $\tilde{x}^{CMPR}_v = NONE$.*

*The first four properties above are analogous to that of Proposition A.1 satisfied by $\tilde{x}^{VMPR}_v$. Moreover, the six cases above are disjoint and exhaustive, so they completely characterize $\tilde{x}^{CMPR}$.*

*Proof.* *We first observe that the six cases are disjoint and exhaustive: these are the same cases as those of the VMPR structure Proposition A.1, except that the last case was split into the two cases* (5) *and* (6) *by pivoting on the truth value of a somewhat involved condition depending on the non-root ancestors of $v$.*

*We now show each of the implications. The first 3 will be a simple consequence of VMPRs being valid, and the fourth will be a simple consequence of VMPRs being most parsimonious. The last two cases, however, require some work. The proof for each implication is as follows:*

1. *All VMPRs $\tilde{x}$ have $\tilde{x}_{root} = 0$ since they are valid (Definition A.1). Thus $\tilde{x}^{CMPR}_{root} = 0$.*
2. *By irreversibility of the mutation process and heritability of missing data, in a valid $\tilde{x}$ if a leaf has state 0, all its ancestors must have state 0, therefore $\tilde{x}^{CMPR}_v = 0$.*
3. *By irreversibility of the mutation process and heritability of missing data, in a valid $\tilde{x}$ the ancestors of a leaf with state $s > 0$ can only have state $s$ or 0. Therefore, if $v$ has two descendent leaves with distinct positive states $s_1, s_2$, we must have $\tilde{x}_v = 0$ for all valid reconstructions $\tilde{x}$ of $x$. Therefore $\tilde{x}^{CMPR}_v = 0$.*
4. *The parsimony score of the subtree rooted at $v$ when all states in the subtree are assigned states of $-1$ is zero, and any other assignment has parsimony score of at least 2 (take a state $s \neq -1$ in the subtree and go down two distinct daughter lineages - some change to $-1$ will occur on both lineages). Therefore, in any VMPR $\tilde{x}$ of $x$ if the subtree rooted at $v$ does not have all states equal to $-1$, we can change them all to $-1$ and this will still yield a valid reconstruction, while lowering the parsimony score by at least 1, contradiction. Thus all VMPRs $\tilde{x}$ of $x$ have $\tilde{x}_v = -1$ and so $\tilde{x}^{CMPR}_v = -1$.*

5. *Consider the ancestor $g$ which satisfies the clause. We claim that for all VMPRs $\tilde{x}$ of $x$ we have $\tilde{x}_g = s$. If this is true, then $\tilde{x}_v$ cannot be $-1$ (since then by heritability of missing data $x_{\mathcal{L}(v)} = \{-1\}$), and $\tilde{x}_v$ cannot be $0$ nor some other state $s' \neq s$ (since the mutation process is irreversible). As a consequence, we would have that $\tilde{x}_v = s$ in all VMPRs $\tilde{x}$ of $x$, and so $\tilde{x}_v^{CMPR} = s$, as we want to show. Therefore, let us prove that for all VMPRs $\tilde{x}$ of $x$ we have $\tilde{x}_g = s$. Suppose otherwise to reach a contradiction. Let $\tilde{x}$ be a VMPR of $x$ with $\tilde{x}_g \neq s$. Then, the only option is that $\tilde{x}_g = 0$ (because the ancestors of a state $s$ can only be $0$ or $s$ by irreversibility of the mutation process and heritability of missing data). Now consider the subtree rooted at $g$, and consider the maximal connected component of $0$ states at and below $g$. Change all these states to $s$, and let $\tilde{x}'$ be the new states. We claim that $\tilde{x}'$ is still a valid reconstruction of $x$. Firstly, $\tilde{x}'$ is a reconstruction of $x$ since none of the leaf states have been perturbed, since $0 \notin x_{\mathcal{L}(g)}$ by the assumption that $\{s\} \subseteq x_{\mathcal{L}(g)} \subseteq \{-1, s\}$. Next, to check validity of $\tilde{x}$, note that by assumption $g$ is not the root, so the state of the root is still all zeros. As for transitions, we only need to check the transitions that were affected by swapping $0$ to $s$. Firstly, the edge going into $g$ was originally $0 \to 0$ and now it is $0 \to s$, which is still valid. Now, inspecting the transitions affected in the subtree rooted at $g$, we have that any $0 \to 0$ transitions that became $s \to s$ are still valid. Transitions $0 \to s$ become $s \to s$ which are valid, and similarly transitions $0 \to -1$ become $s \to -1$ which are valid. This accounts for all affected transitions. Thus, $\tilde{x}'$ is a valid reconstruction of $x$. Now let us analyze the parsimony score of $\tilde{x}'$. The transition at the root was $0 \to 0$ and now became $0 \to s$, so the parsimony score got worse by $1$. Among the remaining changes, only the transitions $0 \to s$ which became $s \to s$ change the parsimony score, and they each improve it by exactly $1$. We claim that there exist at least two such transitions. Indeed, by hypothesis, there are at least two leaves $l_1, l_2$ descending from $g$ on different daughter lineages through $u_1$ and $u_2$ which have a state of $s$. Consider the two distinct paths leading from $g$ to $l_1$ and $l_2$. In $\tilde{x}$, each of these paths must at some point transition from $0$ to $s$ (note that transitioning from $0$ to $-1$ is impossible because by heritability of missing data it would imply $x_{l_i} = -1$ for that leaf, a contradiction). Hence we obtain at least two new transitions $s \to s$ - one on each path - which jointly improve the parsimony score by at least $2$. Hence Par$(\tilde{x}') \leqslant$ Par$(\tilde{x}) - 1$. This contradicts the assumption that $\tilde{x}$ was VMPR, and so we are done.*

6. *It suffices to show that in this case there exist two different VMPRs $\tilde{x}, \tilde{x}'$ of $x$ with $\tilde{x}_v = s$ and $\tilde{x}'_v = 0$. We construct these explicitly. For $\tilde{x}$ we choose $\tilde{x}^{VMPR}$. By Proposition A.1, we have that $\tilde{x}^{VMPR} = s$. To construct a VMPR $\tilde{x}'$ with $\tilde{x}'_v = 0$, start from $\tilde{x}^{VMPR}$. Travel up the ancestors of $v$ until we find the last ancestor $g$ such that $\tilde{x}_g^{VMPR} = s$. Let $v = a_1, a_2, \ldots, a_j = g$ be the sequence of ancestors visited. Note that $g$ cannot be the root node since $\tilde{x}^{VMPR}$ is valid. If $p$ is the parent of $g$, then the only possibility is $\tilde{x}_p^{VMPR} = 0$ (by irreversibility of the mutation process and heritability of missing data). For every $a_i$, the fact that $\tilde{x}_g^{VMPR} = s$ means that the states that descend from $a_i$ are either $-1$ or $s$. However, by condition 6 only one of the daughter subtrees of $a_i$ can contain state $s$, so that the other one must consist fully of $-1$. The characterization of $\tilde{x}^{VMPR}$ further implies that all these subtrees that have missing states at all leaves, have all internal nodes with state $-1$ too. This way, consider the following modification to $\tilde{x}^{VMPR}$: change the state of all nodes $u = a_1, a_2, \ldots, a_j = g$ from $s$ to $0$. This is still a valid reconstruction of $x$. The transition into $g$ from $p$ changes from $0 \to s$ to $0 \to 0$, so we win a parsimony score of $1$. All the transitions $s \to -1$ from node $a_i$ into the subtrees with missing states at the leaves become $0 \to -1$, so the parsimony remains unchanged by these transition changes. All that remains is the transition from $v$ into its unique subtree which contains a leaf with state $s$. This transition must originally have been $s \to s$ (since otherwise missing data is heritable), so that in $\tilde{x}'$ the transition is now $0 \to s$ and so we lose a parsimony score of $1$. The net change in the parsimony score from $\tilde{x}$ to $\tilde{x}'$ is thus zero, and so Par$(\tilde{x}') =$ Par$(\tilde{x})$, completing the proof.*

□

Figure A.6 shows a larger example of CMPR, highlighting the subtlety of conditions 5 and 6 of the Theorem. As a remark, when there is no missing data, condition 6 of Theorem A.3 can never hold, implying that the CMPR will reconstruct all ancestral states. In particular, this means that the VMPR is unique when there is no missing data. Thus, CMP only makes a difference when there is missing data.

The characterization of the CMPR given by Theorem A.3 allows us to implement it easily in time $\mathcal{O}(n)$ where $n$ is the number of nodes in the tree (i.e., the same time complexity as Fitch-Hartigan): we can just take the imputations given by Algorithm 1 and after the fact set to `NONE` all nodes that satisfy condition 6 of the CMPR Theorem A.3. To determine which nodes satisfy this condition, we can use a bottom-up tree traversal to determine which nodes $v$ satisfy condition 5 with $g = v$, and then propagate this information down with a top-down tree traversal to determine which nodes $v$ satisfy condition 5 for any ancestor $g$. Any nodes that satisfy the original condition 5 of the VMPR Proposition A.1 but not condition 5 of the CMPR Theorem A.3 are exactly those with ambiguous states that need to be set to `NONE`. We provide the algorithm below in Algorithm 2:

Armed with the characterization of the CMPR $\tilde{x}^{\text{CMPR}}$ of $x$ provided by Theorem A.3, we are ready to show that the likelihood $p_\theta(\tilde{Z}^{(1)} = \tilde{x}^{(1)})$ where $\tilde{x} = \tilde{x}^{\text{CMPR}}$ is tractable; in $\tilde{x}^{\text{CMPR}}$, states that are not reconstructed and thus marked as `NONE` are without loss of generality replaced by $-1$ such that they are marginalized out as in an ignorable missing data mechanism:

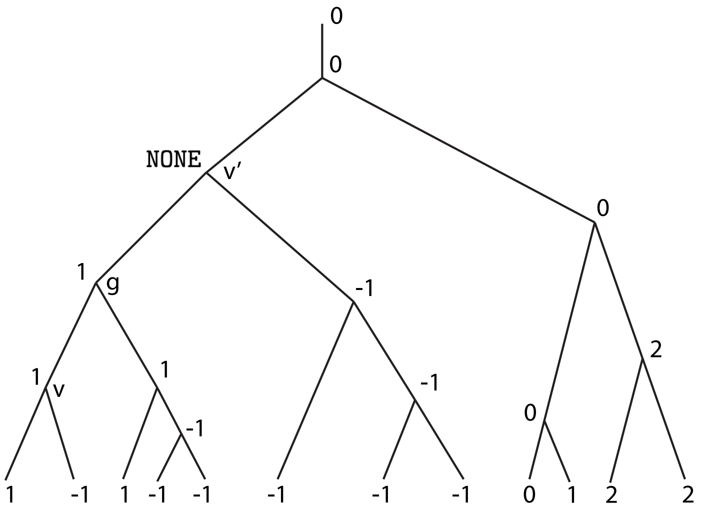


Fig. A.6. A larger example of conservative maximum parsimony reconstruction (CMPR). Node $v$ is reconstructed with state 1 because it satisfies condition 5 of Theorem A.3 via the node labeled $g$. In contrast, node $v'$ is not reconstructed (represented with the symbol NONE) since it instead satisfies condition 6 of Theorem A.3. In particular, this means that CMPR is mapping the mutation to state 1 present at node $g$ to either the edge connecting $g$ to $v'$, or the edge connecting $v'$ to its parent – without making an arbitrary choice. Instead, naive maximum parsimony would have committed the mutation to one of these two edges. For example, the VMPR Algorithm 1 would have assigned $v'$ the state 1, meaning it would have mapped the mutation to the edge connecting $v'$ to its parent. In this example, the CMPR and the VMPR from Algorithm 1 agree on all other nodes.

**Algorithm 2 Conservative Maximum Parsimony Reconstruction**

**procedure** CMPR($\mathcal{T}$, $x$)
  $\tilde{x} \leftarrow$ VMPR$(\mathcal{T}, x)$
  **for** $v$ in preorder traversal of internal nodes of $\mathcal{T}$ **do**
    **if** $v$ is the root, or $0 \in x_{\mathcal{L}(v)}$, or $x_{\mathcal{L}(v)}$ contains two distinct positive states, or $x_{\mathcal{L}(v)} = \{-1\}$ **then**
      VMPR_cond_5[$v$] ← False
      CMPR_cond_5[$v$] ← False
    **else**
      VMPR_cond_5[$v$] ← True
      Let $s$ be the unique positive state in $x_{\mathcal{L}(v)}$
      **if** $|\{u : u \text{ child of } v, s \in x_{\mathcal{L}(u)}\}| \geqslant 2$ **then**
        CMPR_cond_5[$v$] ← True
      **else**
        $p \leftarrow$ parent$(v)$
        CMPR_cond_5[$v$] ← CMPR_cond_5[$p$]
  **for** $v$ in the internal nodes of $\mathcal{T}$ **do**
    **if** VMPR_cond_5[$v$] is True and CMPR_cond_5[$v$] is False **then**
      $\tilde{x}_v \leftarrow$ NONE
  **return** $\tilde{x}$

THEOREM A.4 ($\tilde{x}^{\text{CMPR}}$ has no unobserved confounders). *Let $\tilde{x} = \tilde{x}^{CMPR}$ be the CMPR of the leaf states $x$ of one character, with NONE replaced by $-1$. For a node $v$, denote by $gpa(v)$ it closest reconstructed, non-missing ancestor in $\tilde{x}$ (not including $v$), that is to say its closest ancestor $u$ with $\tilde{x}_u \neq -1$. Then:*

$$p_\theta(\tilde{Z}^{(1)} = \tilde{x}^{(1)}) = \prod_{v \in V, u = gpa(v)\,:\, \tilde{x}_v \neq -1} p_\theta(\tilde{Z}_v = \tilde{x}_v | \tilde{Z}_u = \tilde{x}_u). \tag{A.4}$$

*Proof.* *It suffices to show that there are no unobserved confounders, that is to say, that if a node $v$ has $\tilde{x}_v = -1$, then all subtrees of $v$ except possibly one are fully missing. Indeed, in such case, we can repeatedly prune away all these missing subtrees and the missing node (replacing two edges by one) without changing the likelihood, leaving us with a tree where **all** nodes are observed. In this new tree, each remaining node $v$ is connected to $u = gpa(v)$. We would thus obtain the factorization* *in Eq. A.4. To prove that if a node $v$ has $\tilde{x}_v = -1$ then all subtrees of $v$ except possibly one are fully missing, we proceed by contradiction, supposing otherwise. Then $v$ (which is not the root) has at least two distinct subtrees with at least one observed leaf. Let $s_1$ and $s_2$ be the states of two such leaves. If we have $s_1 \neq s_2$, then implication 3 of Theorem A.3 implies that $\tilde{x}_v = 0$, contradiction. Therefore, the only non-missing state in these subtrees is a single state $s$, and now implication 5 of Theorem A.3 implies that $\tilde{x}_v = s$, again a contradiction, so we are done, since each observed state is conditionally independent from its observed non-descendants given its closest observed ancestor, giving the decomposition of Eq.* (A.4). □

Finally, we should note that the Sankoff algorithm can be used to compute the CMPR in $\mathcal{O}(nk^2)$ time where $k$ is the number of states, and in fact in time $\mathcal{O}(nk)$ by leveraging irreversibility, and moreover in time $\mathcal{O}(n)$ by first precomputing the valid set of states based on states below (which can only be $-1, 0$, and one other state). We used this for testing our implementation, but note that the combinatorial characterization of the CMPR is not elucidated by such algorithm, and thus it does not explain why line 11 is true.

### Tree Simulation Details

Trees were simulated using a subsampled birth-death process. In this simulation, each cell has a birth rate and a death rate. The amount of time before a birth or a death event occurs follows an exponential distribution with the given birth and death rates. Additionally, a cell cannot divide before at least $0.01$ units of time have passed, called the *offset*. Initial birth and death rates are set such that birth rate is ten times higher than death rate and such that under a birth-death process with these rates and offset, the expected population size after $1$ unit of time is $40000$. This yields a birth rate of $15.75$ and a death rate of $1.575$. Whenever a cell divides, its fitness changes with probability $6.4\%$. When such a change in fitness occurs, $90\%$ of the time the birth rate becomes lower by multiplying it by $0.93$. The remaining $10\%$ of the time the birth rate becomes higher by multiplying it by $2.14$. Once the cell population reaches $40000$, we terminate the simulation and sample $400$ leaves uniformly at random, to match a sampling probability of $1\%$. The chronogram induced by these $400$ cells is then the ground truth chronogram used in the simulations. The fitness parameters described above were chosen to obtain trees that showcased interesting fitness variation, as displayed in Supplementary Figure 4.

For the smaller trees with just $40$ leaves, birth and death rates are set such that birth rate is ten times higher than death rate and such that under a birth-death process with these rates and offset, the expected population size after $1$ unit of time is $4,000$. We simulate without changes in fitness, since this is a very small population. Once the cell population reaches $4,000$, we terminate the simulation and sample $40$ leaves uniformly at random, to match a sampling probability of $1\%$. The first $9$ of the $50$ trees are shown in Figure A.7.

For the larger trees with $2,000$ leaves, initial birth and death rates are set such that birth rate is ten times higher than death rate and such that under a birth-death process with these rates and offset, the expected population size after $1$ unit of time is $20,000$. Whenever a cell divides, its fitness changes with probability $4.8\%$. When such a change in fitness occurs, $90\%$ of the time the birth rate becomes lower by multiplying it by $0.93$. The remaining $10\%$ of the time the birth rate becomes higher by multiplying it by $2.14$. Once the cell population reaches $20,000$, we terminate the simulation and sample $2000$ leaves uniformly at random, to match a sampling probability of $10\%$. We note that simulating $200,000$ cells and subsampling $2,000$ at a probability of $1\%$ would be computational too slow, which is why we subsample at a probability of $10\%$ instead. The first $9$ of the $50$ trees are shown in Figure A.8.

### LAML Command Line

We ran LAML using the following command line:

```
run_laml -c character-matrix.csv -t tree.nwk -o laml --nInitials 1 --maxIters 0
```

Here, '-c' specified the character matrix, '-t' specifies the initial tree topology, '-o' specifies the output location, '–nInitials' specifies the number of initial points, and '–maxIters' specifies the number of topology search steps. Using '–maxIters 0' ensures that the initial topology is not modified, while '–nInitials 1' maximizes the speed of the method; we observed no accuracy improvements by using a larger value of 'nInitials' on the intMEMOIR dataset, while runtime scales linearly in 'nInitials', which is why we set '–nInitials 1'.

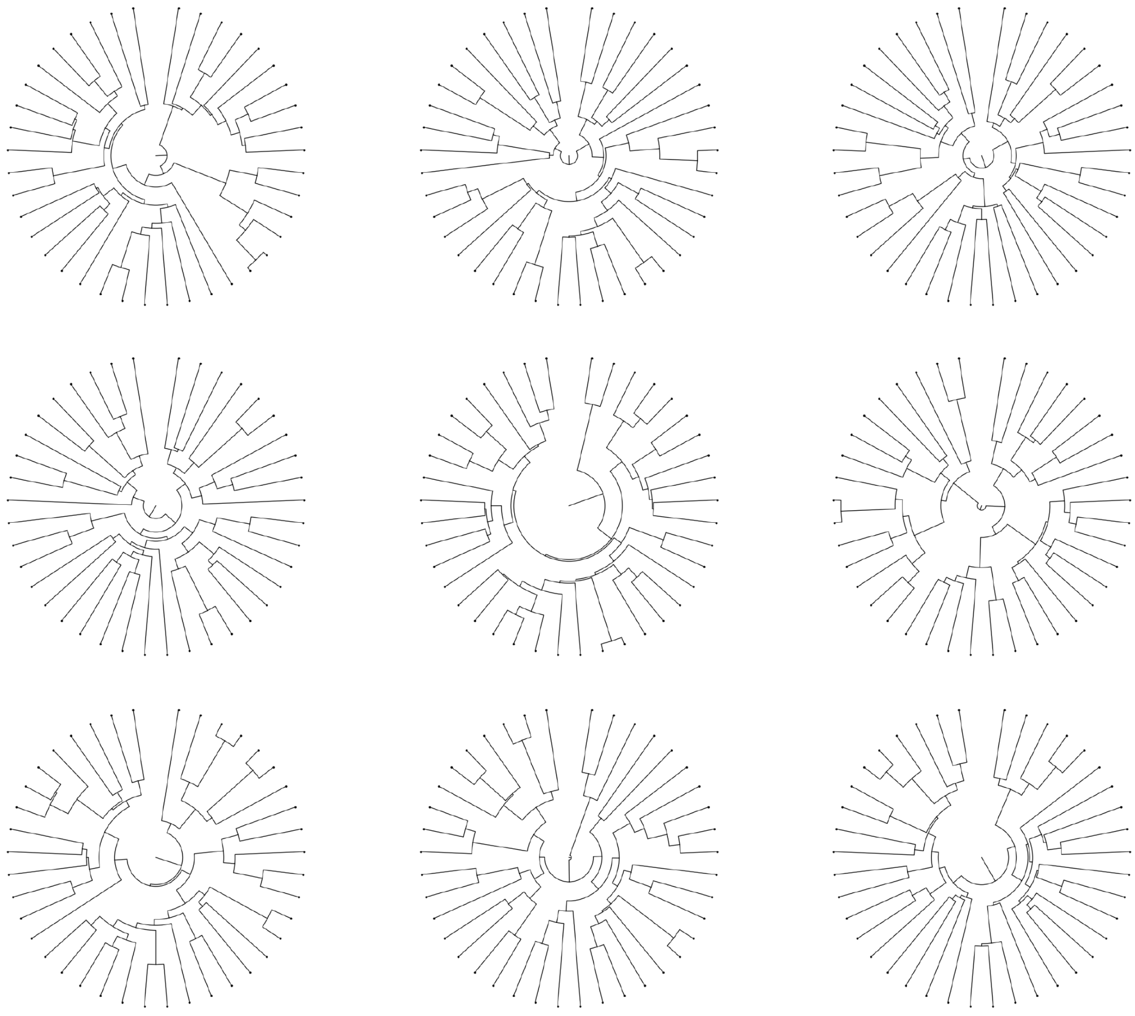

Fig. A.7. **Sample ground truth trees with** 40 **leaves**. Ground truth trees corresponding to the first 9 random seeds.

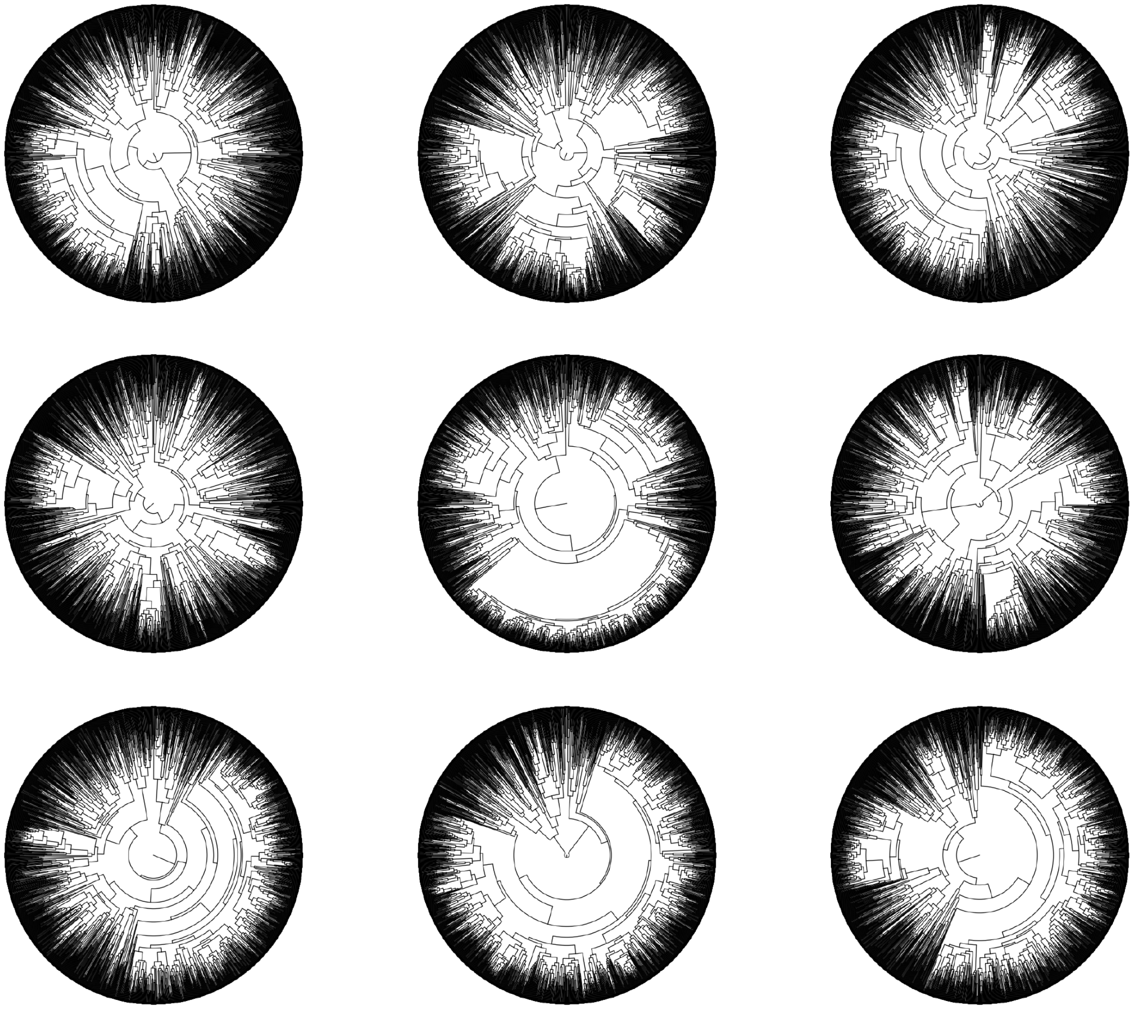

Fig. A.8. **Sample ground truth trees with** 2000 **leaves**. Ground truth trees corresponding to the first 9 random seeds.

### SUPPLEMENTARY FIGURES

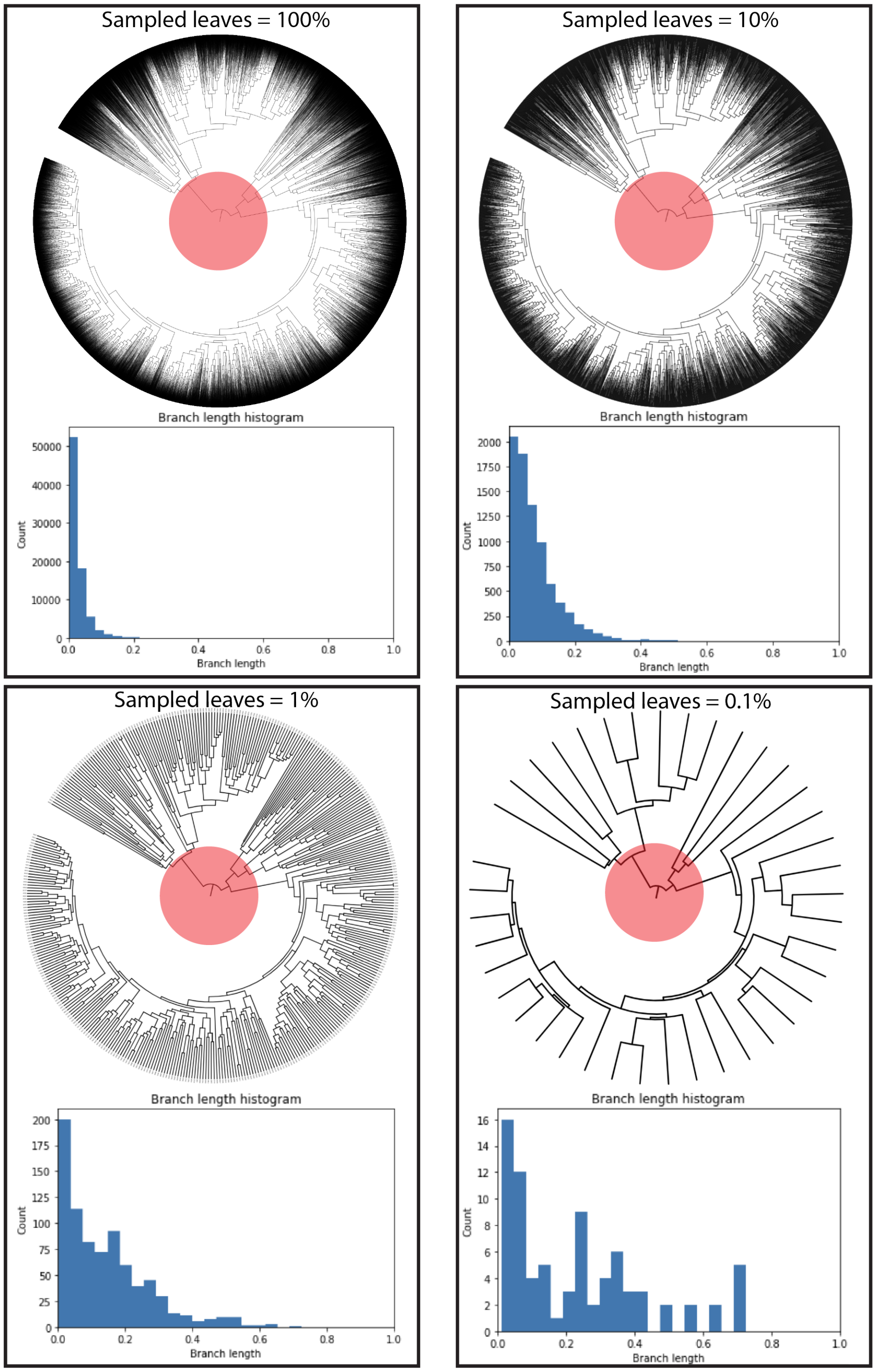


Supplementary Fig. 1. **The effect of sampling leaves on branch lengths**. Sampling leaves causes short edges at the bottom of the tree to get pruned away first and replaced with longer edges. This dramatically changes the branch length distribution. Note how the top of the tree (highlighted) is perfectly preserved even when sampling as few as 0.1% of the leaves in the original tree, while the bottom of the tree is significantly remodelled. The effect of sampling must thus be carefully considered when interpreting single-cell chronograms, developing new methods, and designing regularization schemes.

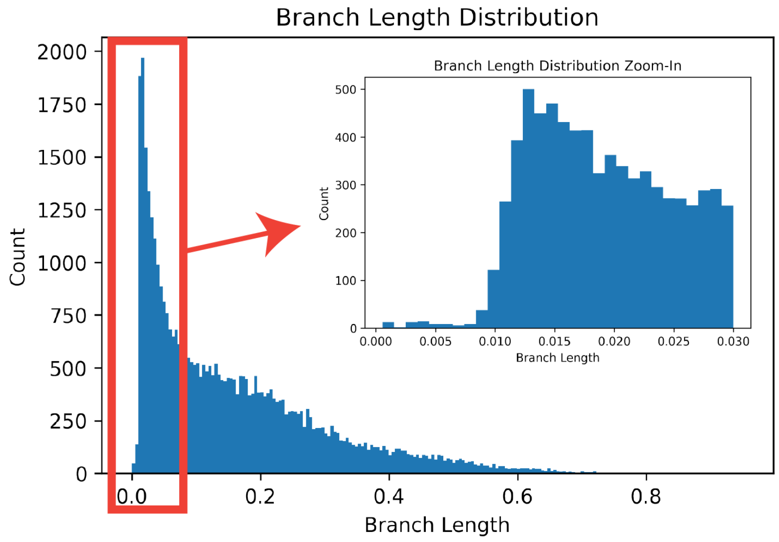


Supplementary Fig. 2. **Distribution of branch lengths of the simulated ground truth trees**. A value of $\epsilon = 0.01$ is a reasonable lower bound on branch lengths in our simulated trees.

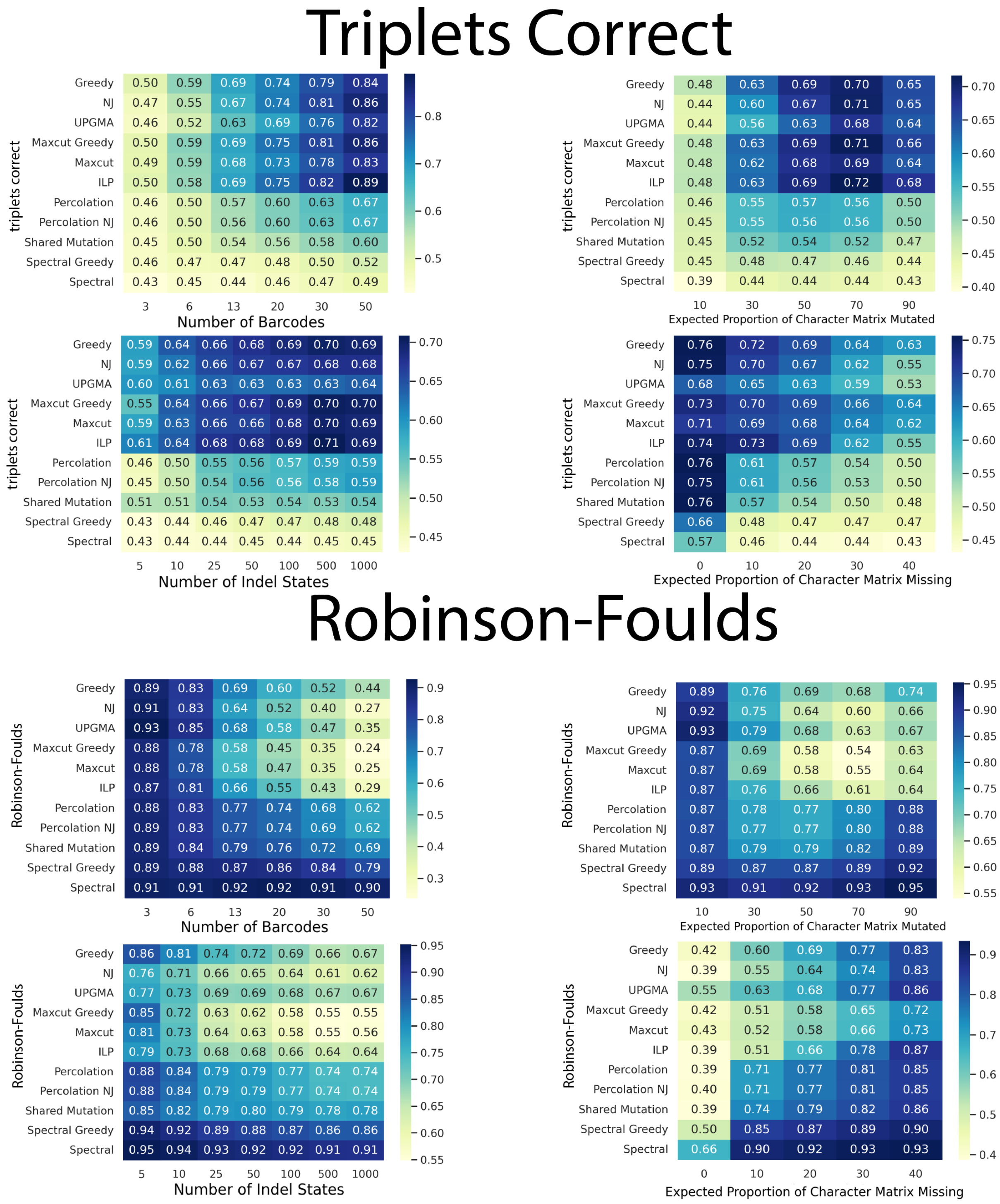


Supplementary Fig. 3. **Selection of topology estimation method**. We evaluated a collection of topology estimation algorithms (called 'solvers') from the Cassiopeia package to determine which one to use in our analysis of branch length estimators. We used the Triplets Correct and Robinson-Foulds metrics to select the algorithm. Note that these metrics do not require branch lengths, only topologies. We find that for both metrics, the Maxcut solver is among the best performers while also being much faster than the ILP solver, so we use it in the remainder of this work.

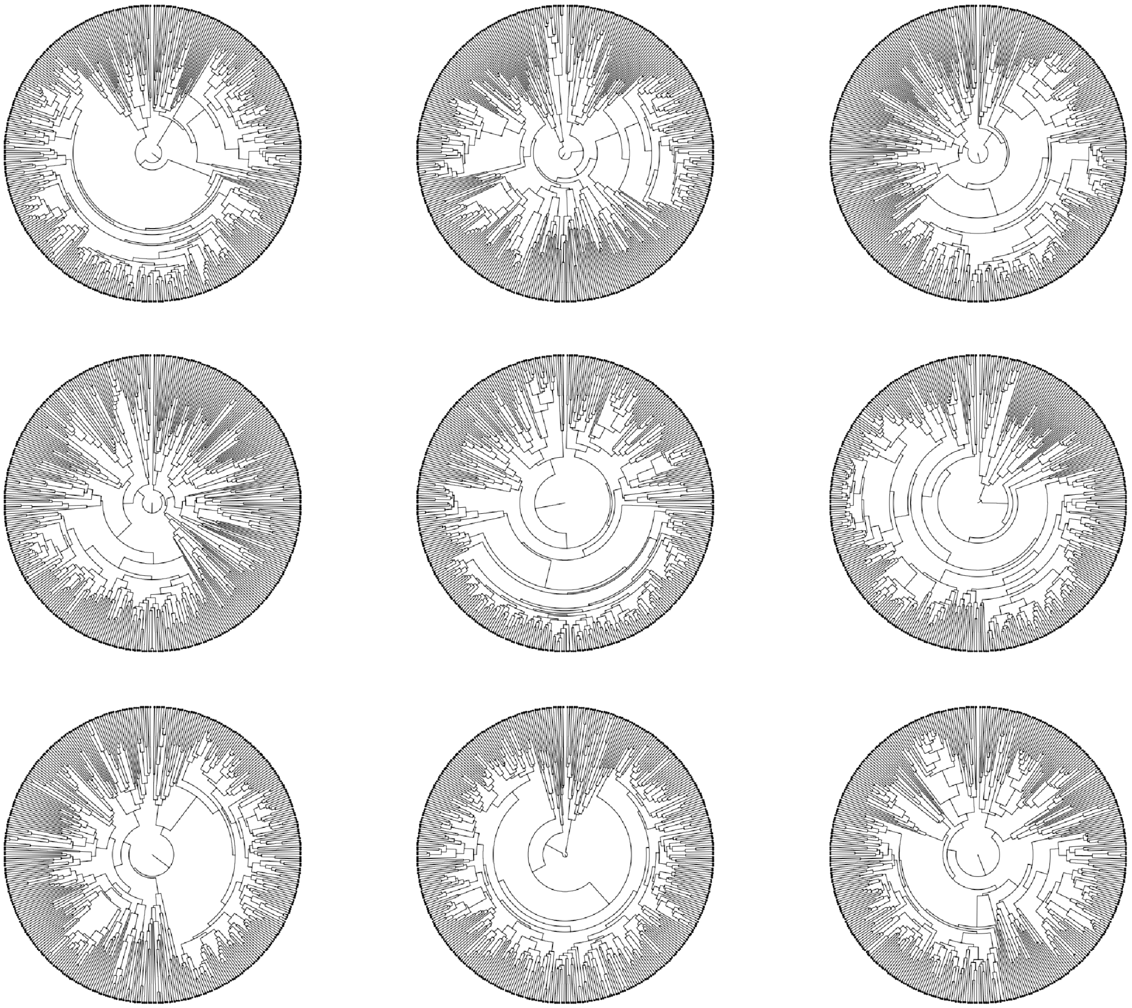

Supplementary Fig. 4. **Sample ground truth trees**. Ground truth trees corresponding to the first 9 random seeds. Our simulated trees are diverse and showcase subclones with different proliferative capacity.

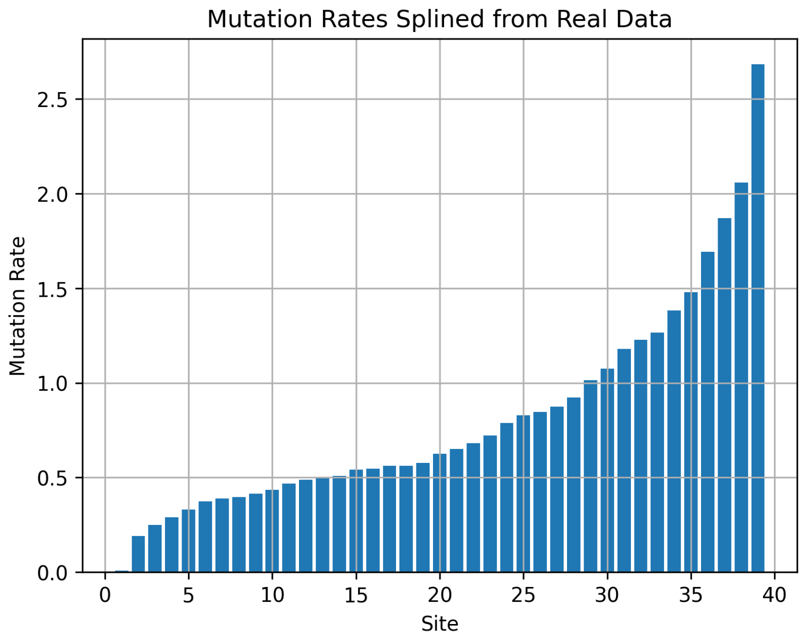


Supplementary Fig. 5. **(Mutation rates)** For our simulations, we use different mutation rates for different sites. These site-specific mutation rates were splined from real data estimates and exhibit a broad range of values.

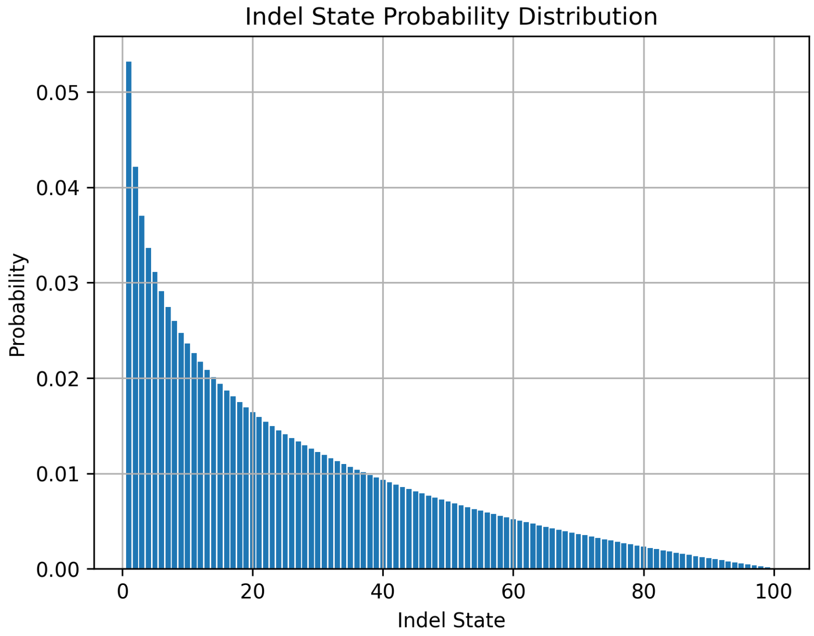


Supplementary Fig. 6. **(Indel Probabilities)** For our simulations, upon CRISPR/Cas9 cutting we introduce indels with different probabilities. Some indels are introduced with higher probability than others. This is an important feature of real data which leads to high degrees of *homoplasy*, wherein common indels are introduced independently in different cell lineages, complicating topology reconstruction and branch length estimation.

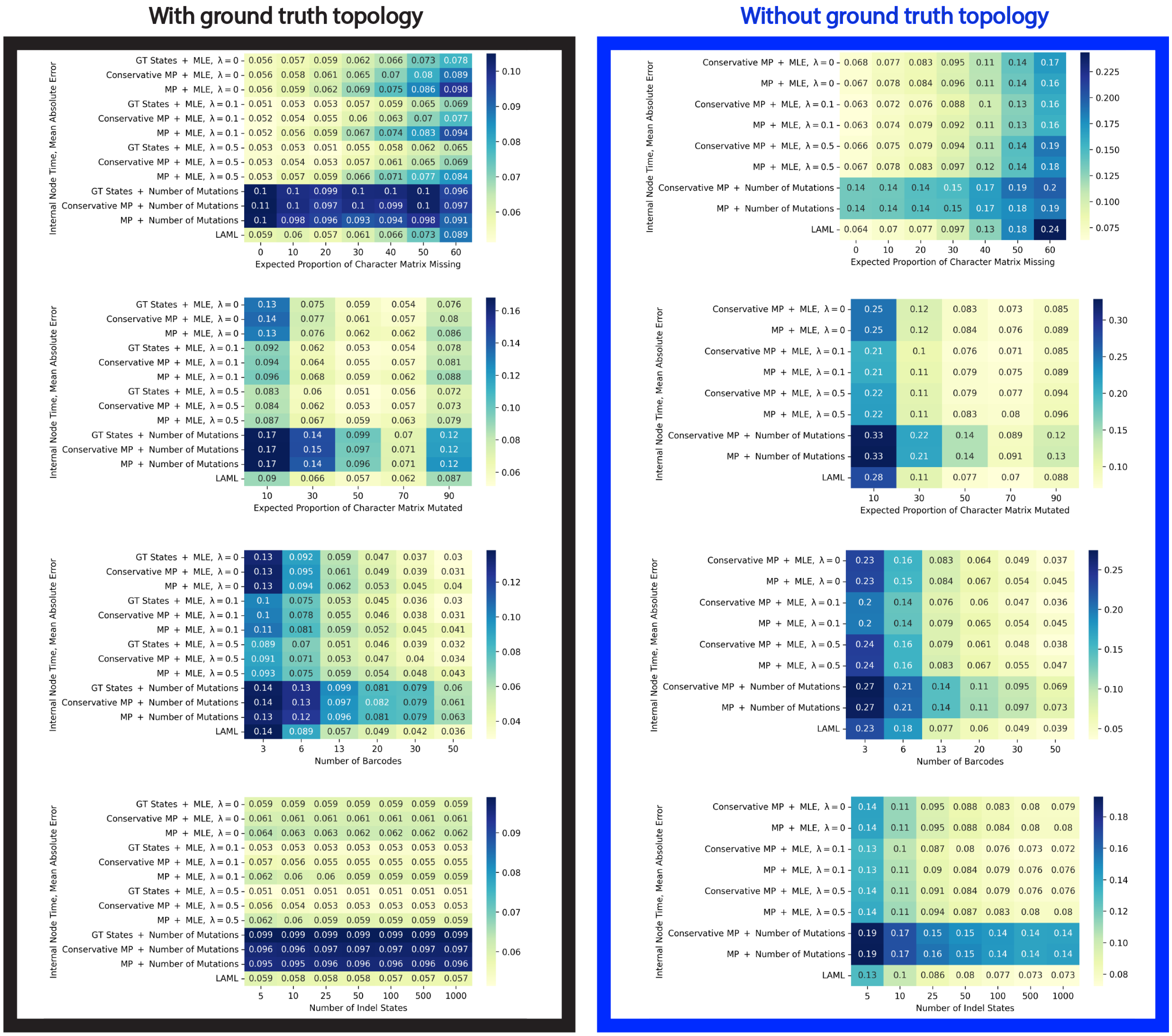


Supplementary Fig. 7. **Performance of branch length estimation methods under different parameter regimes for the “internal node time” task**. On the left we show performance when the ground truth topology is known, and on the right we show the performance when the topology is not know and must be reconstructed – as is the case in all real-life applications. In this latter case, the Maxcut algorithm from the Cassiopeia package is used. Each number displayed is the average over the 50 simulated trees for the given parameter regime.

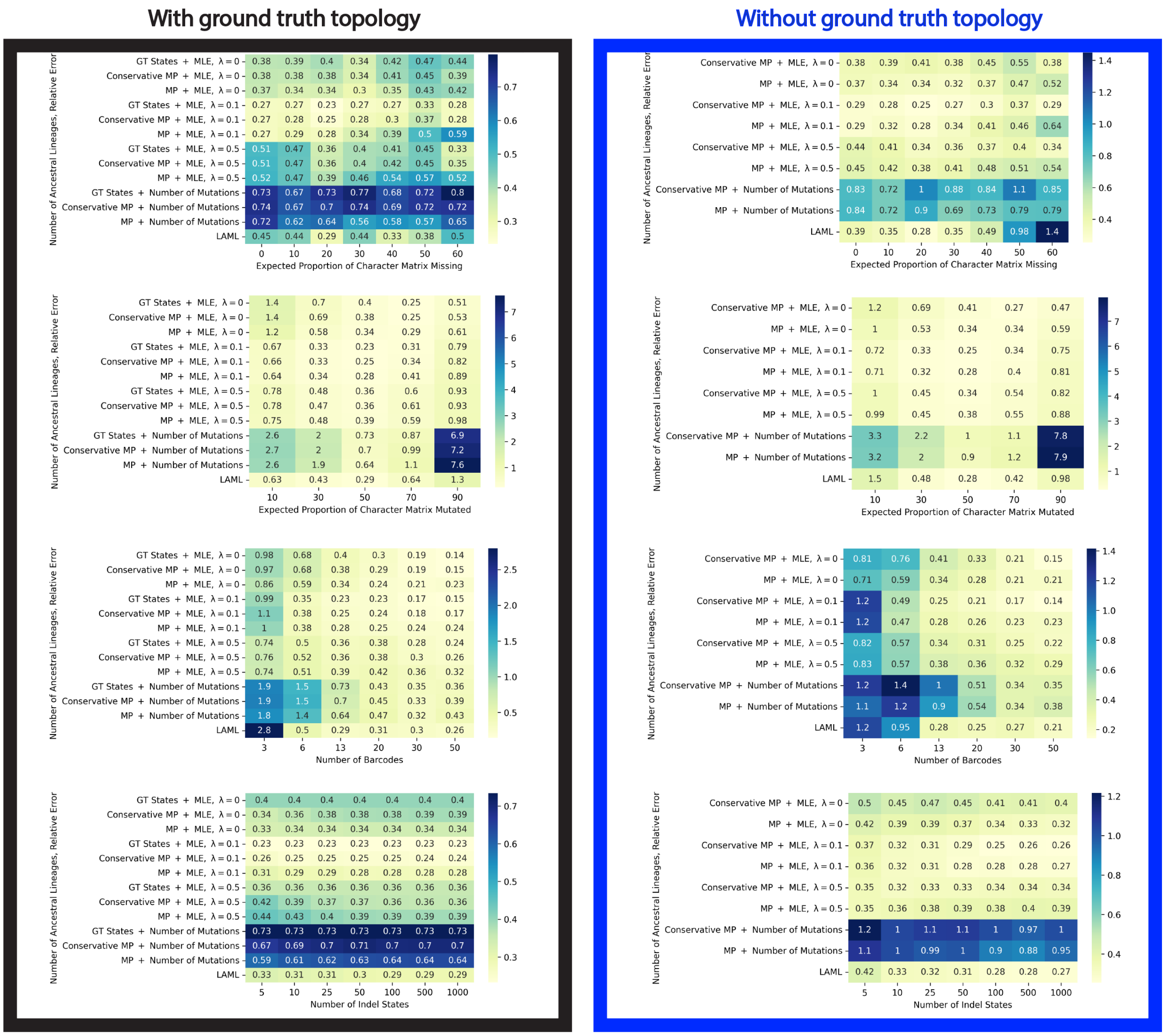


Supplementary Fig. 8. **Performance of branch length estimation methods under different parameter regimes for the "ancestral lineages" task**. On the left we show performance when the ground truth topology is known, and on the right we show the performance when the topology is not know and must be reconstructed – as is the case in all real-life applications. In this latter case, the Maxcut algorithm from the Cassiopeia package is used. Each number displayed is the average over the 50 simulated trees for the given parameter regime.

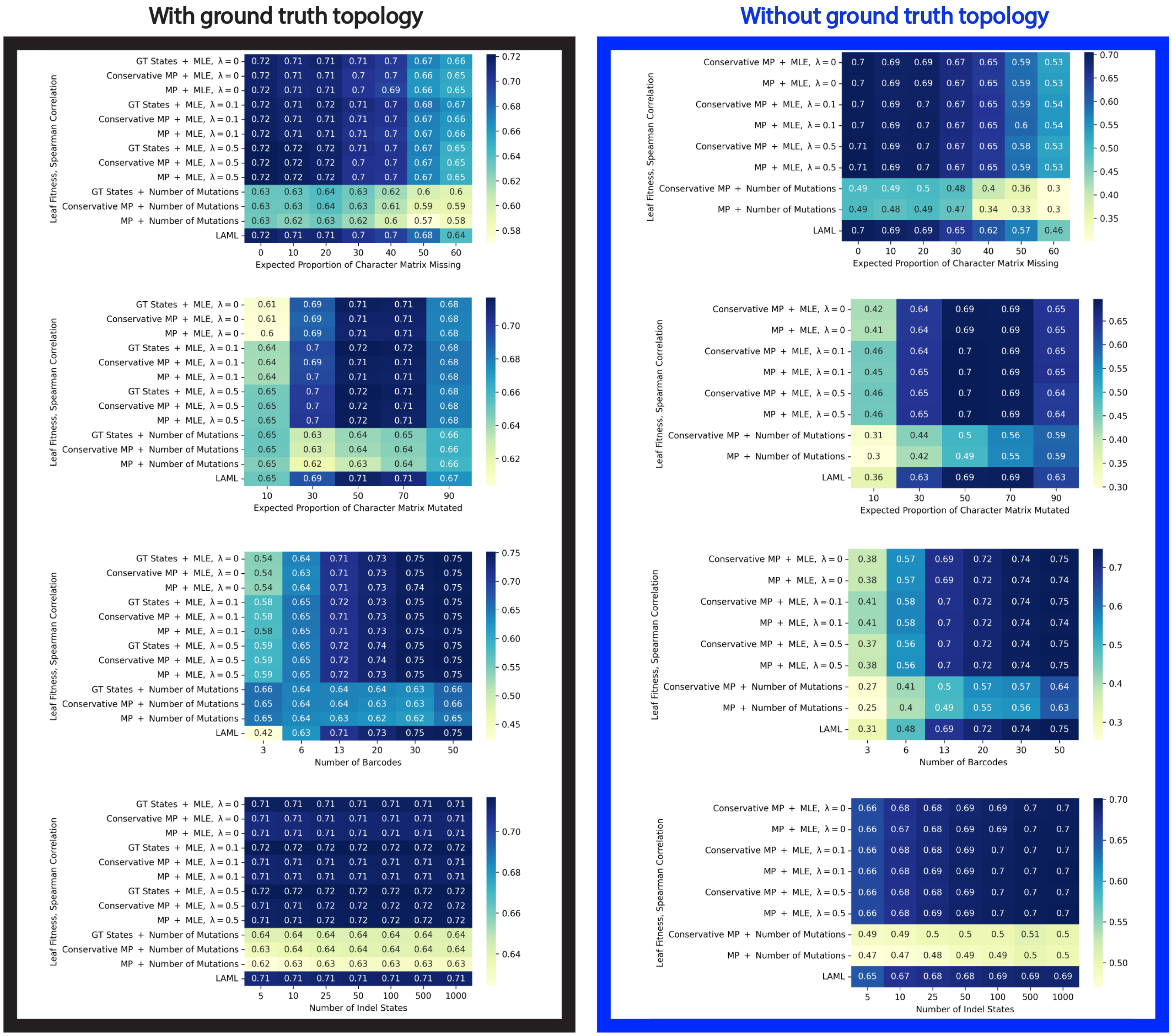


Supplementary Fig. 9. **Performance of branch length estimation methods under different parameter regimes for the "fitness inference" task**. On the left we show performance when the ground truth topology is known, and on the right we show the performance when the topology is not know and must be reconstructed – as is the case in all real-life applications. In this latter case, the Maxcut algorithm from the Cassiopeia package is used. Each number displayed is the average over the 50 simulated trees for the given parameter regime.

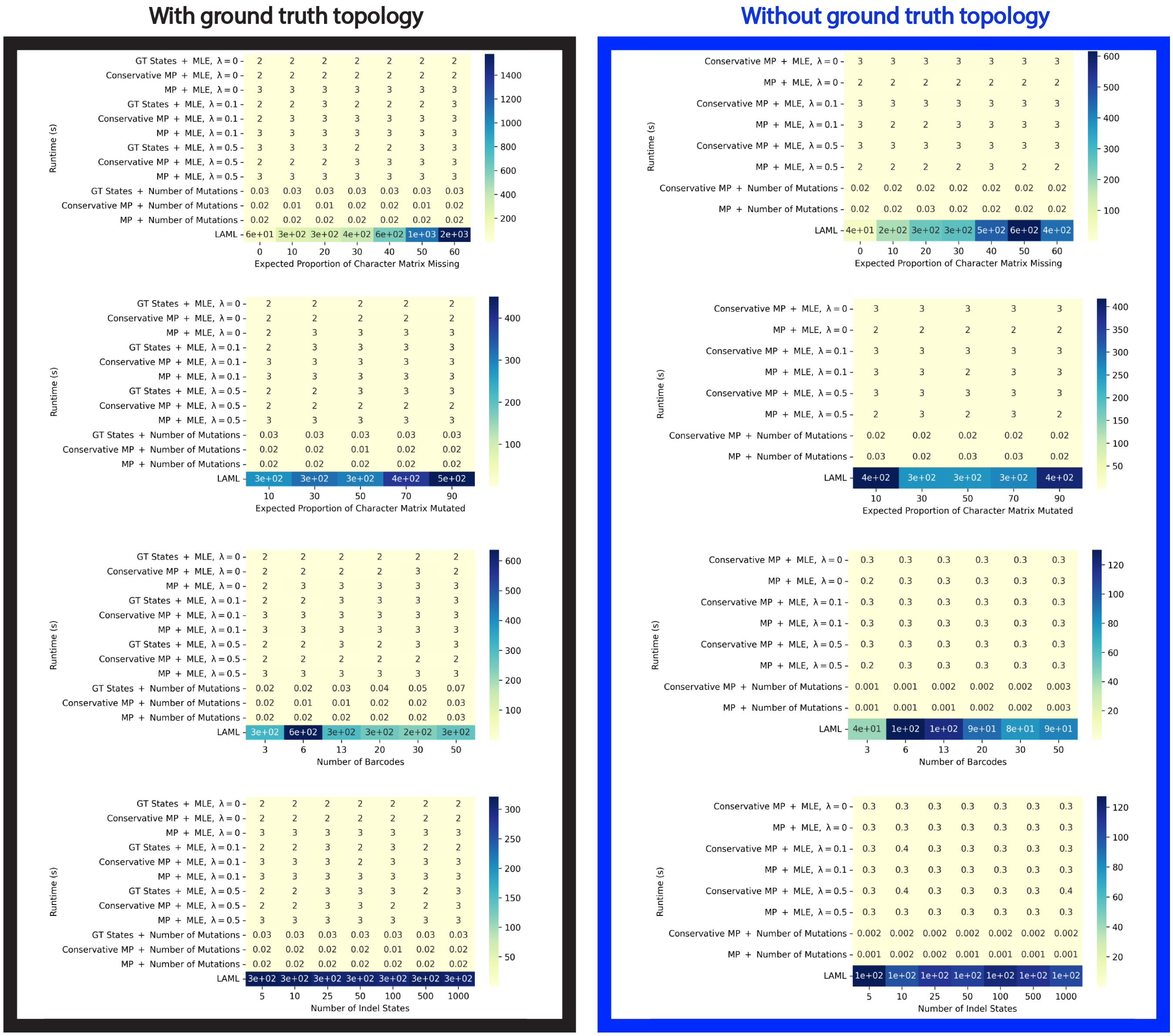


Supplementary Fig. 10. **Runtime of branch length estimation methods under different parameter regimes**. On the left we show performance when the ground truth topology is known, and on the right we show the performance when the topology is not know and must be reconstructed – as is the case in all real-life applications. In this latter case, the Maxcut algorithm from the Cassiopeia package is used. Each number displayed is the average over the 50 simulated trees for the given parameter regime. Runtime is shown in seconds, with one significant digit.

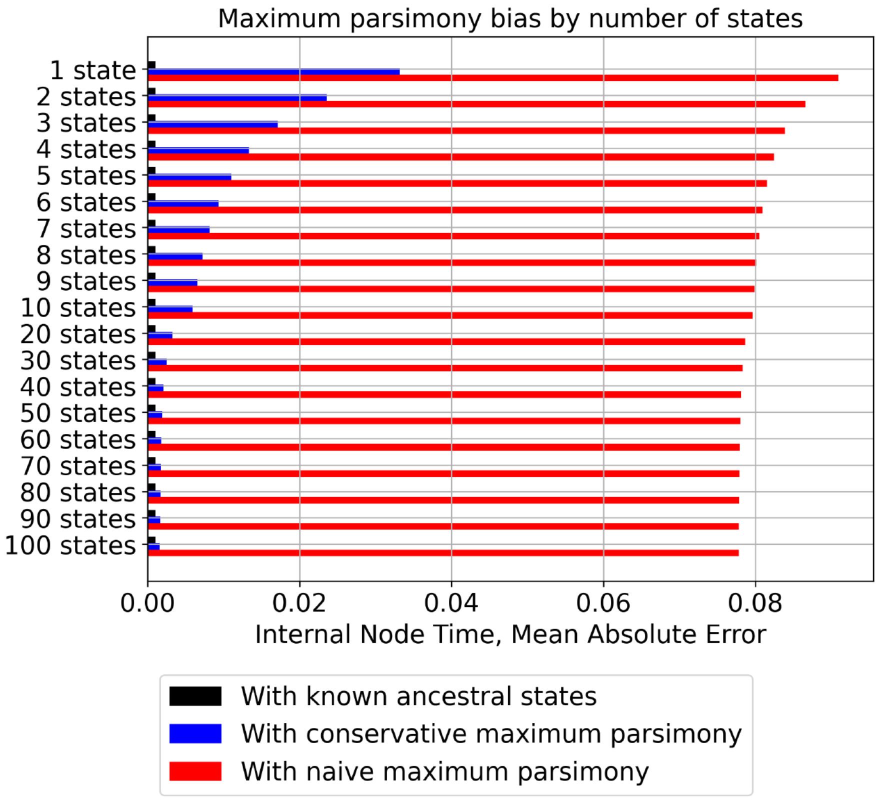


Supplementary Fig. 11. **Conservative maximum parsimony enables negligible bias, even when there is abundant sequencing dropouts and no epigenetic silencing**. We repeated the experiment from Figure 3 but with 60% sequencing dropouts and 0% epigenetic silencing. Supporting our theoretical arguments, the results show negligible bias from using conservative maximum parsimony reconstruction (CMPR).

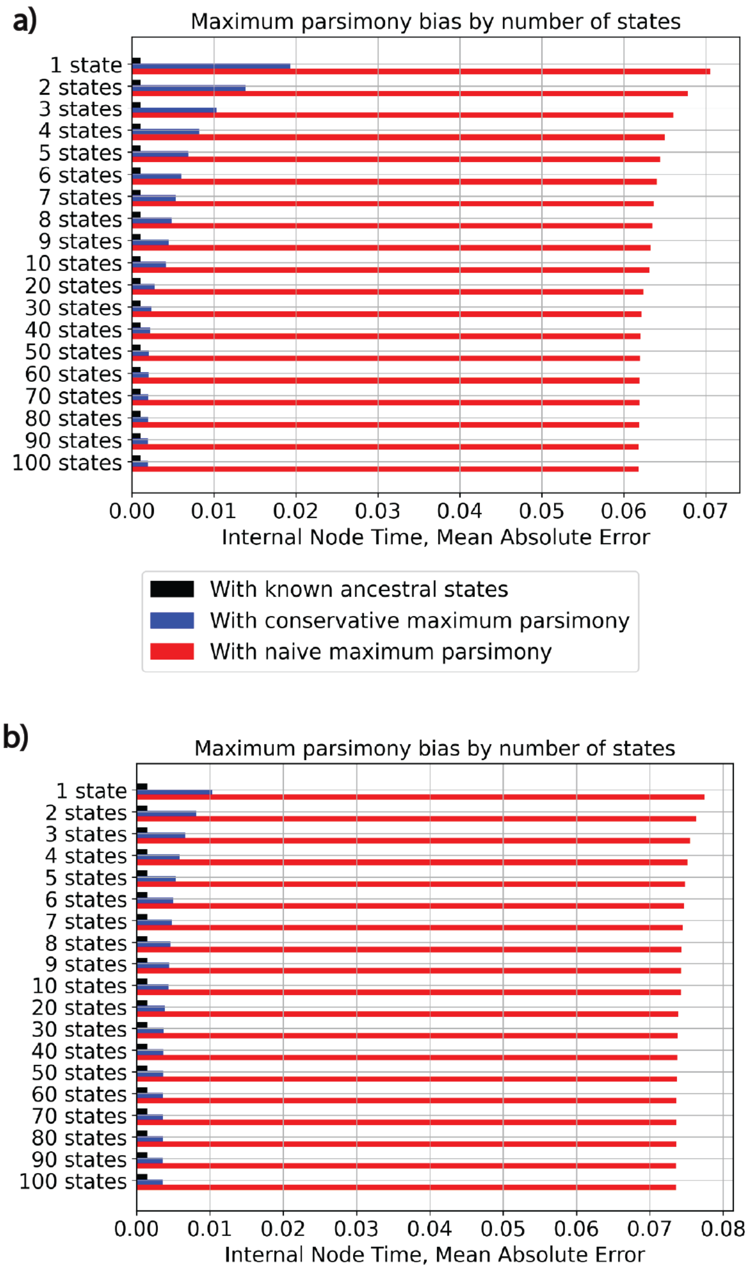


Supplementary Fig. 12. **Conservative maximum parsimony enables negligible bias even in the presence of double resections**. a) We repeated the experiment from Figure 3 but included double-resection events. This introduces minimal bias for large state spaces, and improves results on small state spaces due to the indirectly increased number of states. b) We repeated the same experiment as part a but with a large barcode size of 10 while keeping the number of sites fixed at 100,000. CMP still shows little bias for large state spaces, showing the effectiveness of our method even in the light of double-resection misspecification.

a)

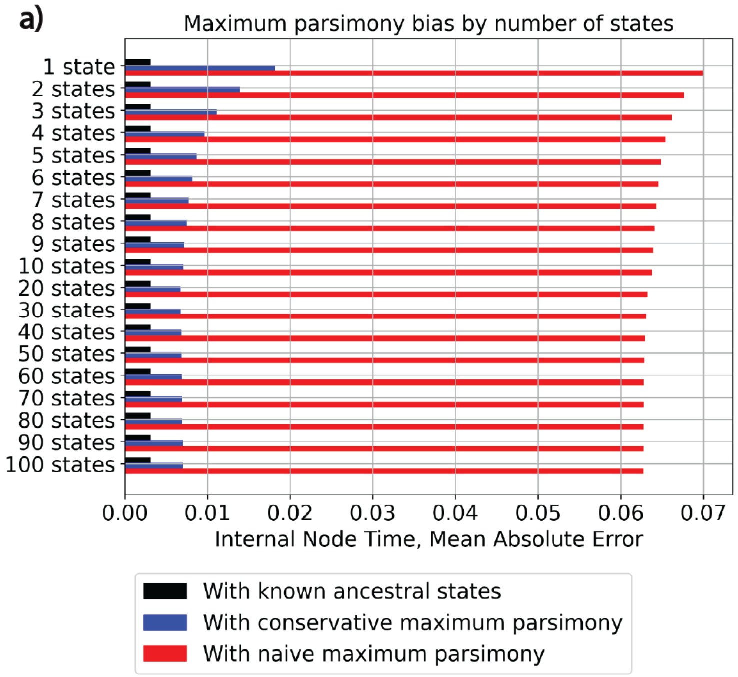


b)

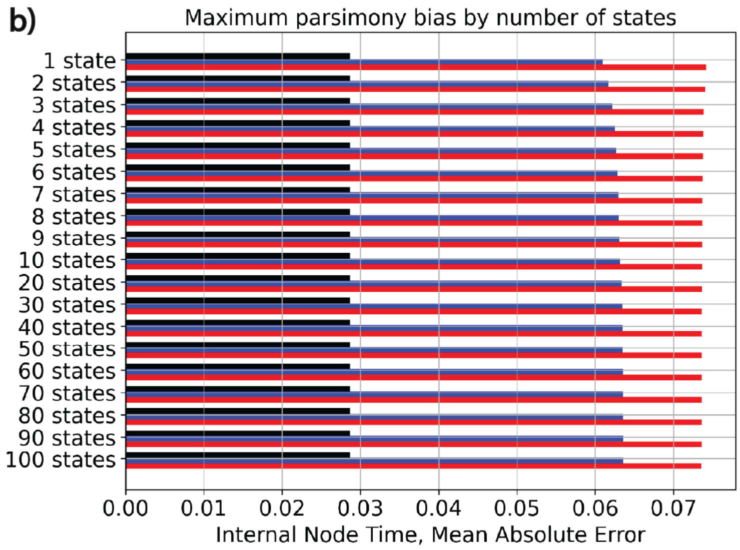


Supplementary Fig. 13. **Naive representation of double-resection events introduces bias**. a) We repeated the experiment from Supplementary Figure 19a but naively encoded double-resection events. This introduces noticeable bias even when the ground truth states are known. b) We repeated the experiment from Supplementary Figure 19b but naively encoded double-resection events. This introduces massive bias even when the ground truth states are known. This shows the importance of our choice of double-resection encoding.

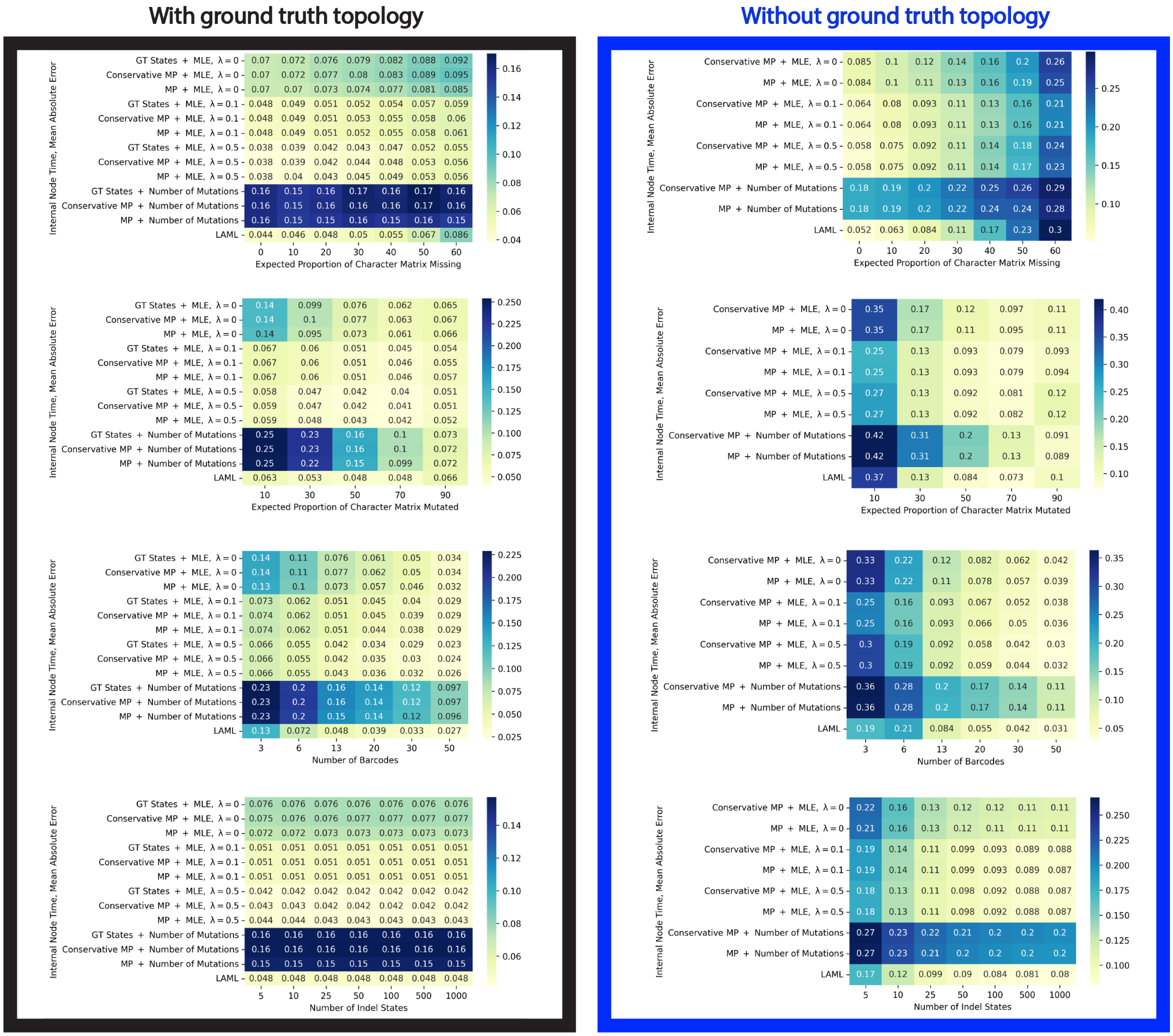


Supplementary Fig. 14. **Performance of branch length estimation methods under different parameter regimes for the “internal node time” task, on larger trees with** 2000 **leaves**. On the left we show performance when the ground truth topology is known, and on the right we show the performance when the topology is not know and must be reconstructed – as is the case in all real-life applications. In this latter case, the Maxcut algorithm from the Cassiopeia package is used. Each number displayed is the average over the 50 simulated trees for the given parameter regime.

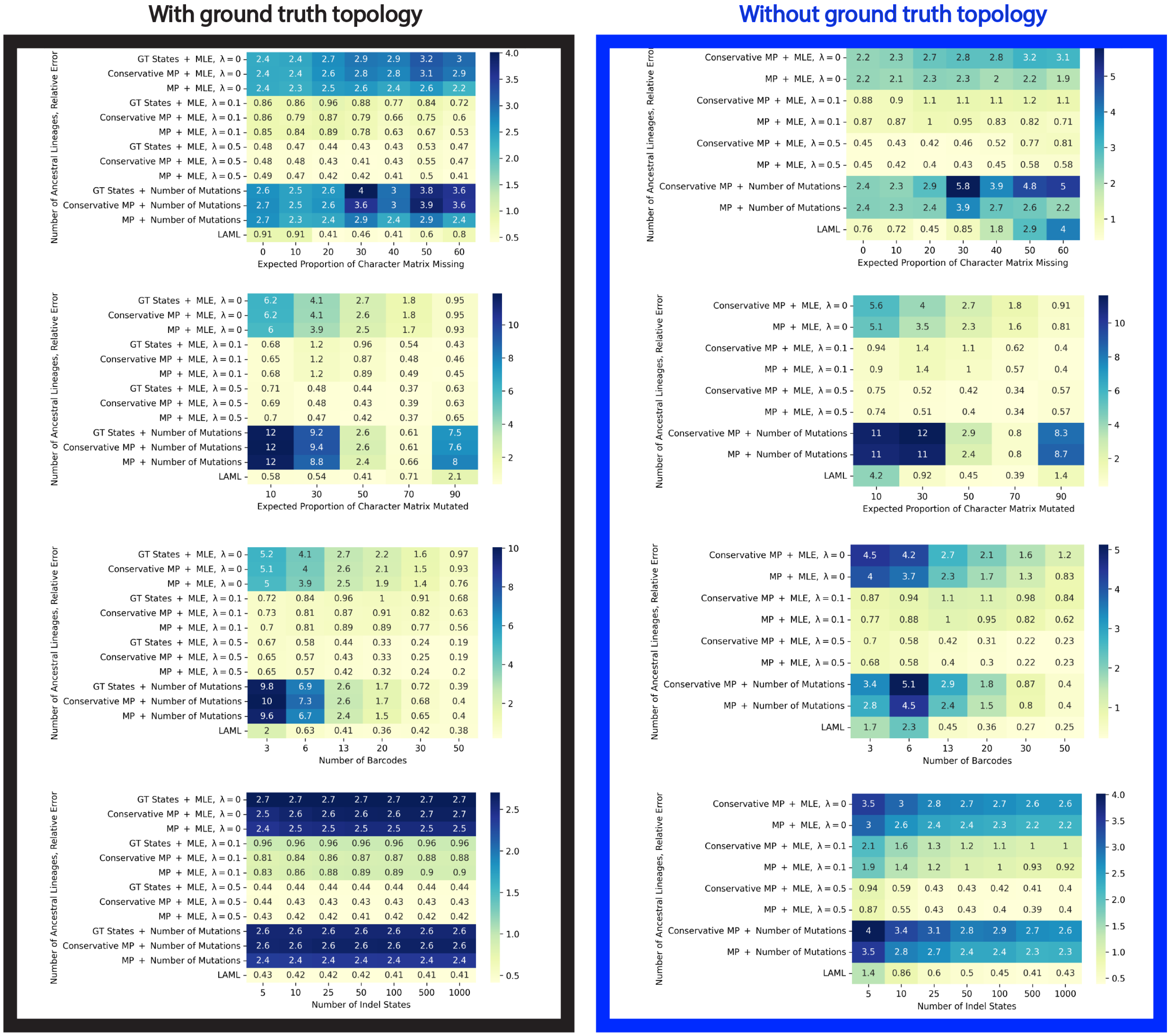


Supplementary Fig. 15. **Performance of branch length estimation methods under different parameter regimes for the “ancestral lineages” task, on larger trees with** 2000 **leaves**. On the left we show performance when the ground truth topology is known, and on the right we show the performance when the topology is not know and must be reconstructed – as is the case in all real-life applications. In this latter case, the Maxcut algorithm from the Cassiopeia package is used. Each number displayed is the average over the 50 simulated trees for the given parameter regime.

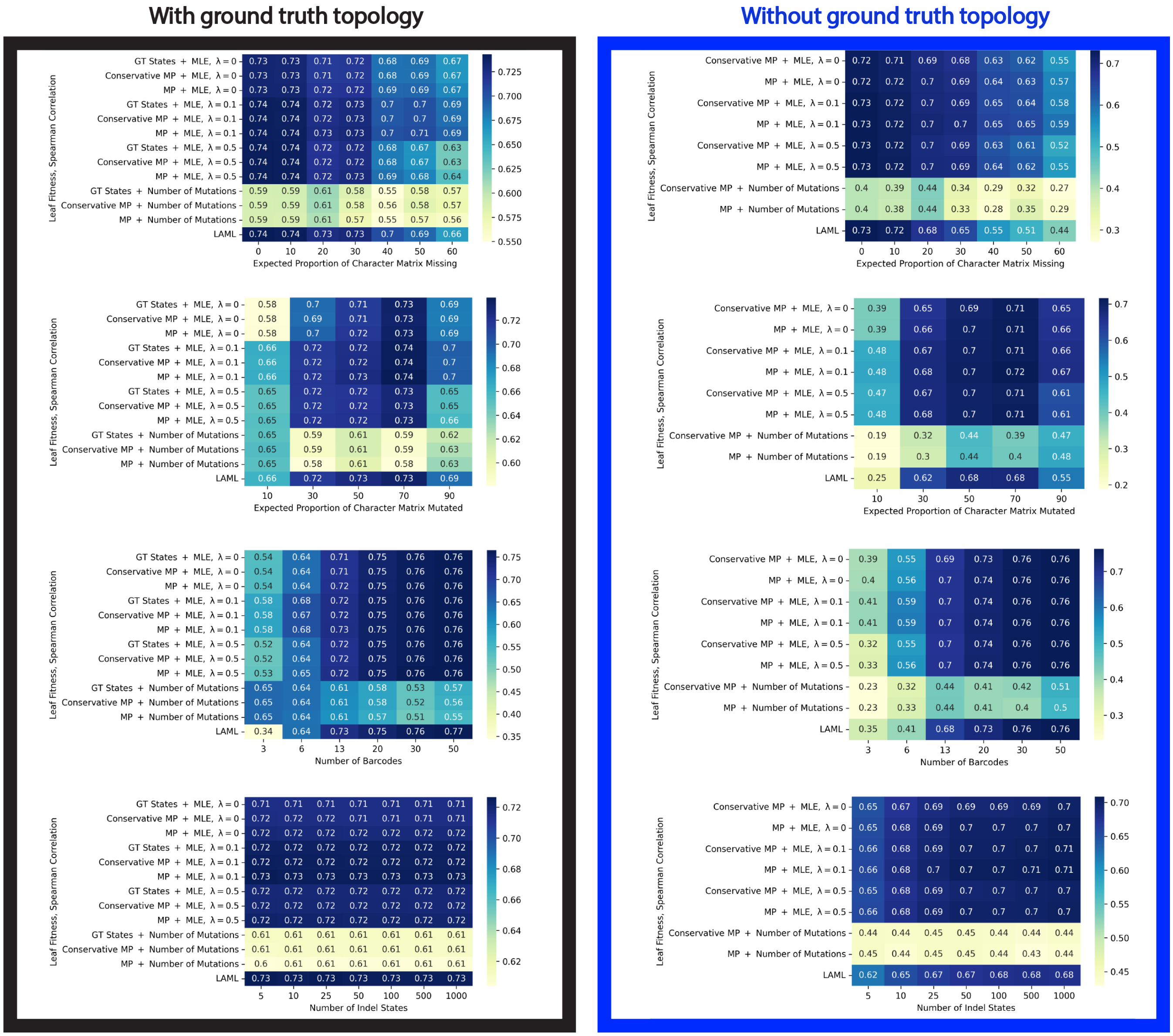


Supplementary Fig. 16. **Performance of branch length estimation methods under different parameter regimes for the "fitness inference" task, on larger trees with** 2000 **leaves**. On the left we show performance when the ground truth topology is known, and on the right we show the performance when the topology is not know and must be reconstructed – as is the case in all real-life applications. In this latter case, the Maxcut algorithm from the Cassiopeia package is used. Each number displayed is the average over the 50 simulated trees for the given parameter regime.

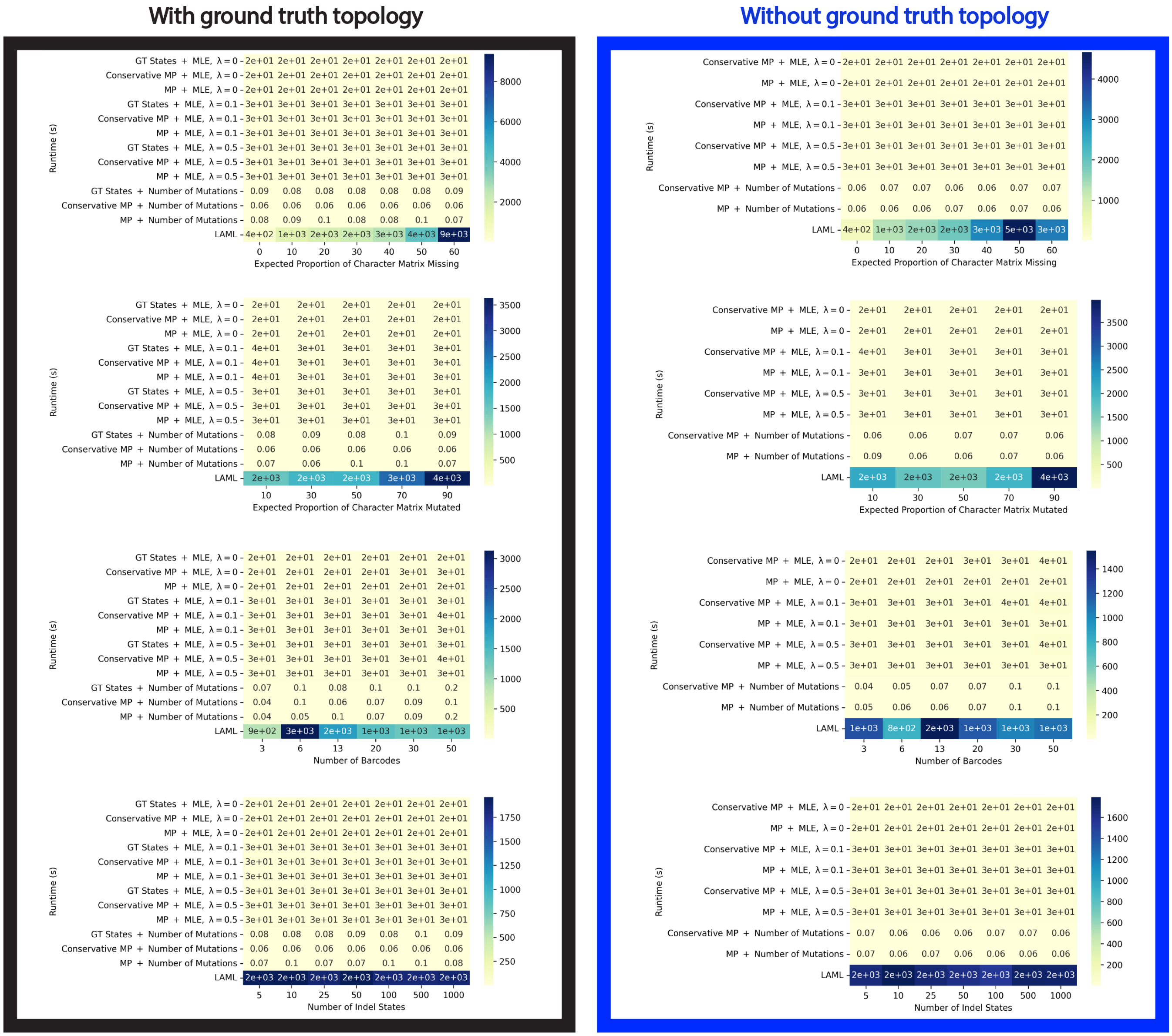


Supplementary Fig. 17. **Runtime of branch length estimation methods under different parameter regimes, on larger trees with** 2000 **leaves**. On the left we show performance when the ground truth topology is known, and on the right we show the performance when the topology is not know and must be reconstructed – as is the case in all real-life applications. In this latter case, the Maxcut algorithm from the Cassiopeia package is used. Each number displayed is the average over the 50 simulated trees for the given parameter regime. Runtime is shown in seconds, with one significant digit.

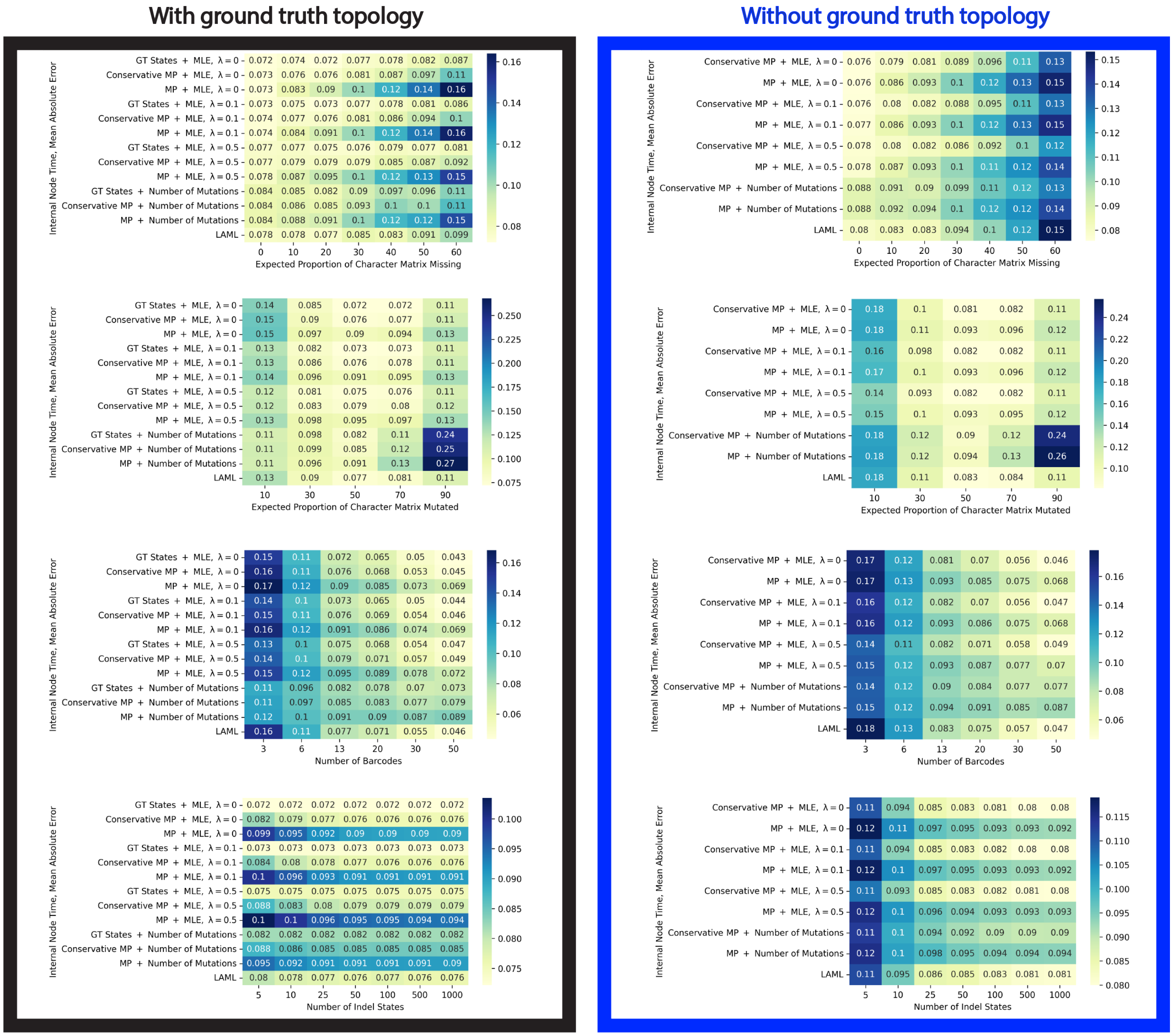


Supplementary Fig. 18. **Performance of branch length estimation methods under different parameter regimes for the "internal node time" task, on smaller trees with only** 40 **leaves**. On the left we show performance when the ground truth topology is known, and on the right we show the performance when the topology is not know and must be reconstructed – as is the case in all real-life applications. In this latter case, the Maxcut algorithm from the Cassiopeia package is used. Each number displayed is the average over the 50 simulated trees for the given parameter regime.

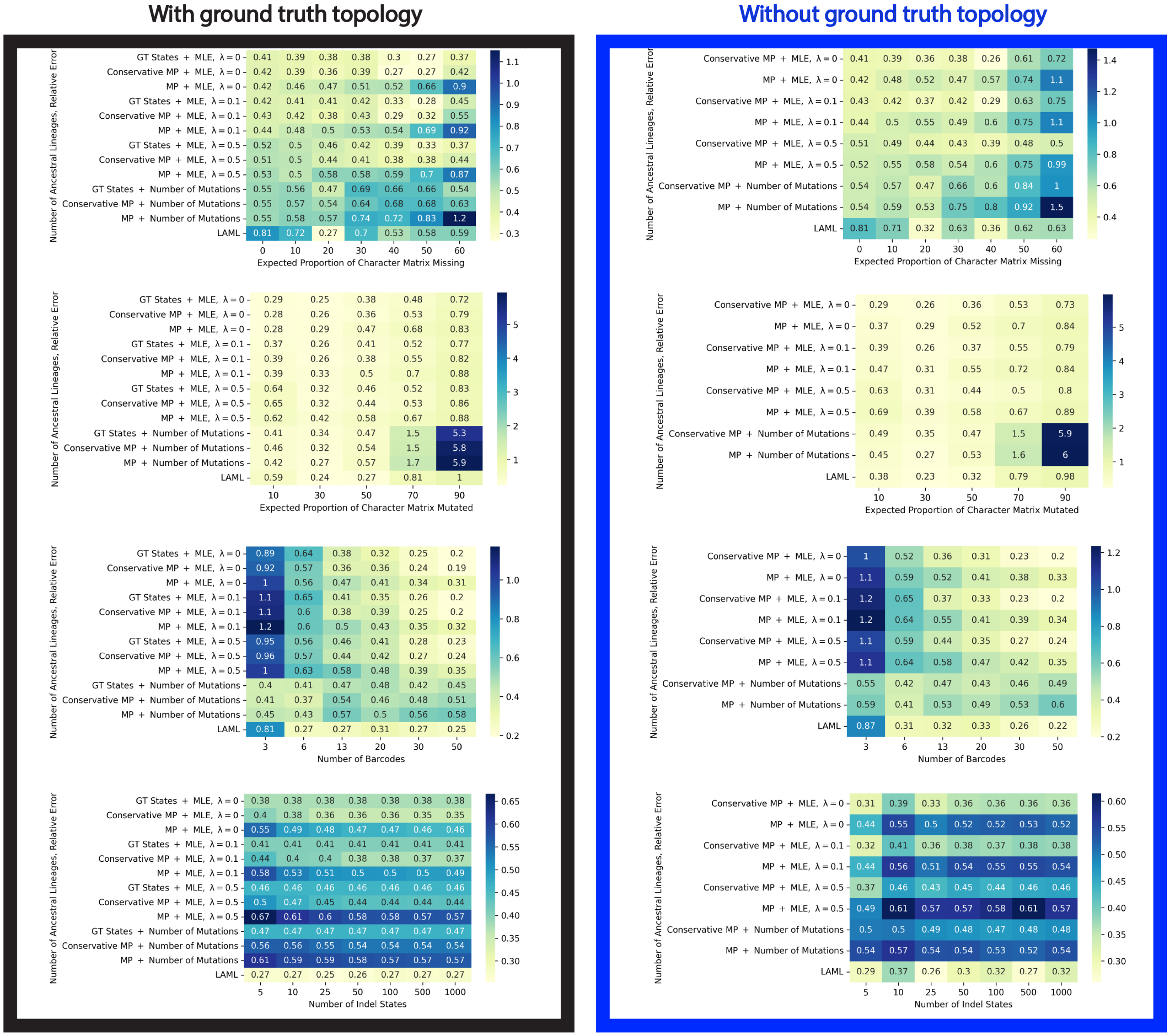


Supplementary Fig. 19. **Performance of branch length estimation methods under different parameter regimes for the "ancestral lineages" task, on smaller trees with only** 40 **leaves**. On the left we show performance when the ground truth topology is known, and on the right we show the performance when the topology is not know and must be reconstructed – as is the case in all real-life applications. In this latter case, the Maxcut algorithm from the Cassiopeia package is used. Each number displayed is the average over the 50 simulated trees for the given parameter regime.

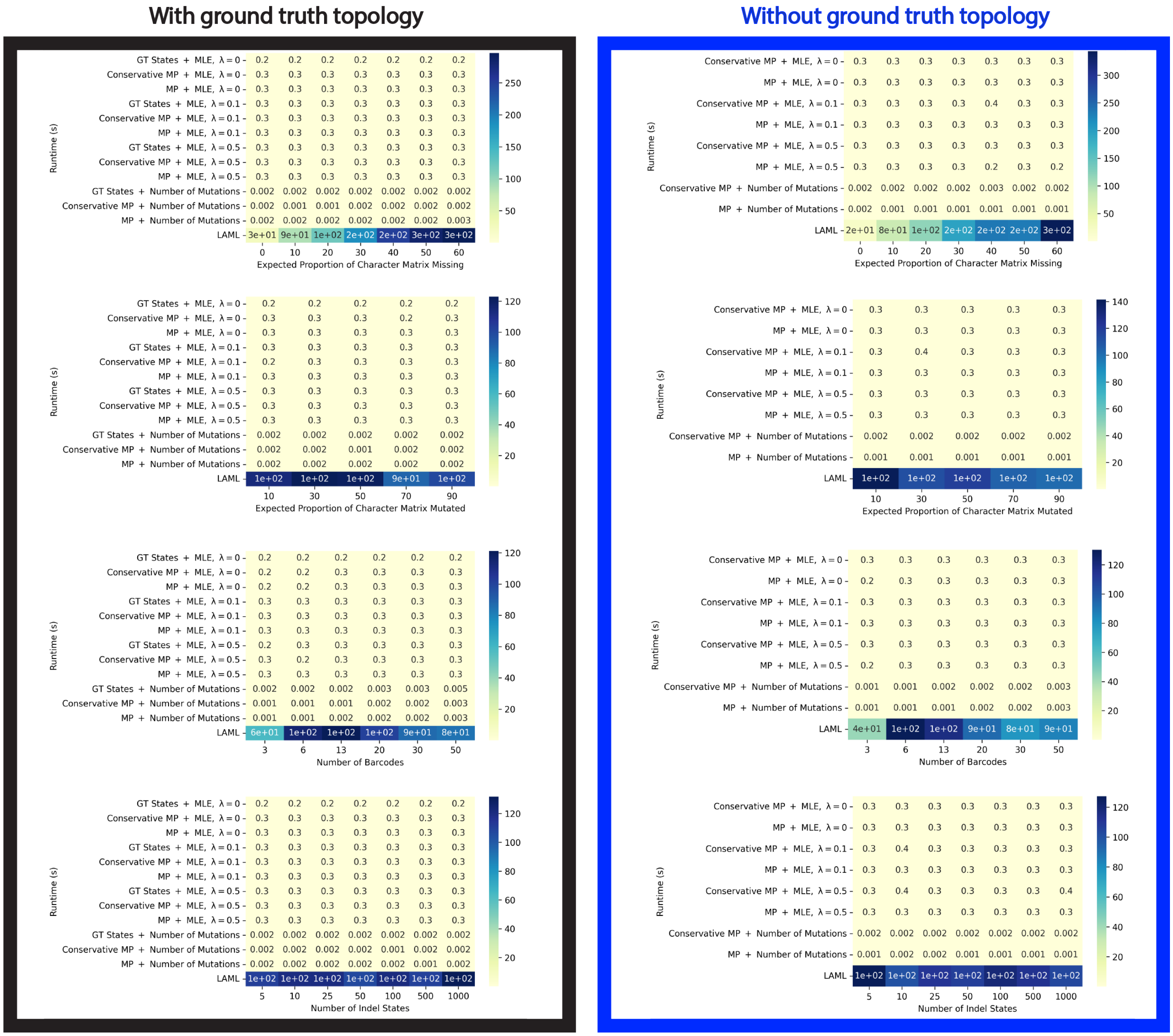


Supplementary Fig. 20. **Runtime of branch length estimation methods under different parameter regimes, on smaller trees with only** 40 **leaves**. On the left we show performance when the ground truth topology is known, and on the right we show the performance when the topology is not know and must be reconstructed – as is the case in all real-life applications. In this latter case, the Maxcut algorithm from the Cassiopeia package is used. Each number displayed is the average over the 50 simulated trees for the given parameter regime. Runtime is shown in seconds, with one significant digit.

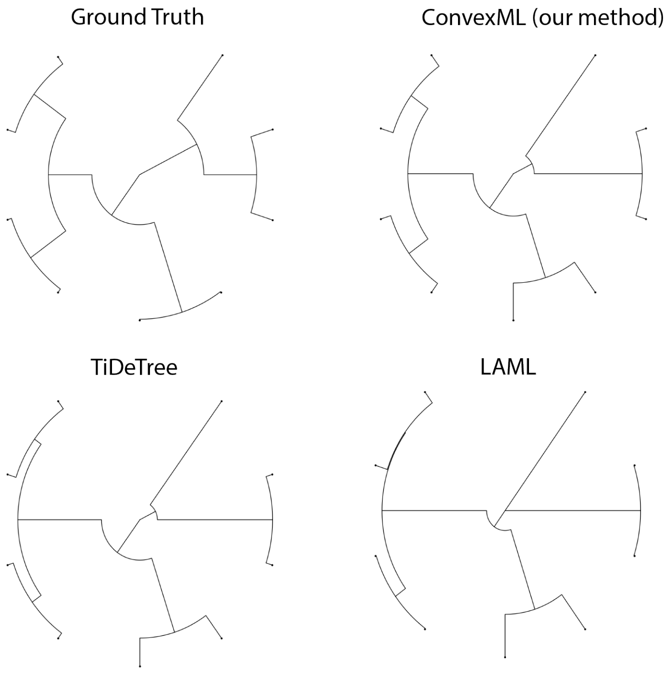


Supplementary Fig. 21. **Sample reconstructions on the intMEMOIR dataset**. Despite being an MLE method, ConvexML produces high-quality estimates which are quantitatively comparable to the more sophisticated Bayesian TiDeTree method. In contrast, despite marginalizing out ancestral states, the MLE method LAML produces degenerate branch length estimates akin to those predicted by our Theorem due to the lack of regularization.

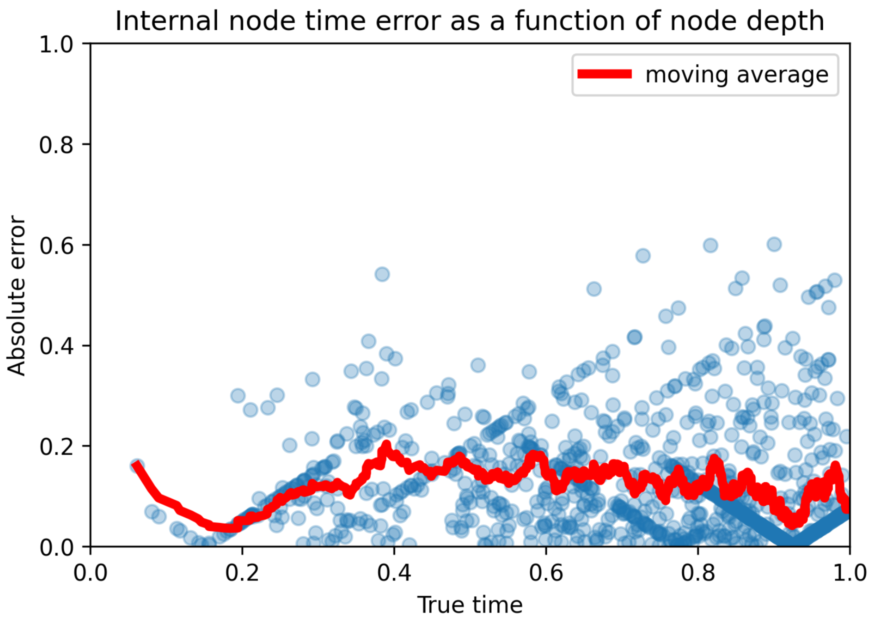


Supplementary Fig. 22. **ConvexML's error as a function of node depth**. We can see that the estimation error is stable across different node depths, showing that the mean absolute error metric is robust. Sets of point on a diagonal are the result of our minimum branch length constraint.
